# Cardiopharyngeal Mesoderm specification into cardiac and skeletal muscle lineages in gastruloids

**DOI:** 10.1101/2023.05.15.540476

**Authors:** Laurent Argiro, Céline Chevalier, Caroline Choquet, Nitya Nandkishore, Adeline Ghata, Anaïs Baudot, Stéphane Zaffran, Fabienne Lescroart

**Author notes:** These authors contributed equally to this work (Laurent Argiro, Céline Chevalier, Caroline Choquet). co-last authors (Stéphane Zaffran, Fabienne Lescroart).

## Abstract

Cardiopharyngeal mesoderm contributes to the formation of the heart and head muscles. However, the mechanisms governing cardiopharyngeal mesoderm specification remain unclear. Indeed, there is a lack of an *in vitro* model replicating the differentiation of both heart and head muscles to study these mechanisms. Such models are required to allow live-imaging and high throughput genetic and drug screening. Here, we show that the formation of self-organizing or pseudo-embryos from mouse embryonic stem cells (mESCs), also called gastruloids, reproduces cardiopharyngeal mesoderm specification towards cardiac and skeletal muscle lineages. By conducting a comprehensive temporal analysis of cardiopharyngeal mesoderm establishment and differentiation in gastruloids and comparing it to mouse embryos, we present the first evidence for skeletal myogenesis in gastruloids. By inferring lineage trajectories from the gastruloids single-cell transcriptomic data, we further suggest that heart and head muscles formed in gastruloids derive from cardiopharyngeal mesoderm progenitors. We identify different subpopulations of cardiomyocytes and skeletal muscles, which most likely correspond to different states of myogenesis with “head-like” and “trunk-like” skeletal myoblasts. These findings unveil the potential of mESC-derived gastruloids to undergo specification into both cardiac and skeletal muscle lineages, allowing the investigation of the mechanisms of cardiopharyngeal mesoderm differentiation in development and how this could be affected in congenital diseases.

## Introduction

Deployment of progenitors and their proper allocation to correct cell lineages are fundamental for the formation of organs. Defects in the specification and differentiation of progenitors into particular cell lineages lead to congenital anomalies. For example, a recent report showed improper cardiopharyngeal mesoderm progenitor development in a 22q11.2DS mouse model, where head muscle and heart morphogenesis were impaired ^1^.

### Cardiopharyngeal mesoderm includes bipotent progenitors for heart and head muscles

The heart forms from two cell populations, namely the first and second heart fields. The first heart field forms essentially the left ventricle with small contributions to the right ventricle and atria ^2, 3^. Second heart field progenitors, located in cardiopharyngeal mesoderm, contribute to cardiac muscles (myocardium) of the outflow tract, right ventricle and atria ^2–7^. Cardiopharyngeal mesoderm populates the pharyngeal arches and also gives rise to head and a subset of neck skeletal muscles, in addition to cardiac muscles ^8–11^. Clonal analyses in the mouse model have revealed the existence of bipotent progenitors in cardiopharyngeal mesoderm that form both head and heart muscles in vertebrates with different contributions along the antero-posterior axis of the pharyngeal region of the embryo ^8, 12, 13^. Bipotent progenitors were thus found for distinct subsets of myocardium and specific head and neck skeletal muscles ^12, 13^. Lineage tracing of early nascent mesoderm expressing the bHLH transcription factor Mesp1 showed that bipotent head and heart muscle progenitors are in fact present at the onset of gastrulation ^14^. Cardiopharyngeal mesoderm is evolutionary conserved since similar multipotent cardiopharyngeal progenitors have been shown to give rise to heart and pharyngeal muscles in tunicates ^9, 15^.

### Cardiopharyngeal mesoderm contributes to skeletal muscles of the head and neck with a particular genetic program

For its skeletal muscles derivatives, cardiopharyngeal mesoderm contributes to specific head/neck muscles ^9, 10^. Skeletal muscles derived from cardiopharyngeal mesoderm have distinct genetic regulatory programs compare to trunk and limb skeletal muscles ^16^. Trunk and limb skeletal muscles derive from somitic paraxial mesoderm. Somitic myogenesis is regulated by Pax3 (a homeobox paired-domain transcription factor), which regulates the myogenic regulatory transcription factors MyoD and Myf5. Strikingly, *Pax3/Myf5* mutant embryos failed to develop trunk and limb muscles while head muscles were broadly not affected, thus decoupling trunk (somitic) from head (cardiopharyngeal) myogenesis ^17^. In cardiopharyngeal mesoderm, the myogenic factors MyoD/Myf5 are regulated by a different core of transcription factors that together mark the cardiopharyngeal mesoderm but are individually not strictly restricted to this population. They include Tbx1, Lhx2, MyoR (also called Msc), Tcf21 (also called Capsulin) and Pitx2 ^18–22^. Myogenin (MyoG) then marks the start of myoblast differentiation. The next steps of muscle differentiation are common between cardiopharyngeal and somitic mesoderm derived progenitors ^16^.

### Insights on how cardiopharyngeal mesoderm specification occurs mostly arise from studies in tunicates

Major insights into the specification of multipotent cardiopharyngeal mesoderm progenitors and the mechanisms of cardiopharyngeal lineage segregation have come from studies in Ciona. In this tunicate model, Tbx1/10 activates COE/Ebf for the specification of the muscular lineage ^23, 24^. It is however not completely clear how comparable are tunicates and vertebrates. The heart and pharyngeal muscles derive from only 2 Mesp+ progenitors in tunicates, while about 250 Mesp1+ progenitors contribute to heart development in mouse ^9, 25^. Despite the number of important studies using lineage tracing and clonal analysis, it is challenging to analyze cardiopharyngeal mesoderm specification in vertebrates due to its difficult accessibility and the absence of restricted and specific markers. There is a need of a simple model that could faithfully recapitulate vertebrate, and more specifically mammalian, cardiopharyngeal mesoderm development and could allow live-imaging and large throughput analyses.

### Pluripotent stem cells-derived models to study Cardiopharyngeal mesoderm

Pluripotent stem cells have emerged as an interesting tool in general to model how transcription factors and signaling molecules interact to control cell fate decisions and lineage specification ^26, 27^. Using mouse embryonic stem cells (mESCs), Chan et al. have shown that Mesp1+-derived progenitors have a dual heart and skeletal muscle differentiation potential, when cultured in a pro-cardiogenic or a pro-skeletal myogenic culture medium ^28^. Mouse and human ESCs have also been used to investigate the differences in molecular cues involved in myogenic specification between cardiopharyngeal versus somitic mesoderm ^29^. Both studies however, required changes in cell culture medium and growth factors to promote either cardiac or skeletal muscle differentiation ^28, 29^. Therefore, there is a lack of an *in vitro* model that allows parallel differentiation of cardiac and skeletal muscle tissue and faithfully recapitulates *in vivo* cardiopharyngeal mesoderm early development.

### Gastruloids model early cardiac development

The establishment of gastruloids provides an alternative model for cardiopharyngeal mesoderm development. The model of gastruloids was first described in 2014 with axial elongation and symmetry breaking from the aggregation of a restricted number of mESCs cultured with a pulse of Wnt activation (with Chiron treatment) ^30, 31^. This model faithfully recapitulates early mouse development mimicking *in vivo* gene expression ^32^. Recently, the model has been pushed further towards early organogenesis, with development of heart-like structures with first and second heart field components ^33, 34^ or somites ^35, 36^. Despite the formation of somites, skeletal myogenesis, either cardiopharyngeal or somitic, has not yet been demonstrated in gastruloids.

Here, we have adapted the self-organizing mESCs-based gastruloid protocol for longer times of culture to investigate whether gastruloids can model cardiopharyngeal mesoderm specification and differentiation. Multiplex fluorescent *in situ* hybridization shows that cardiopharyngeal mesoderm specification occurs in gastruloids in a similar spatio-temporal organization as observed in mouse embryos. Using single-cell RNAseq analysis along a time-course from day 4 to late day 11 of culture, we demonstrate the presence of three subpopulations of cardiomyocytes and two subpopulations of myoblasts. We find that gastruloids can undergo myogenesis and that gastruloid-derived myoblasts can arise from both cardiopharyngeal and somitic paraxial mesoderm. Therefore, gastruloids can be used to reconstruct cardiopharyngeal mesoderm development.

## Results

### Cardiopharyngeal mesoderm markers are expressed in gastruloids

To generate gastruloids that form cardiopharyngeal mesoderm and characterize their differentiation to cardiac and skeletal myogenic fates, we optimized previous protocol described by Rossi et al. ^34^ to culture gastruloids for an extended time, until at least day 11 (Fig. 1a) (see Methods). In brief, mESCs were aggregated at day 0, following centrifugation, and treated with a Wnt agonist (Chiron) for 24h from day 2. Cardiogenic factors (bFGF, VEGF and ascorbic acid) were added to the culture media at 96 h (day 4) for 3 days. We noted elongation from day 4 and beating regions appearing at day 7, on the anterior part of the gastruloid (Fig. 1b.).

**Figure 1:**
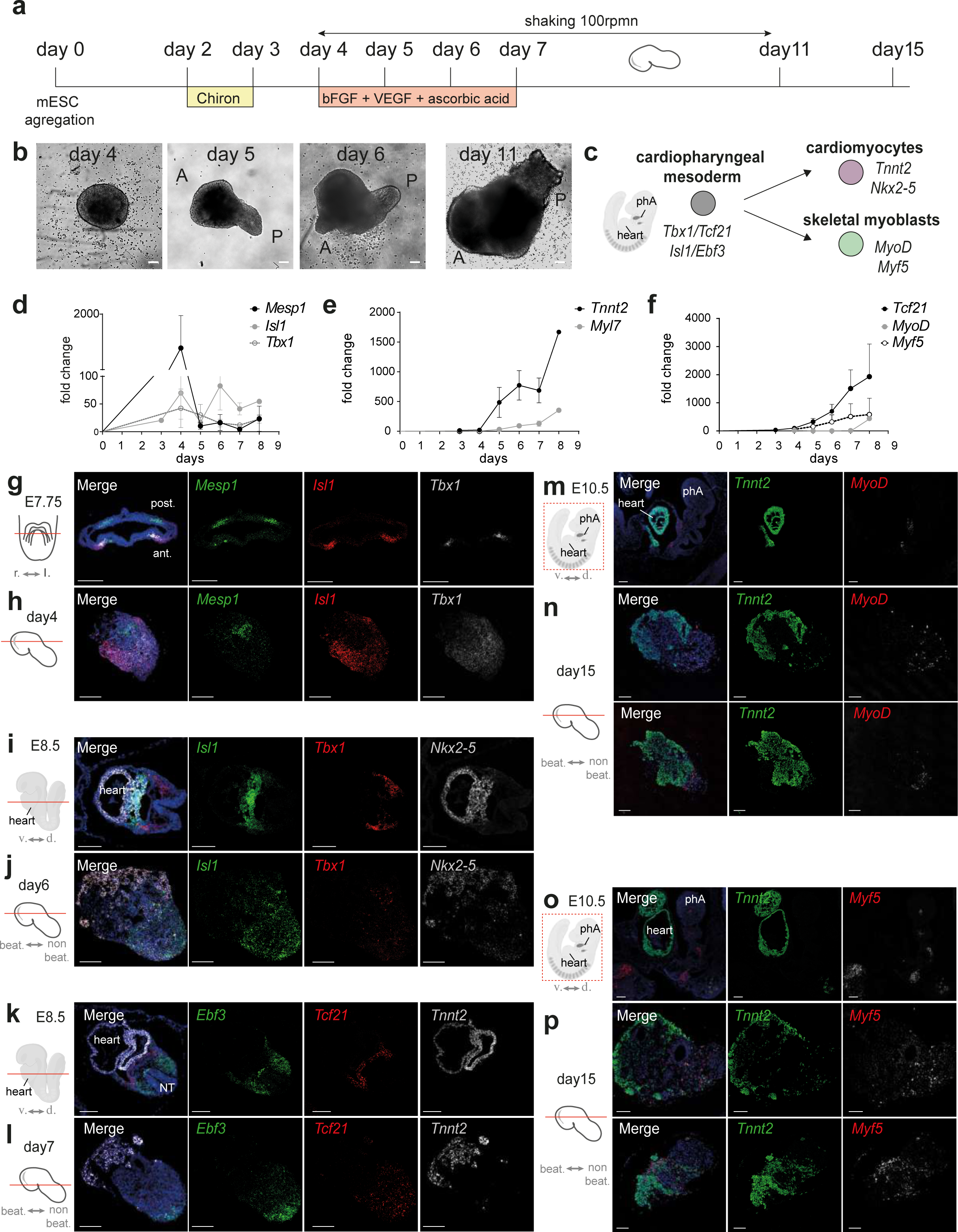
Gastruloids faithfully recapitulates cardiopharyngeal mesoderm specification and differentiation toward its myogenic fates. **a.** Experimental scheme of the generation of gastruloids from mouse embryonic stem cells (mESCs). Gastruloids were cultured in N2B27 with the addition of small molecules as indicated below the time line. **b.** Representative pictures of gastruloids at day 4, day 5, day 6 and day 11. A, anterior, P, posterior. **c.** Scheme of cardiopharyngeal mesoderm specification in mouse toward the myogenic fates. Markers of each state are indicated in italics. **d-f.** Expression profiles of *Mesp1*, *Isl1* and *Tbx1* (**d**), *Tnnt2* and *Myl7* (**e**) and *Tcf21*, *MyoD* and *Myf5* (**f**) throughout the culture of gastruloids as measured by quantitative RT-PCR. Results are normalized on the expression of *Tbp*. Fold changes are represented relative to expression on day 0. (mean with standard error of mean (SEM)). **g-h.** Representative confocal images of sections across an E7.75 mouse embryo (**g**) and a gastruloid at day 4 (**h**) after RNAscope experiment with *Mesp1* (green), *Isl1* (red) and *Tbx1* (white) probes. **i-j.** Representative confocal images of sections across an E8.5 mouse embryo (**i**) and a gastruloid at day 6 (**j**) after RNAscope experiment with *Isl1* (green), *Tbx1* (red) and *Nkx2-5* (white) probes. **k-l.** Representative confocal images of sections across an E8.5 mouse embryo (**k**) and a gastruloid at day 7 (**l**) after RNAscope experiment with *Ebf3* (green), *Tcf21* (red) and *Tnnt2* (white) probes. **m-p.** Representative confocal images of sections across E10.5 mouse embryos (**m** and **o**) and gastruloids at day 15 (**n** and **p**) after RNAscope experiment with *Tnnt2* (green) and MyoD (**m-n**) or *Myf5* (**o-p**). r, right; l, left. v, ventral; d, dorsal; beat., beating; non beat., non-beating., ant., anterior, post., posterior, phA, pharyngeal arch. NT, neural tube. Scale bars: 100 µm.

To investigate whether key markers of cardiopharyngeal mesoderm and its derivatives (Fig. 1c) were expressed during the time-course culture, we used quantitative RT-PCR. We observed the transient expression of the bHLH transcription factor *Mesp1* and the expression of the cardiopharyngeal mesoderm transcription factors *Islet1* (*Isl1*) and *Tbx1*, as previously described ^34^ (Fig. 1d). We also observed an increasing expression of transcripts encoding the cardiac specific myosin (*Myl7*) and troponin (*Tnnt2*) at day 5 (Fig. 1e). Interestingly, the cardiopharyngeal marker, *Tcf21* ^20, 24^, began to be expressed at day 4. The myogenic transcription factors *Myf5* and *MyoD* were expressed at day 6 and day 8 respectively (Fig. 1f). Importantly, similar kinetics of expression were also observed for *Mesp1*, *Isl1* and *Tcf21* with another wild-type mESC line (see Methods for details on the lines and Supplementary Fig. S1). These data demonstrate that cardiopharyngeal mesoderm and downstream myogenic transcriptional programs are robustly activated in gastruloids under these culture conditions.

### Comparison with mouse embryo shows similar spatio-temporal gene expression

To explore whether gastruloids faithfully mimic mouse cardiopharyngeal mesoderm development, we compared gastruloids with mouse embryos at equivalent developmental stages. Our goal was to investigate the expression of markers of the cardiopharyngeal mesoderm, cardiomyocytes, and the robustness of the cardiopharyngeal mesoderm specification. We used multiplex fluorescent *in situ* hybridization or RNAscope to explore gene expression. Comparison of day 4 gastruloids with E7.75 cardiac crescent embryos showed expression of *Mesp1* in non-overlapping domains from the expression of *Isl1* and *Tbx1* (Fig. 1g-h). In the embryo, *Mesp1* was observed in the posterior mesoderm (likely corresponding to the somitic mesoderm), while *Tbx1* and *Isl1* expression overlapped in the anterior region of the embryo (Fig. 1g). Similarly, *Tbx1* and *Isl1* expression overlapped in gastruloids (Fig. 1h). We then investigated *Tbx1, Isl1*, and *Nkx2*-5 expression at later time points in E8.5 embryos and day 6 gastruloids (Fig. 1i-j). In the embryo, *Nkx2-5* expression labelled the heart tube while *Isl1* and *Tbx1* were expressed in second heart field progenitors, located in pharyngeal mesoderm behind the differentiated heart tube (Fig. 1i). *Nkx2-5* was also expressed in the anterior part of the gastruloid, marking cardiomyocytes. In day 6 gastruloids, *Tbx1* and *Isl1* were expressed in a domain adjacent to that of *Nkx2-5* (Fig. 1j). Similarly, we found that *Tcf21* and *Ebf3* were expressed in pharyngeal mesoderm of E8.5 embryos behind the heart tube expressing *Tnnt2* (Fig. 1k). In day 7 gastruloids, *Ebf3* and *Tcf21* were also found in a close domain of expression adjacent to the cardiomyocytes (*Tnnt2*+) (Fig. 1l). Taken together these findings show that cardiopharyngeal mesoderm markers are expressed in a similar spatio-temporal pattern in both gastruloids and mouse embryos.

To further investigate whether myoblast differentiation occurs and is faithfully recapitulated in gastruloids, we also compared the expression of myoblast markers in both models. At E10.5, *Myf5* and *MyoD* positive cells were detected in the pharyngeal arches of mouse embryos, close to the heart (Fig. 1m and o). Similarly, *Myf5* and *MyoD* were detected close to cardiomyocytes (*Tnnt2*-positive cells) in gastruloids collected at day 15 (Fig. 1m-p). We noted however that *Myf5* and *MyoD* did not show a compact expression as observed in the embryo indicating the absence of arch morphogenesis. These data support the existence of skeletal myogenesis in gastruloids cultured for more than 10 days. Together, this comparison shows that gastruloids and mouse embryos display similar spatio-temporal gene expression during cardiopharyngeal mesoderm development.

### Single-cell RNA-sequencing reveals cardiopharyngeal mesoderm subpopulations in gastruloids

To further assess the potential of gastruloids to form cardiopharyngeal mesoderm and investigate in detail the different cell populations, we performed single-cell RNA-sequencing (scRNA-seq) on gastruloids following a time course. Gastruloid cells were collected at day 4, day 5, day 6 and day 11. For each time point, at least 8 gastruloids were dissociated into single-cell suspensions and approximately 7,000 cells were sequenced. After quality controls, we analyzed 6,646 cells at day 4, 6,704 cells at day 5, 5,284 cells at day 6 and 8,024 cells at day 11. We performed Leiden clustering and differential gene expression analysis at each time point (Fig. 2 and Supplementary Table S1). Integration of the gastruloid single-cell data with the published atlas of mouse embryonic cells ranging from embryonic day (E) 6.5 to E8.5 ^37^ (see Methods 4-5) showed that cells collected at day 4, day 5 and day 6 likely overlap with cells of E7.25-E7.75, E8.0-E8.25 and E8.5 embryos respectively (Supplementary Fig. S2), in agreement with our fluorescent *in situ* hybridization experiments (Fig. 1g-p). However, gastruloid cells collected at day 11 do not overlap in the UMAP with cells of the atlas, maybe reflecting distinct transcriptional profiles (Supplementary Fig. S2j-k). Gastruloids cells at day 11 might thus reflect more differentiated states, usually found after E8.5 in mouse embryos. This observation indicates that while we can confidently compare our gastruloid single-cell dataset at day 4, day 5 and day 6 with the cells of the atlas, distinct annotation criteria had to be applied to day 11 clusters. Cell type annotation transfer, from the atlas, was first applied to the primary clusters of day 4, 5 and 6 (See Methods 4). Manual annotation was then performed based on this label transfer and on differential gene expression (Supplementary Table S1). The clusters of day 11 were only annotated manually based on differential gene expression analysis (Supplementary Table S1) and tissue database (https://tissues.jensenlab.org/About).

**Figure 2:**
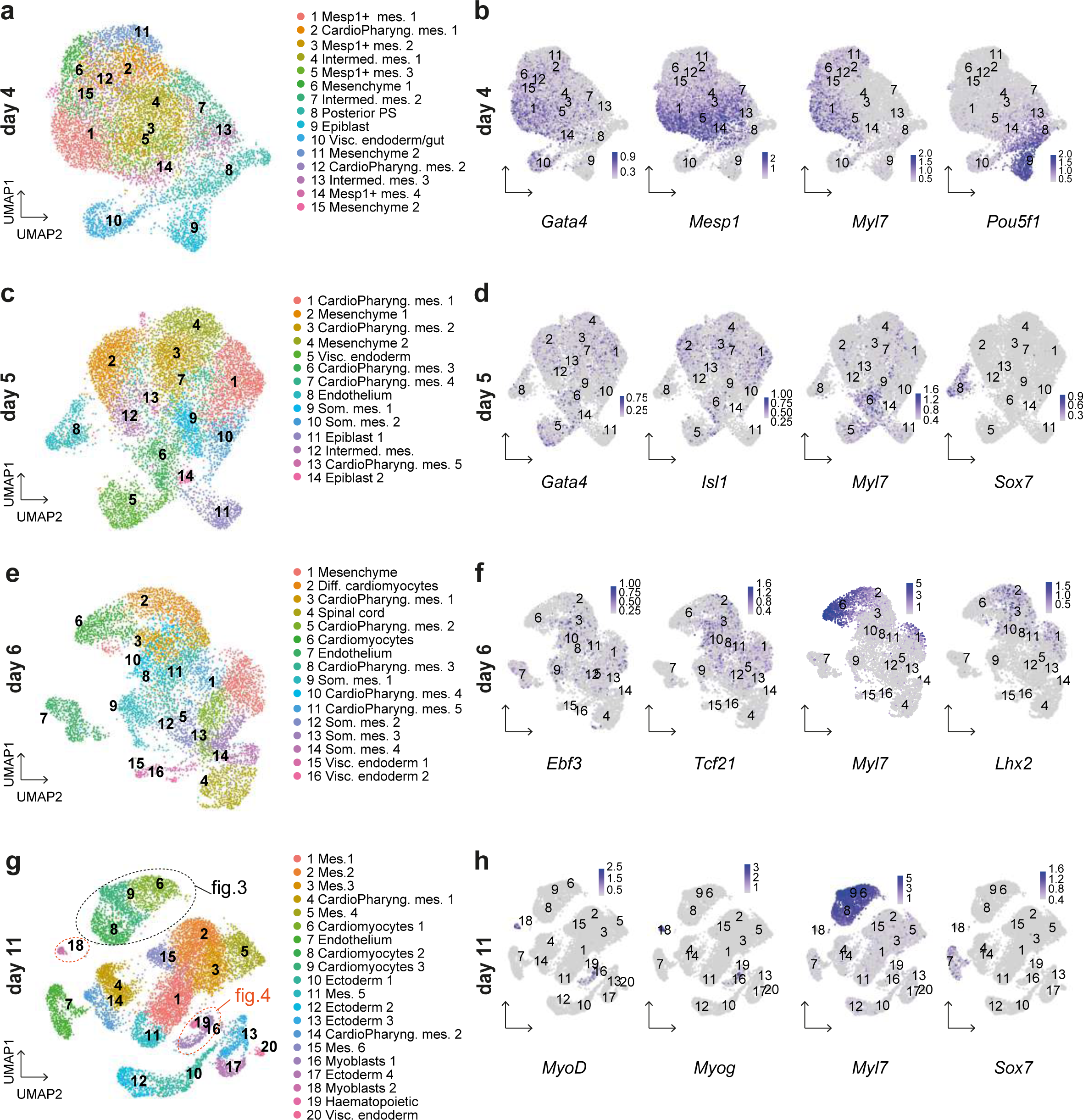
Transcriptomic analysis reveals cardiopharyngeal mesoderm subpopulations in gastruloids. UMAP representation of Leiden clustering of gastruloids at day 4 (**a**), day 5 (**c**), day 6 (**e**) and day 11 (**g**). FeaturePlots of key markers in gastruloids at day 4 (**b**), day 5 (**d**), day 6 (**f**) and day 11 (**h**). Scale bars represent expression levels. Mes., mesoderm; CardioPharyng., cardiopharyngeal; Intermed., intermediate; Visc., visceral; Som. Somitic; Diff., differentiating; PS, primitive streak.

To assess the cellular heterogeneity of the gastruloids, we analyzed the different clusters obtained at each time point. At day 4, we observed a significant number of clusters of mesodermal cells (Fig. 2a-b; Supplementary Fig. S3) with four clusters showing high *Mesp1* expression (clusters 1, 3, 5 and 14). Analysis of the single-cell data also showed high expression of the *Gata4* and *Gata6* transcription factors in most clusters (all clusters except clusters 8, 9 and 13) (Fig. 2b; Supplementary Fig. S4). These transcription factors are known to be involved in cardiopharyngeal mesoderm specification ^38, 39^. *Hand1*, a marker of the extraembryonic mesoderm and cardiac progenitor cells in the recently identified juxtacardiac field ^40, 41^ was also highly expressed at day 4 (except in clusters 8, 9 and 10). Interestingly, with scRNA-seq, we detected *Myl7* as early as day 4. Cluster 9 showed expression of the pluripotency marker *Pou5f1* indicative of epiblast-like cells (Fig. 2b; Supplementary Fig. S3). *Mesp2*, the closest homolog of *Mesp1,* was only expressed at very low levels in gastruloids (Supplementary Fig. S4). At day 5, the cell type annotation transfer from the atlas was enriched in mesenchymal and pharyngeal mesodermal cells while nascent mesoderm was barely observed (Supplementary Fig. S5). We identified 5 clusters annotated as pharyngeal mesoderm (clusters 1, 3, 6, 7, and 13) (Fig. 2c). The first fully differentiated cells were found in cluster 8. This cluster corresponds to endothelial cells and is marked by the expression of *Sox7* (Fig. 2c-d Supplementary Fig.S5). At this stage, *Mesp1* started to be downregulated while *Gata6* was still highly expressed (Supplementary Fig. S6). Expression of *Isl1* is also detected and *Tbx1* at a very low level (Fig. 2d, Supplementary Fig. S6). At day 6, we uncovered 16 clusters (Fig. 2e, Supplementary Fig. S7). We detected two clusters of differentiating or differentiated cardiomyocytes (clusters 6 and 2) expressing high level of *Myl7* and other cardiac myosin genes. All these data are consistent with the previous report on gastruloids ^34^. We also found 5 clusters of pharyngeal mesodermal cells (clusters 3, 5, 8, 10 and 11). Expression of the transcription factors involved in cardiopharyngeal mesoderm development, including *Tbx1*, *Tcf21*, *Lhx2* and *Ebf3,* was also detected (Fig. 2f and Supplementary Fig.S8). These data reveal the emergence of cardiopharyngeal mesoderm in differentiating gastruloids.

Finally, to investigate the potential differentiation of cardiopharyngeal mesoderm derivatives in gastruloids, we analyzed scRNA-seq data at day 11. We identified 20 clusters (Fig. 2g). One cluster (cluster 20) likely corresponds to visceral endoderm tissue, with expression of *Epcam*, *Sox17*. We also uncovered 4 different clusters (clusters 10, 12, 13 and 17) of ectodermal cells with enrichment of genes linked with neural derivatives, including neural crest-like cells (Supplementary Tables S1). We identified a cluster of endothelial cells, with expression of *Sox7* (cluster 7) (Fig. 2h) and a cluster of blood/hematopoietic lineages (cluster 19). Clusters 1, 2, 3, 4, 5, 11 and 14 were annotated as mesoderm and included mesenchymal cells and undifferentiated progenitors (Fig. 2g, Tables S1). The presence of high expression levels of *Myl7* in clusters 6, 8 and 9 indicates that these clusters contain cardiomyocytes. Strikingly, we found 2 clusters (clusters 16 and 18) with expression of the myogenic regulatory genes *MyoD* and *Myogenin* (*Myog*) (Fig. 2g-h). Together, our data thus reveal different cardiopharyngeal mesoderm subpopulations in the gastruloid model and show that cardiomyocytes and skeletal myoblasts markers are found in this model.

### Three subtypes of cardiomyocytes differentiate in gastruloids

In order to investigate whether the cardiomyocyte clusters at day 11 represent different cardiac subpopulations, we performed a detailed analysis of the clusters 6, 8 and 9, i.e., the three clusters containing cardiomyocytes. Among the genes differentially expressed in cluster 6, we found *Actg1*, *Cald1*, *Actb*, *Acta2*, *Vsnl1*, *Gucy1a1*, *Ptma*, *Map1b*, *Shox2* and *Ptn* (Fig. 3a-b and Supplementary Table S2). This cluster thus showed the expression of *Cald1*, *Acta2,* expressed in smooth-muscle cells and throughout the myocardium of the immature heart tube ^42, 43^ as well as genes involved in sinus venosus/sinus atrial node development (*Vsn1*, *Shox2*, *Tbx3*) ^44–46^ (Fig. 3a-b). In cluster 9, *Nppa*, *Itga6*, *Myl7*, *Myh6* were among the genes differentially expressed (Fig. 3c-d and Supplementary Table S1). These genes are usually enriched in atrial cardiomyocytes. Cluster 8 on the other hand was enriched in genes such as *Myl2*, *Myl3*, *Myh7*, *Mpped2*, *Pln* (Fig. 3e-f and Supplementary Table S1). These genes were previously shown to be the signature of ventricular cardiomyocytes in different independent studies ^47–49^. ScRNA-seq thus identified 3 cardiomyocyte clusters with distinct transcriptional signatures. We can further speculate that gastruloids contain ventricular, atrial and conductive-like cardiomyocytes.

**Figure 3:**
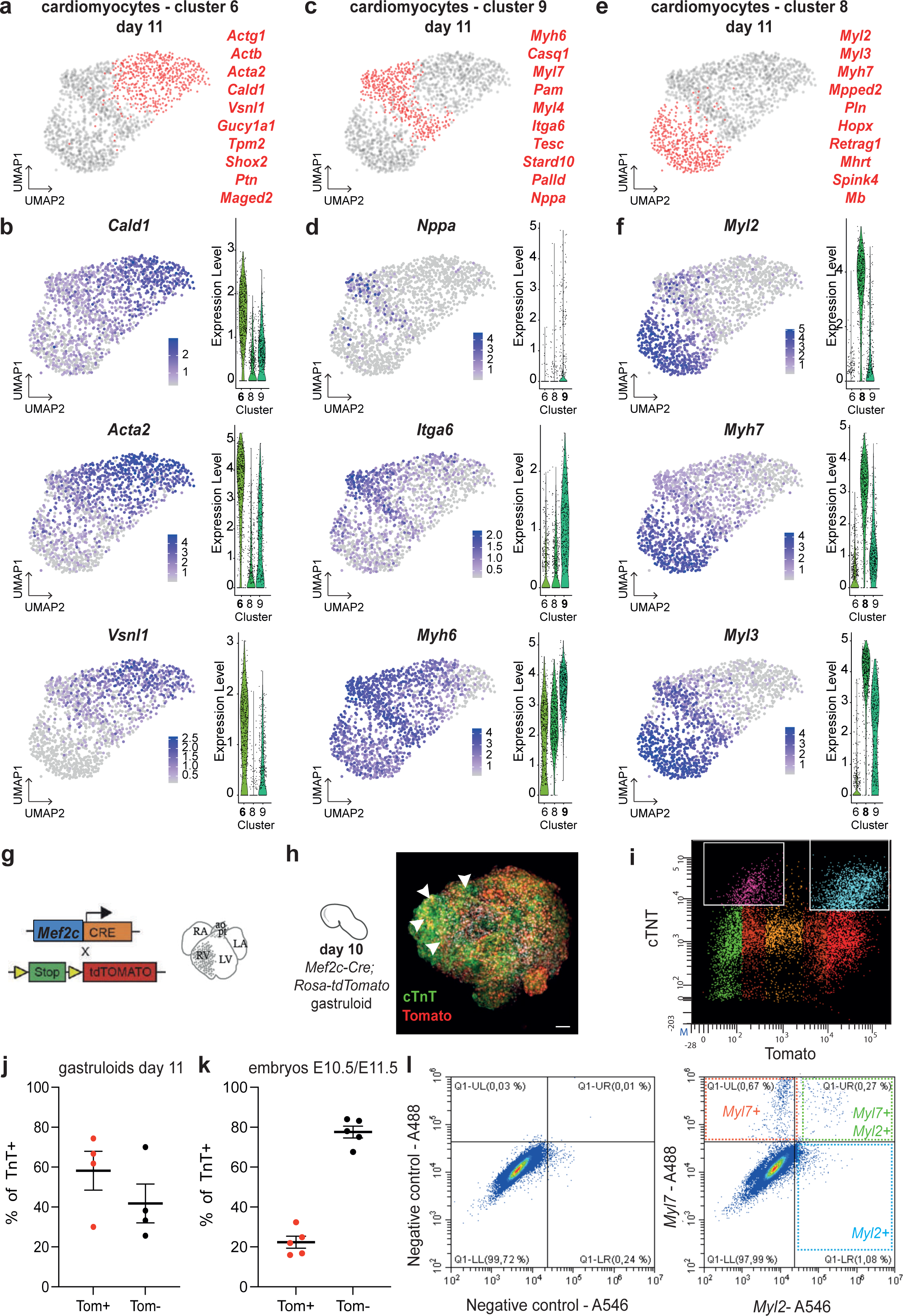
Three subtypes of cardiomyocytes differentiate in gastruloids. **a, c, e.** UMAP representation of day 11 cardiomyocytes. Red dots represent cells of cluster 6 (**a**), cluster 9 (**c**) or cluster 8 (**e**). The most differentially expressed genes of the cluster are shown in red. **b, d, f.** Feature and violin plots of key differentially expressed genes in the cardiomyocyte cluster 6 (**b**), cluster 9 (**d**) and cluster 8 (**f**) at day 11. **g.** Scheme of the experimental design with the use of the *Mef2c-AHF-enhaner-Cre; Rosa-tdTomato* (*Mef2c-Cre; Rosa-tdTomato*) line that label the right ventricle (RV) and outflow tract. La, left atrium; RA, right atrium; LV, left ventricle; ao, aorta; pt, pulmonary trunk. **h.** Confocal image (maximum intensity projection) of a *Mef2c-Cre; Rosa-tdTomato* gastruloid at day 10 after immunostaining with a marker of cardiomyocytes (cardiac troponin T – cTnT in green). tdTomato positive cells and nuclei are depicted in red and white, respectively. Arrowheads indicate cTnt+ tdTomato+ cells. Scale bar: 100 µm. **i.** FACS analysis of the combined expression of tdTomato and cTnT expression in all of the living gastruloids’ cells at day 11. **j-k.** Graphs showing the proportion of tdTomato+ (Tom+) cells in cardiomyocytes (cTnT+ cells) derived from day 11 gastruloids (**j**) and embryonic cells at E10.5/E11.5 (**k**) (in at least 4 independent experiment (mean with standard error of mean (SEM)). **l.** FACS analysis of the combined expression of *Myl7* and *Myl2* expression in all of the living gastruloids’ cells at day 11 after HCR experiment. Negative controls with no probe are shown in left.

In order to validate these results, we performed lineage tracing with a cell line that labels specific subpopulations of cardiomyocytes in gastruloids. For that purpose, we rederived mESCs from *Mef2c-AHF-enhancer-Cre; Rosa^tdTomato/+^*mouse blastocysts. *Mef2c-AHF-enhancer-Cre* mouse line is known to specifically label progenitors that contribute to outflow tract, right ventricle and a subpopulation of venous pole myocardium (dorsal mesenchymal protrusion) as well as non-cardiac cardiopharyngeal mesoderm derivatives ^50–52^ (Fig. 3g). *Mef2c-AHF-enhancer-Cre; Rosa^tdTomato/+^* (*Mef2c-Cre; Rosa-tdTomato*) derived mESCs were able to form beating gastruloids with expression of tdTomato in their anterior domain as expected, but without notable spatial organization (Fig. 3h). Flow cytometry at day 11 revealed that *Mef2c-Cre; Rosa-tdTomato*-derived gastruloids contain a significant proportion of cardiomyocytes, as shown by the expression of cardiac troponin T (cTnT) (Fig. 3i). Thus, tdTomato+ cardiomyocytes represented an average of 58.2% (ranging from 30 to 74.4%) of all cardiomyocytes in gastruloids (Fig. 3j). These data indicate that gastruloids can undergo differentiation into two distinct cardiomyocyte fates: either a tdTomato positive second heart field fate (outflow tract, right ventricle for example) or a first heart field enriched tdTomato negative fate.

Flow cytometry in E10.5/E11.5 mouse embryos showed less variability with between 0.9 and 1.6% of tdTomato+ cells. In addition, staining of mouse embryos with cTnT antibody showed that Mef2c-AHF-enhancer-Cre derived (tdTomato+) cardiomyocytes accounted for about 22.9% of all cardiomyocytes in embryos (Fig. 3k). These data validate the existence of at least two subpopulations of cardiomyocytes in gastruloids with proportions significantly different from mouse embryos.

To further validate these data, we performed *in situ* hybridization with cardiomyocyte specific probes followed by flow cytometry using the HCR approach (see Methods). We found a significant shift of the cloud of cells in the flow cytometry chart indicative that a significant proportion of cells *Myl7-*positive alone (corresponding to clusters 6, 8 and 9 of the sc-RNAseq data) or in combination with *Myl2* (differentially expressed in cluster 8) in gastruloids at day 11 (Fig. 3l). This result suggests that *Myl7+, Myl2+* ventricular-like cardiomyocytes and *Myl7+Myl2-* non-ventricular cardiomyocytes are present in gastruloids. This result further validates the scRNAseq analysis and confirms the existence of different subpopulations of cardiomyocytes in gastruloids.

### Skeletal myogenesis in gastruloids

To ascertain whether gastruloids could generate skeletal muscle cells, we analyzed the two clusters (clusters 16 and 18) with expression of genes associated with myoblast differentiation (Fig. 4a). Differential gene expression between clusters 16 and 18 was not conclusive (Supplementary Table S3). We thus decided to investigate in detail the genes involved in cardiopharyngeal and/or somitic mesoderm muscle differentiation and their expression in the two myoblast clusters (Fig. 4b). We noticed an increased expression of genes associated with muscle precursors such as *Pax7* and *Myf5* in cluster 16 compared to cluster 18, as shown by the violin plots (Fig. 4c-d). While *Met* has been shown to be downregulated along differentiation in some particular cardiopharyngeal or somites derived myoblasts ^53, 54^, *Met* was expressed in more cells and at higher levels in cluster 16 than in cluster 18 (Fig. 4e). In contrast, markers of more committed myoblasts such as *MyoD*, *Myog* and *Myh3* were enriched in cluster 18 (Fig. 4f-h). These data indicate that myogenesis occurs in the gastruloid model and that clusters 16 and 18 express markers of muscle precursors and committed myoblasts, respectively.

**Figure 4:**
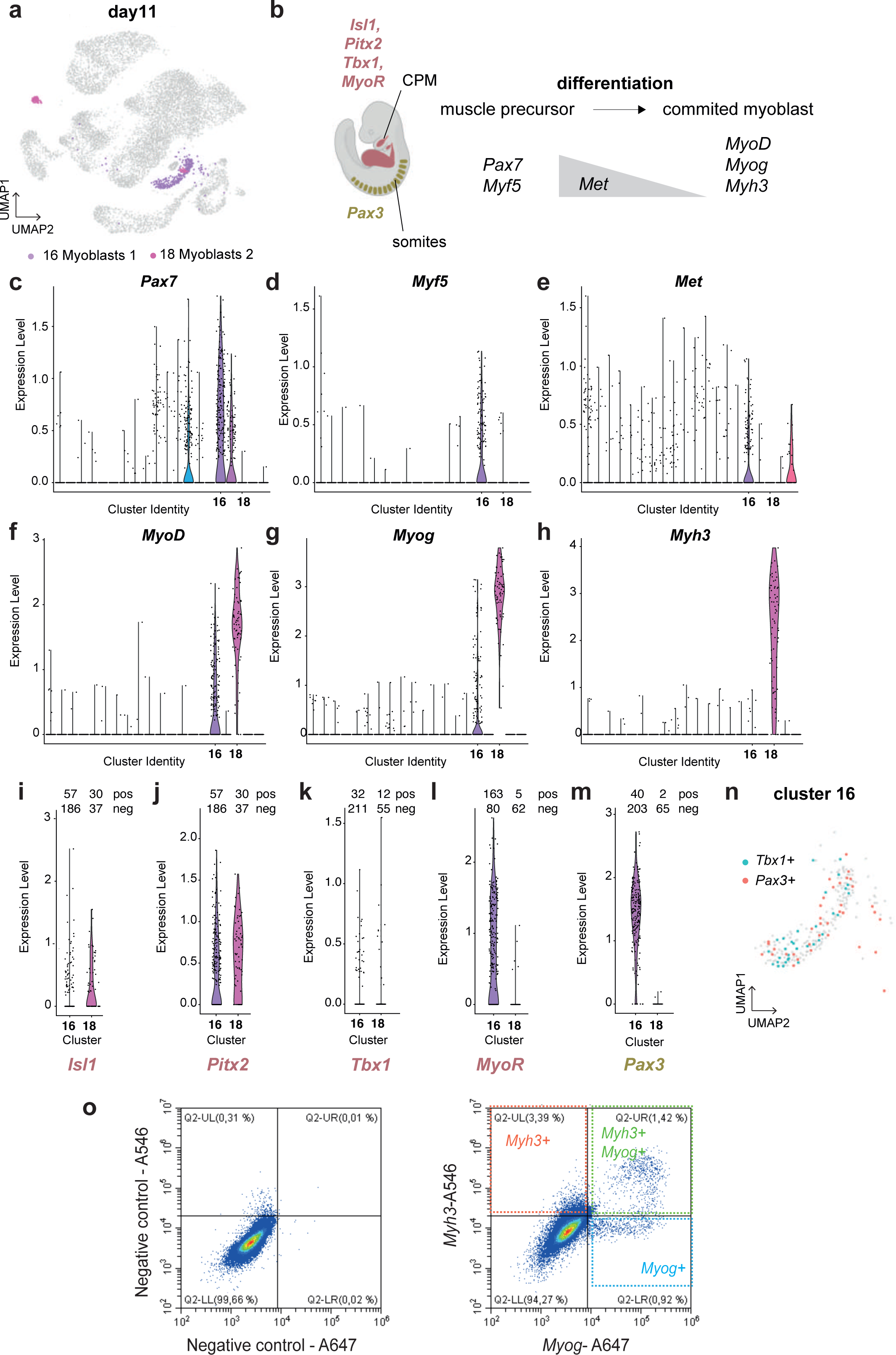
Skeletal myogenesis takes place in gastruloids from the cardiopharyngeal and somitic mesoderm. **a.** UMAP representation of day 11 myoblast clusters highlighted in purple (cluster 16) and in pink (cluster 18). **b.** Scheme of the genetic regulation of skeletal myogenesis in mouse from the cardiopharyngeal (red) or somitic mesoderm (green). Skeletal myogenesis starts from muscle precursors expressing *Pax7* and *Myf5* that differentiate toward committed myoblasts expressing *MyoD*, *Myog* and *Myh3*. **c-h.** Violin plots showing the level of expression of *Pax7* (**c**), *Myf5* (**d**), *Met* (**e**), *MyoD* (**f**), *Myog* (**g**) and *Myh3* (**h**) in the different clusters of gastruloids at day 11. **i-m.** Violin plots showing the level of expression of *Isl1* (**i**), *Pitx2* (**j**), *Tbx1* (**k**), *MyoR* (**l**) and *Pax3* (**m**) specifically in the two myoblast clusters (clusters 16 and 18) at day 11. **n.** UMAP representation of day 11 cluster 16. Blue and red dots represent *Tbx1* and *Pax3* expressing cells, respectively. **o.** FACS analysis of the combined expression of *Myh3* and *Myog* expression in gastruloid cells at day 11 after HCR experiment. Negative controls with no probe are shown on the left.

In order to dissect the cardiopharyngeal versus somitic origin of these myoblasts, we investigated the expression of transcription factors that were specific to either the cardiopharyngeal or somitic muscle progenitors (Fig. 4b). Expression of the cardiopharyngeal mesoderm genes, *Isl1*, *Pitx2*, *Tbx1* and *MyoR*, was found in both cluster 16 and 18 but only in a small number of cells (Fig. 4i-l). *Tcf21* was not expressed in the myoblast clusters, likely having already been downregulated. *Pax3* expression was also recorded in about 40 out of 243 cells of cluster 16 and in 2 out of 67 cells of cluster 18 (Fig. 4m). Deeper analysis of cluster 16 showed a mutually exclusive expression of *Tbx1* and *Pax3* in the population of muscle precursors (Fig. 4n), suggesting the activation of distinct myogenic programs. Our scRNA-seq analysis at day 11 thus indicates that both cardiopharyngeal and somitic progenitors undergo myogenesis in the gastruloid model.

To validate this data and further support the existence of myogenesis in gastruloids, we performed *in situ* hybridization with skeletal myocyte specific probes followed by flow cytometry using the HCR approach (see Methods). Flow cytometry interestingly showed a shift in the cloud of cells with two cell populations expressing *Myog* (Fig. 4o). As found in the single-cell data, we identified a subpopulation of cells expressing *Myog* alone (cluster 16) and a subpopulation of cells expressing *Myog* and *Myh3* (cluster 18). Overall, these observations demonstrate the existence of convergent cardiopharyngeal and somitic myogenic programs in the gastruloid model.

### Muscular trajectories are found in gastruloids over time

Time series scRNA-seq data allows the reconstruction of transcriptional trajectories from a progenitor cell state toward differentiated cell states ^55^. We hypothesize that we could similarly infer the transcriptional trajectories from cardiopharyngeal mesoderm progenitors toward cardiomyocyte and myoblast cell states. We merged the single-cell data from the 4 different time points to create a transcriptional time series of the growing gastruloids (Fig. 5a). We first used CellRank (see Methods) ^56^. We focused on a differentiated cell state (called macrostate), that includes cells from the cardiomyocyte cluster 8 from the day 11 dataset, and computed the fate probabilities towards this macrostate. Interestingly, we found high fate probability with cells from clusters 6 and 9 from day 11, as well with clusters annotated as cardiomyocytes at day 6 (Fig. 5b). We then focused on a second macrostate that includes cells from the myoblast cluster 18 from the day 11 dataset. We found high fate probabilities with cells from the myoblast cluster 16 also from day 11 (Fig. 5c). These data indicate the existence of muscular trajectories toward the cardiac and skeletal muscle states.

**Figure 5:**
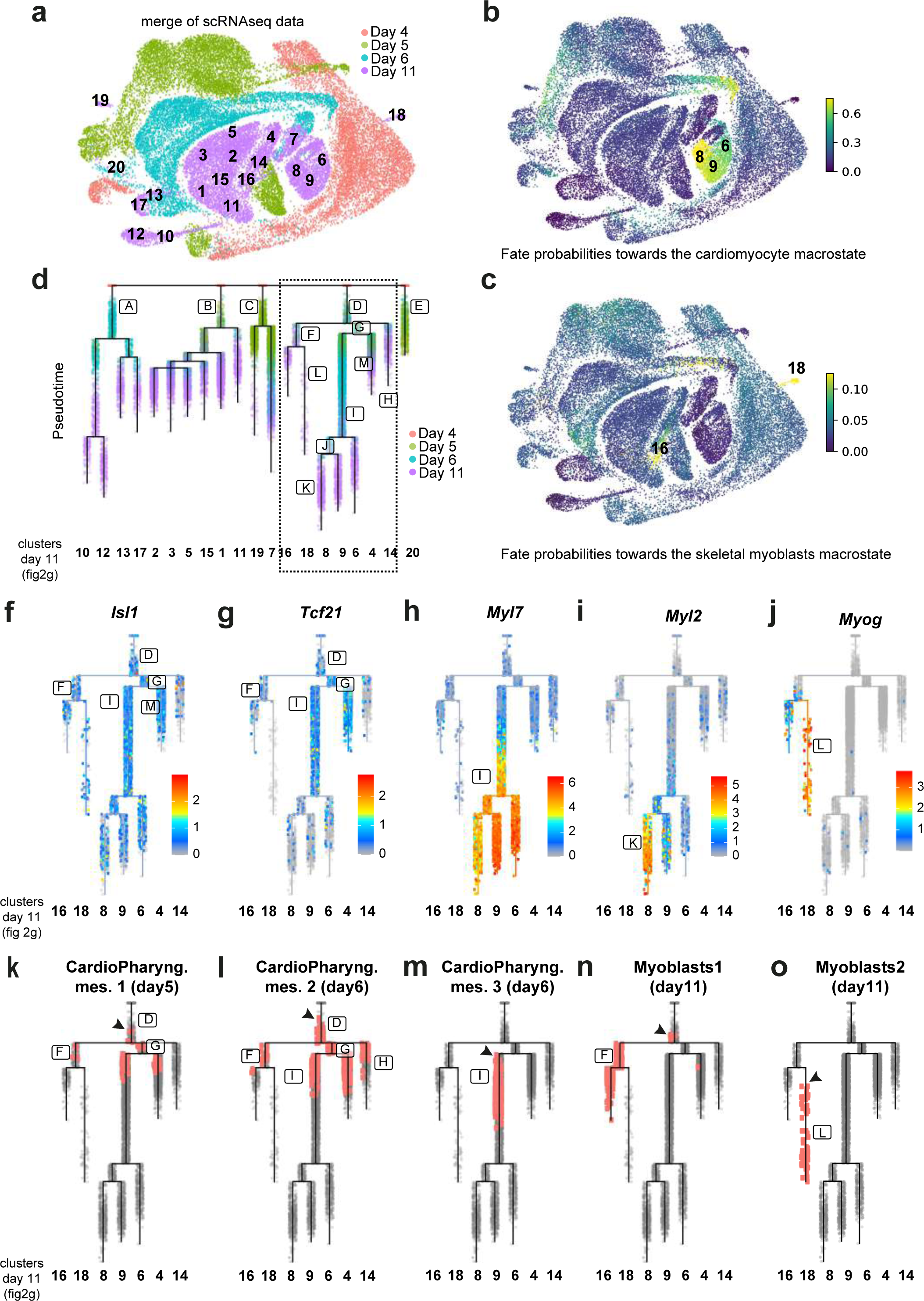
Myogenic trajectories are found in gastruloids over time. **a.** UMAP representation of all gastruloid cells merged from day 4 (red), day 5 (green), day 6 (blue) and day 11 (purple). Numbers represent identified clusters of day 11 (as defined in Fig. 2g). **b-c.** Fate probabilities towards the cardiomyocyte macrostate (**b**) or towards the skeletal myoblast macrostate (**c**). Scale bars represent absorption probabilities. Numbers represent key cardiomyocytes and myoblast clusters of day 11 **d.** URD trajectories showing transition from cells at day 4 (root) towards cells at day 11 (tips). Framed letters represent branches along the tree. The dotted box represents the lineage branch detailed in f-o. **f-j.** Expression of *Isl1* (**f**), *Tcf21* (**g**), *Myl7* (**h**), *Myl2* (**i**) and *Myog* (**g**) across URD trajectory. Scale bars represent expression levels. **k-o.** Representation of clusters’ cells across URD trajectory. Red dots represent cells of the cardiopharyngeal mesoderm (CardioPharyng. mes.) cluster 1 of day 5 (in **k**, Fig. 2c, cluster 1), of the cardiopharyngeal mesoderm (CardioPharyng. mes.) cluster 2 of day 6 (in **l**, Fig. 2e, cluster 5), of the cardiopharyngeal mesoderm (CardioPharyng. mes.) cluster 3 of day 6 (in **m**, Fig. 2e, cluster 8), of the myoblast cluster 1 of day 11 (in **n**, Fig. 2g, cluster 16) and of the myoblast cluster 2 of day 11 (in **o**, Fig. 2g, cluster 18).

To reconstruct fate decision trees for the various populations, we used URD ^57^ (see Methods) as a computational reconstruction method, with the merge of all time points (days 4, 5, 6 and 11). URD analysis computed transcriptional trajectories from cells at day 4 (named root) towards the different clusters at day 11 (named tips) (Fig. 5d). We observed 5 branches emerging from day 4. The first branch (branch A) heads towards the ectodermal lineages (clusters 10, 12, 13 and 17) while the last branch (E) heads towards the endoderm lineages (cluster 20). The 3 other branches (branches B, C and D) are mesodermal, and include the endothelial (cluster 7) and hematopoietic lineages (cluster 19). Notably, one of these mesodermal branches, branch D, heads towards the cardiomyocytes (clusters 6, 8 and 9) and myoblast clusters (clusters 16 and 18). Thus, the URD analysis shows that transcriptional trajectories can be found toward both the cardiomyocytes and skeletal muscles.

To further dissect these trajectories, we specifically focused on branch D. This branch also includes clusters 4 and 14. These two clusters were previously defined as mesodermal derivatives (Fig. 2g). We investigated gene expression in branches M and H to characterize further these clusters (Fig. 5d). We found that branches M and H showed *Gata6* expression as well as *Hand1* and *Mab21l2*, which mark the juxta-cardiac field ^40, 41^ (Supplementary Fig. S10). We also showed expression of *Tbx18* and *Wt1* (Supplementary Fig. S10), which mark the proepicardial organ and their derivatives ^58^. These data suggest that these clusters might contain epicardial cells. We then investigated how key markers of the cardiopharyngeal mesoderm and their derivatives are expressed in the tree. We found that *Isl1*, *Tcf21,* but also *Lhx2* and *Ebf3* are expressed in branch D, in the root as well as along the trajectories towards the cardiomyocyte and myoblast clusters (Fig. 5f-g and Supplementary Fig. S10). *Myl7* is specifically expressed in the three cardiomyocyte clusters (6, 8 and 9) and the cardiomyocyte branching point I (Fig. 5h), while *Myl2* is expressed after the branching bifurcation only in cluster 8 (Fig. 5i). As expected *Myog* is expressed in branch L that heads towards the myoblast cluster 18 (Fig. 5j). *Tbx1*, *MyoR,* markers of the cardiopharyngeal muscle progenitors, *Pax3,* markers of the somitic muscle progenitors, and the early myogenic transcription factor *Myf5* are all expressed in branch F that heads towards the two myoblast clusters 16 and 18 (Supplementary Fig. S10). We also explored where cells of clusters from days 5 and 6 would be found in this tree (Fig. 5k-m). Strikingly, cells of the cardiopharyngeal mesoderm 1 cluster, identified at day 5, were found almost at the root of the tree (branch D), before the branching bifurcation between the cardiomyocytes and myoblast lineages (Fig. 5k). Similarly, cells of the cardiopharyngeal mesoderm 2 cluster, identified at day 6, were found before the branching point (branch D) (Fig. 5l). However, we found that cells of the cardiopharyngeal mesoderm 3 cluster, identified at day 6, were restricted to the cardiomyocyte lineages (branch I). These cells are located after the branching bifurcation between the cardiomyocytes and myoblast lineages (branch D) (Fig. 5m). We then focused on the two myoblast clusters identified at day 11. Cells of cluster 16 were also found in branch F, before the branching point that heads toward the myoblast clusters 16 and 18 (Fig. 5n). Cells of cluster 18, on the other hands, were found only after branch bifurcation, in branch L (Fig. 5o). These data further confirm that myoblast cells from cluster 18 are in a more differentiated state that cells from cluster 16. Together these trajectory inferences show that muscular transcriptional trajectories are found in gastruloids. They also show that a specific branch of the tree includes the cardiopharyngeal mesoderm and heads towards both cardiomyocytes and myoblasts clusters.

## Discussion

Our results provide evidence that cardiopharyngeal mesoderm markers are expressed in a similar spatio-temporal pattern in gastruloids and the mouse embryo. We also showed that cardiopharyngeal mesoderm differentiates in the gastruloid model into different subpopulations of cardiomyocytes as well as skeletal myoblasts. Gastruloids thus faithfully recapitulate the developmental timing of cardiopharyngeal mesoderm specification and differentiation (Fig. 6).

**Figure 6:**
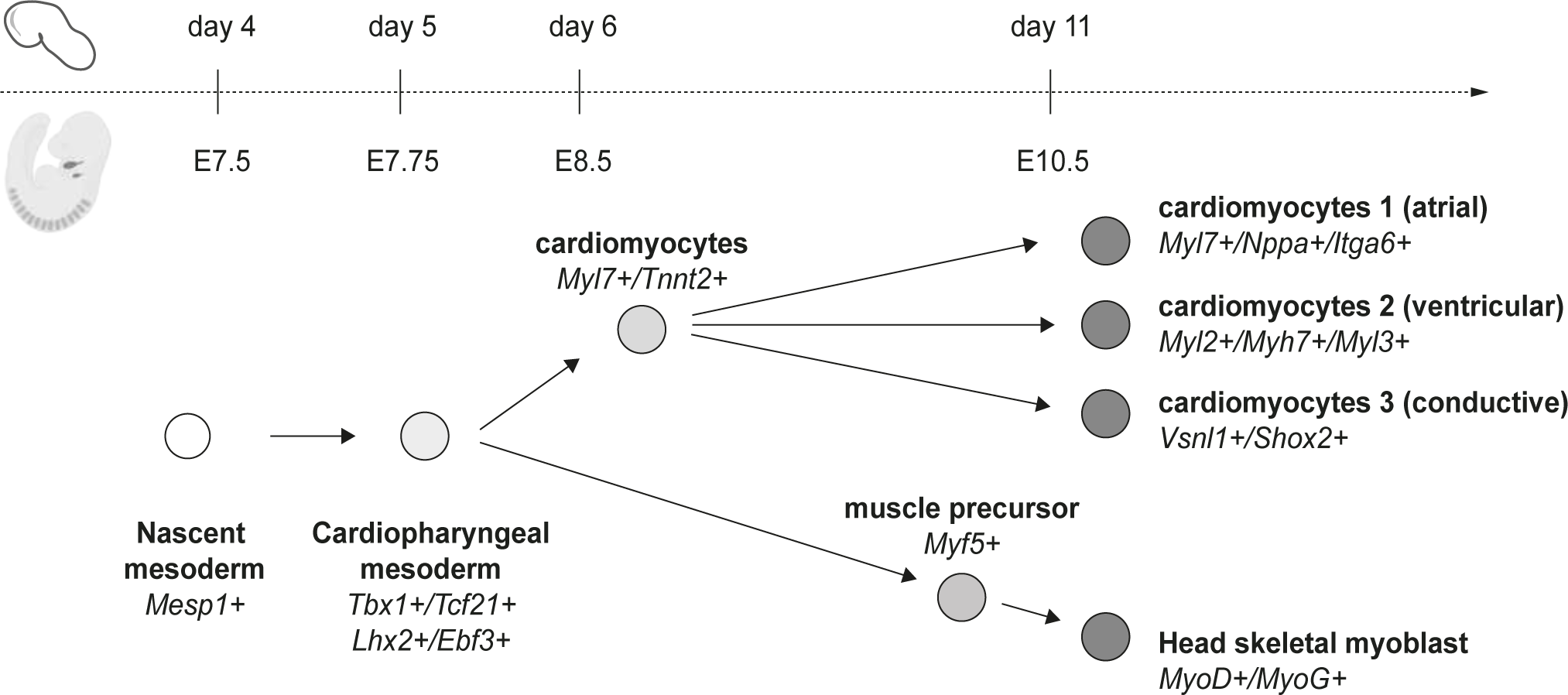
Schematic representation of the developmental timing of cardiopharyngeal mesoderm specification and differentiation in gastruloids in comparison with the mouse model.

To the best of our knowledge, here we provide the first *in vitro* model for parallel differentiation of cardiomyocytes and skeletal muscles. Indeed, in addition to cardiomyocyte differentiation previously reported by Rossi and colleagues ^34^, our study demonstrates that skeletal myogenesis occurs in gastruloids. Using scRNA-seq analysis we found that the mesodermal origin (cardiopharyngeal or somitic) was not reflected in the clustering of skeletal muscle progenitor cells. In contrast, we identified two clusters that segregate different states of myogenesis. This finding indicates that transcriptional similarity due to the state of myogenesis might predominate over its developmental origin, raising challenges in addressing whether myogenesis in gastruloids derives from cardiopharyngeal or somitic mesoderm. We first showed the expression of key markers of cardiopharyngeal progenitors such as *Tcf21*, *Tbx1*, *Lhx2*, *MyoR*, and *Pitx2* ^16, 20, 23, 24^. Additionally, we identified transcriptional trajectories from the cardiopharyngeal mesoderm towards both myoblasts and cardiomyocytes. These results together with the close proximity between the cardiomyocytes and myoblasts in gastruloids indicate that skeletal myogenesis, at least in part, occurs from the cardiopharyngeal mesoderm. Based on the exclusive expression of *Tbx1* (cardiopharyngeal marker) and *Pax3* (somitic marker) in myoblast cluster 16 we propose that myoblasts also derive from somitic mesoderm and that gastruloids thus allow skeletal myogenesis of both head and trunk programs. Furthermore, transcriptomic trajectories computed with URD showed a branch with both cardiomyocytes and myoblasts. But expression of both somitic (*Pax3*) and cardiopharyngeal mesoderm markers is found in that particular branch. This highlights the known limitations with the use of single-cell transcriptomics approaches for lineage inference. ScRNAseq alone provides information only on gene expression, and attempts to reconstruct lineage trajectories from transcriptomic data can be misleading and do not allow a complete lineage reconstruction ^59^. Only clonal analysis will confirm whether cardiopharyngeal mesoderm bipotent progenitors are found in gastruloids but this requires specific tools and analysis.

Interestingly, different subpopulations of cardiomyocytes were identified in gastruloids. Gene expression analysis suggests that these subpopulations correspond to ventricular (*Myl2+*, *Myl3+*, *Myh7+*), atrial (*Myh6*+, *Nppa*+) and immature/conductive cardiomyocytes *(Shox2+, Vsnl1+)* ^44–47, 49^. However, atrial and ventricular myocardium share many genes in common during development. They are distinguished later by acquisition of their mature electro-mechanical properties (conduction, voltage tension, some structural proteins and ions channels). Thus, few genes are differentially expressed *in vivo*: *Myl7* is only restricted to atrial myocardium in fetal mouse embryo, while *Myl2* is restricted in ventricular myocardium. Finally, *Nppa* is detected in the atrium and the ventricle during development, but its expression became atrial specific immediately before birth ^60^. It is thus not clear how mature are the cardiomyocytes in gastruloids. Further work will be required to characterize these different subpopulations, their spatial organization and their origin.

Our results reveal robust myogenic differentiation to heart and head muscle in the gastruloid system. This is supported by reproducibility with similar observations from different replicates. We also showed redundancy with similar conclusions obtained when using scRNA-seq, RNAscope, HCR, lineage tracing and immunofluorescence. The use of different mESC lines showed the robustness of the gastruloids for the specification of the cardiopharyngeal mesoderm. However, our gastruloid model also presents some limitations. In contrast to gastruloids from human induced pluripotent stem cells, which have been shown to form a heart-tube like structure ^33^ or human cardioids which have been shown to form multi-chambers heart like structures ^61^, our murine gastruloid model does not show robust heart morphogenesis. We also have not observed cardiac crescent like structures as was previously described ^34^. This indicates that further studies and protocol adaptations are required to form these different chamber structures and to improve cardiac morphogenesis. Interactions between pharyngeal endoderm and the heart are known to be crucial for heart morphogenesis ^62, 63^. We hypothesize that with the robust development of cardiopharyngeal mesoderm in our study, the spatial proximity between the cardiac derivatives and the endoderm/gut structure might be perturbed. Affecting the proximity with the endoderm might impact cardiac morphogenesis.

Morphogenesis and physical forces are also different in gastruloids, compared to embryos. Our results indicate that while cardiopharyngeal mesoderm is formed in gastruloids, there is no indication of pharyngeal arch morphogenesis. Further studies are required to determine how these transient structures are formed in the embryo. The gastruloid model offers a possibility to reconstruct such processes and investigate whether physical constraints could play a role as shown for example with matrigel embedding for somitogenesis ^35, 36^. Additionally, we know that pharyngeal arches are also colonized by neural crest cells, which are largely absent in the gastruloid model ^32^. Adapting the protocol to support both neural crest cell and cardiopharyngeal mesoderm development might improve pharyngeal arch morphogenesis. While cranial/cardiac neural crest cells are known to be critical for outflow tract and valve development ^64–66^ as well as for the patterning of head skeletal muscles ^67^, our results support that neural crest cells are not required for cardiac and cardiopharyngeal mesoderm specification.

As previously reported, we have also noted variability among the lines of mESC used. Notably, mESC derived from *Mef2c-AHF-Cre/tdTomato* blastocysts seem to have less potential for skeletal myogenesis. We speculate that it could be due to a different genetic background. Alternatively, established and rederived mESC lines from mouse embryos might not strictly have the same pluripotency status. In particular, we found that different mESC clones exhibited a truly different beat and myogenic potential. Due to the lack of specific markers, it was also difficult to address whether other cardiopharyngeal mesoderm derivatives such as connective tissue or smooth muscle ^51, 53, 68^, could develop in gastruloids.

Several reports have already indicated that gastruloids, somitoids, segmentoids and embryonic trunk-like structures, from mouse or human pluripotent stem cells, could faithfully recapitulate paraxial mesoderm early specification with the formation of somites ^35, 36, 69–71^. None of these studies however showed the initiation of myogenesis. However, most of their protocols stopped after 5 to 7 days of culture whereas we have extended the culture of gastruloids up to 15 days. Our kinetics of expression showed that markers of myogenesis are initiated only after day 8. This suggests that somitoids, segmentoids and embryonic trunk-like structures might also have the potential to undergo myogenesis if cultured for a longer period of time.

Together our results show the relevance of the gastruloid model to investigate cardiopharyngeal mesoderm specification and differentiation, and demonstrate that this *in vitro* model could be used as an alternative model to complement or reduce the use of animal models. The protocol is relatively simple and allows the culture of a large number of gastruloids in a single experiment. It thus offers a powerful *in vitro* platform to model congenital heart disease and muscle dystrophies (22q11.2DS for example), and to deconstruct cardiopharyngeal mesoderm specification as well as myogenesis. Cardiopharyngeal mesoderm development is still not fully understood *in vivo*. The field lacks reporter lines and face difficulties to carefully track the progenitors and their derivatives along a course of about 3 to 5 days, from the emergence of the progenitors at gastrulation to their differentiation into cardiomyocytes or skeletal myoblasts. Cardiopharyngeal mesoderm-containing gastruloids additionally offer the possibility for high resolution live imaging and cell tracking. Furthermore, the model will likely be useful for large scale genetic and drug screens to better understand and potentially identify therapeutic strategies for congenital diseases affecting head and heart muscles. In this context, we expect gastruloids to become indispensable tools in the field of cardiac and skeletal muscle specification to address the mechanisms of lineage specification as early as gastrulation.

## Methods

### Mice colonies

*Mef2c–AHF-Cre* and *Rosa-tdTomato* (*Gt(ROSA)26Sor^tm^*^9^(CAG–tdTomato)*^Hze^*) mice were previously described ^50, 72^. We used CD1/Swiss mice (Charles River) as wildtype animals. Mouse colonies were maintained in certified animal facilities (agreement #C 13 013 08) in accordance with European guidelines. The experiments were approved by the local ethical committee (CEBEA) and the study is compliant with all relevant ethical regulations regarding animal research (Ministère de l’Education Nationale, de l’Enseignement Supérieur et de la Recherche; Authorization N 32-08102012).

### Cell culture

R1/E #1036 mouse embryonic ES cells were purchased from ECACC (cat. Number 07072001). Zx1 mouse ESCs were obtained from M. Kyba ^73^. *Mef2c-cre Rosa tdTomato* cells were derived in the lab from E3.5 bastula collected from mouse ^50^. All cell lines were maintained on 0.1% gelatin coated labware in GMEM-BHK21 medium (Gibco ref 21710-025) supplemented with 8% Fetal Bovine Serum (Pansera, ref P30-2602), 1% non-essential amino acids (Gibco 11140-035), 1% Sodium Pyruvate (Gibco ref 11360-039), 1X Penicillin/Streptomycin (Gibco ref 15070-063), 0.1 mM beta-mercaptoethanol (Sigma ref M3148-100ML). This basal medium was completed with 5ng/ml recombinant mouse LIF (Sigma ref LIF2050), 3µM CHIR99021 (Sigma ref SML1046-5MG) and 1µM PD0325901 (Sigma ref PZ0162-5MG). Cells were passaged at least twice but no more than 10 times before differentiation.

### Gastruloid production

Gastruloids were produced according to previously described protocols ^32, 34^ with slight modifications (centrifugation for aggregation and culture in N2B27 medium after day7). Briefly, mES reaching a 60% confluency were dissociated with 0,5% Trypsin-EDTA (Gibco ref 25300-054). After enzyme neutralization, cells were carefully washed 2 times with D-PBS (Gibco ref 14190-094) and resuspended in N2B27 medium (Neurobasal Medium (Gibco ref 21103049), DMEM/F12 (Gibco ref 11320074), 0,5% B27 supplement (Gibco ref 17504-044), 0,5% N2 supplement (Gibco ref 17502-048), 1X Glutamax (Gibco ref 35050-038), 0.1 mM beta-mercaptoethanol (Sigma ref M3148-100ML). 300 to 600 individual cells were distributed into 40µl of N2B27 in each well of a Ultra-Low-Attachment 96 well round bottom plate (Corning ref 0707). Plates were then centrifuged at 100g for 5 min and placed in a 5% CO2 incubator during 48 hours. Then, 150 µl of N2B27 with 3µM CHIR99021 were added in each well and plates were incubated for 24h. After this period, medium was replaced by N2B27 without CHIR99021 for another 24h. Then from day 4 to day 7, N2B27 was supplemented with 30 ng/ml bFGF (R&D ref 233-FB), 5 ng/ml VEGF (Gibco ref PHC9394) and 0,5mM Ascorbic Acid (Sigma ref 255564-5G) and culture plates were placed on an orbital shaker at 100 rpm. After day 8, N2B27 was used and changed every other day.

### Gene expression analysis by QPCR

Day 0 undifferentiated mES cells and different time points gastruloids were collected during the experiments. Samples were then lysed in Buffer RA1 and RNA was extracted with NucleoSpin RNA kit (Macherey-Nagel ref 740902.250). Total RNAs were then retrotranscribed to cDNA using AffinityScript Multiple Temperature cDNA Synthesis Kit (Agilent ref 200436) and QPCR were run on QuantStudio 5 according to PowerUp™ SYBR® Green instructions (Thermo ref A25742). List of QPCR primers can be found in supplementary information (Table S4). Data were all standardized using *Tbp* as a housekeeping reference gene and on day 0 results and gene expression fold changes were calculated with the ΔΔCt method.

### Single cell RNA-sequencing analysis

Gastruloids collected on day 4, 5, 6 and 11 were dissociated by incubation 10 min at 37°c in 0.5% Trypsin-EDTA. Cells were washed 2 times with D-MEM + 10% FBS and filtrated through a 40µm cell strainer. Cell count and viability were determined on a Countess II (Thermo ref AMQAX100). Single cell-RNA seq libraries were made from 10,000 individual cells using 10X Genomics Chromium Next GEM Single Cell 3’ kit v3.1 (PN-1000268) and the Chromium Controller. Libraries were then sequenced on an Illumina NextSeq 500 at the MMG Sequencing platform.

Single-cell analysis was performed using 1) Seurat, the R toolkit for single-cell genomics ^74^, 2) CellRank, to discover the cellular dynamics ^56^, and 3) URD to obtain transcriptional trajectories ^57^. The analysis was implemented in R v4.1.3 and Python v3.9.10.

### Preparation of the Pijuan-Sala et al. atlas

The Pijuan-Sala et al. atlas has been constructed from mouse embryos to describe gastrulation and early organogenesis ^37^. It contains single-cell analysis at multiple developmental stages between 6.5 to 8.5 days post-fertilization ^37^. The atlas was subsetted to select the cells with more than 200 feature gene counts and the features detected in at least 4 cells. Then, normalization was performed on the feature counts with the “LogNormalize” method. An atlas must be comparable to the dataset under study. Here, and because we work on gastruloids mimicking gastrulation, the stage “mixed_gastrulation” and the cells annotated “endoderm” or “extraembryonic endoderm” were removed from the atlas. Finally, the identification of the highly variable genes, the data scaling, and the PC analysis and UMAP dimensional reduction were performed as detailed in step 2 of the next section.

### Processing of the four individual single-cell datasets

Four single-cell RNA sequencing datasets were generated from gastruloids’cells collected at days 4, 5, 6, and 11. Each dataset was analyzed with the R package Seurat v4.1.1 using the following six steps ^74^.

#### 1. Quality controls

Low quality cells were filtered out with different cutoffs on gene expression. All these cutoffs were set manually and independently on each dataset. The proportion of transcripts mapping to mitochondrial and ribosomal genes were evaluated. We set cutoffs on high mitochondrial and low ribosomal gene expression. In addition, we set two cutoffs on the UMI counts of each cell. Finally, the data were subsetted to keep cells with more than 200 feature gene counts and features detected in at least 4 cells.

#### 2. Preprocessing workflow

Feature counts were normalized using the “LogNormalize” method. Highly variable genes were identified with the “vst” selection mode, and selected the 2000 most variable genes. Then, the data were scaled on all the features and Principal Component (PC) analysis dimension reduction was performed. The 30 first PCs were kept for the rest of the analysis. To visualize the data, Uniform Manifold Approximation and Projection (UMAP) was applied ^75^. Finally, the normalized data were structured with a shared nearest neighbor graph. The number of nearest neighbors was set to 20 (default). We used the Leiden algorithm with the resolution set to 1 to cluster the graph ^76^.

#### 3. Doublets detection

Doublets detection was carried out with the R package DoubletFinder v2.0.3 ^77^. A cell type annotation transfer was first applied. Indeed, according to DoubletFinder recommendations, a cell type annotation transfer allows a better estimate of the number of detectable doublets ^77^. The Seurat tutorial “Mapping and annotating query datasets” was applied to transfer the annotation of cell types, using the atlas of Pijuan-Sala et al. as a reference ^37, 74^. We estimated the number of heterotypic doublets from the percentage of doublets of the used reagent kit of 10x Genomics Chromium 3’ chemistry v3.1. In our case, the rate of doublets amounts to 8%. For each dataset independently, the pK parameter was optimized with the function paramSweep_v3. The first 30 PCs were considered. As a result of the process, every cell was labeled as either singlet or doublet. We selected the cells labeled as singlets, and reapplied the Step 2 preprocessing workflow to the singlet subset.

#### 4. Cell type annotation transfer from the atlas

Like for doublets detection, cell type annotations were transferred using the Seurat functions with default parameters ^74^.

#### 5. Global structure-preserving UMAP plots

The global structure-preserving UMAP dimension reduction was set up using the R package uwot v0.1.14 ^75^. The aim was to keep a similar vizualisation structure among the different datasets. The reference global structure was created using the Pijuan-Sala et al. atlas ^37^. We first integrated our dataset with the Pijuan-Sala et al. atlas using the Seurat framework for datasets integration with default parameters.

Besides, UMAP dimension reduction was carried out on a subset of the atlas containing 200 cells per cell type. From the resulting UMAP coordinates, the centroid of each cell type was calculated. We randomly assigned positions to all the cells of the integrated object (dataset and atlas) around the centroid of their corresponding cell type. The UMAP algorithm was run on the integrated object, with the random positions around cell type centroids as the starting points. In a following step, the UMAP algorithm was run again on the dataset only, with the dataset’s UMAP coordinates in the integrated object as the starting points.

#### 6. Differential expression analysis

Differentially expressed genes (DEGs) were determined using a Wilcoxon Rank Sum test with the Seurat function FindAllMarkers. The function compares each cluster (group 1) to the rest of the data (group 2). Default values were used for min.pct, logfc.threshold, and return.thresh parameters: min.pct allows to speed up the function by selecting features that are expressed over a given proportion of cells within each group, return.thresh selects p-values lower than a given threshold, and logfc.threshold selects features showing at least a 2-fold change between the two groups of cells. We selected the positive markers only.

For each cluster, we also retrieved the most represented cell type and its corresponding percentage of cells in the cluster (see Table S1).

### Processing of the merged single-cell datasets

The four single-cell datasets were also merged. The datasets were reduced to the previously identified high-quality cells (see steps 1 and 3 of the processing pipeline). The identification of the highly variable genes, the data scaling, and the PCA and UMAP dimensional reductions were performed as described in step 2 from the previous section. Steps 4 to 6 were then applied to the merged dataset.

### Day11 differential expression analysis for cardiomyocytes and myoblasts characterization

Additional differential expression (DE) analyses were performed on two subsets of the day11 dataset. First, to better characterize the cardiomyocytes, the clusters identified as cardiomyocytes (ie. clusters 6, 8, and 9) were compared. In this case, each cluster was compared to the two others. The cluster 18 of day 11 was also identified as a cluster of cardiomyocytes (see Table S1). However, the day 11 is poorly annotated because the reference atlas does not reach this time points. We manually assigned the “myoblast” label to this cluster. DE analysis was performed between the clusters 16 and 18, two clusters of myoblast cells. In both analyses, the parameters used for the DE analyses were the same as detailed in step 6 of the previous section.

### Data preparation for file format conversion

CellRank and URD necessitate different file format. We started from raw data and removed the cells and features that were filtered out during the preprocessing analysis of the four datasets (step 1 and 3). The obtained datasets were then merged and the feature counts were normalized using the “LogNormalize” method. The object was saved as a .rds file (R).

### Cell Rank time series framework

In order to use CellRank ^56^ v1.5.1, the Seurat ^74^ object (R) was converted to an AnnData ^78^ v0.7.8 object (Python). The prepared dataset was converted into .h5ad (Python) (see Data preparation for file format conversion section).

The .h5ad object was preprocessed with loading, normalization, logarithmization, highly variable genes identification, PC analysis, and KNN graph computing, using the Scanpy v1.9.1 tool ^79^. We added the metadata of UMAP coordinates, cell type assignation, and Leiden clustering to each cell. CellRank was used with the Waddington Optimal Transport method (WOT), the time series flavor of CellRank ^80^. The WOT kernel was then computed. Due to insufficient proliferation genes in highly variable genes, we were not able to compute the initial growth rates. Hence, according to CellRank specifications, the transition matrix was computed with the parameter growth_iters = 3 ^56^. A symmetric transition matrix was also computed based on transcriptomic similarity, called connectivity kernel. WOT kernel and connectivity kernel both contributed to the estimator with the respective proportions of 0.8 and 0.2. The estimator was initialized with the Generalized Perron Cluster Cluster Analysis (GPCCA) method ^81^. Based on Schur decomposition, 6 macrostates were computed ^82^. Then, fate probabilities were computed towards the macrostates with the function compute_absorption_probabilities, with default parameters. Putative driver genes were identified with the function compute_lineage_drivers.

### Trajectory inference

The R package URD v1.1.1 was used for trajectory inference ^57^. The prepared dataset was converted into an urd object (see Data preparation for file format conversion section). The variable features within each dataset were identified with the findVariableGenes function. A KNN graph was constructed with distances between cells’ gene expression, considering the 60 nearest neighbors of each cell. We removed the outliers, identified visually according to the distance between the first a^nd^ the 60th neighbours of each cell. 92 outlier cells were removed from the analysis. Then, a diffusion map was computed with the parameters sigma=3 and knn=60. The pseudotime was simulated 5 times starting from any cell of the day 4 and the results were averaged. All the clusters of day11 were defined as tips. The function pseudotimeDetermineLogistic was used with the parameters optimal.cells.forward=40 and optimal.cells.forward=80 to bias the input transition matrix of the random walk simulations. 10000 random walk simulations were carried out. Finally, the tree was constructed by merging the tips as soon they share a common path among all the random walk simulations. To build the tree, the parameters of the function buildTree were set as follows: bins.per.pseudotime.window=8, cells.per.pseudotime.bin=28, divergence.method=‘preference’, p.thresh=0.001.

### RNAscope experiments on sections

Gastruloids collected at different time points were washed once in PBS + 0,01% Tween 20 (PBST) and then incubated in 4% PFA overnight at 4°C. After 3 washes in PBST, approximately 8 Gastruloids were embedded in heat liquified Histogel (Epredia HG-4000-012). Solidified blocks were then post-fixed with 4% PFA during 1h at room temperature and dehydrated with successive incubations of 30 min in Ethanol / PBST: 70%, 96%, twice 100%. Xylene was then added twice for 15 min at room temperature. Histogel blocs were finally embedded in heat liquid Paraplast plus (Sigma ref P3683-1KG). Tissue sections cut at 7 µm were then processed according to the protocol of the RNAscope Multiplex Fluorescent v2 kit (ACD-Bio cat. no.323110). The following probes were used: *mm-Ebf3-C3* (576871-C3), *mm-Isl1-C2* (cat no. 451931-C2), *mm-Mesp1-C2* (cat no. 436281-C2), *mm-Mesp1-C3* (cat no. 436281-C3), *mm-Myf5-C1* (cat no. 492911), *mm-Myod1-C2* (cat no. 316081-C2), *mm-Nkx2-5-C2* (cat no. 428241-C2), *mm-Tbx1-C1* (cat no. 481919), *mm-Tcf21-C2* (cat no. 508668-C2). Sections were imaged using an LSM800 confocal microscope (Zeiss).

### Flow cytometry

Gastruloids collected on day 11 were dissociated by incubation 10 min at 37°c in 0,5% Trypsin-EDTA. Cells were washed 2 times with D-MEM + 10% FBS and filtrated through a 40µ cell strainer (Falcon 352340). After a PBS wash, 1 million cells were incubated for 20 min at room temperature with Zombie viability dye (Biolegend ref 77143). Then cells were incubated in PBS + 10% FBS to block Fc receptors, followed by a permeabilization/fixation with Cytofix/Cytoperm BD kit (ref 554714). After washing, anti-cTnt antibody (Thermo ref MA5-12960) was incubated at a 1/400 dilution in BD wash buffer during 20 min at room temperature. Then cells were again washed and followed by an incubation with anti-mouse IgG coupled to PE-Cy7 (Biolegend ref 406613) at a 1/200 dilution in BD wash buffer during 20 min at room temperature. After three washes, cells were resuspended in PBS + 1% BSA (Sigma ref A8412-100ML), filtered on a 40µ cell strainer and analyzed on a BD FACS Aria II cytometer.

Gastruloids made with Mef2c-cre Rosa-tdTomato were prepared by the same procedure up to Zombie viability dye labelling and directly analyzed on FACS Aria II to allow sorting of the Tomato positive and negative cells. RNA from sorted cells were extracted with NucleoSpin RNA extraction kit.

### HCR RNA flow analysis

Gastruloids collected on day 11 were dissociated by incubation 10 min at 37°c in 0,5% Trypsin-EDTA. Cells were washed 2 times with PBS + 10% FBS and filtrated through a 40µ cell. After PBS wash, 1 million cells were incubated for 20 min at room temperature with Zombie viability dye. Cells were then fixed by incubation with 4% PFA (EMS15714) during 1h at room temperature. HCR multiplexed RNA detection was realized according to Molecular Instruments protocols. Cells were washed in PBST: PBS + 0,1% Tween 20, resuspended in 70% Ethanol and kept overnight at 4°c. The day after, cells were washed twice in PBST and incubated at 37°c in hybridization buffer. Probes specific for the different genes were then added at 16nM concentration and incubated overnight at 37°c. Cells were then washed 5 times with wash buffer and 2 times with 5x SSC (Gibco ref 15557-044) + 0,1% Tween 20. Then incubation of the cells with amplification buffer was done during 30 min at room temperature. Amplification hairpin h1 and h2 were prepared separately by heating at 95°C for 90 sec and cooling to room temperature for 30 min. Then hairpins were pooled at 60 nM and incubated with cells overnight at room temperature. After 6 washes with 5x SSC + 0,1% Tween 20. Samples were filtered on Flowmi 70µm (Bel-Art H136800070) and analyzed on a Beckman Coulter Cytoflex LX (AMUTICYT platform). The following HCR probes were used: *Myl2-B2*, *Myl7-B3*, *Myh3*-B2 and *Myog*-B1 with the following amplifier: B1-Alexa647, B2-Alexa546 and B3-Alexa488.

## Data availability

Single cell data analysis can be found at: https://github.com/BAUDOTlab/gastruloid_timeserie_scRNA-seq.

## Acknowledgments

We want to thank M. Kyba for providing the Zx1 mES cell line. Single-cell isolation, sequencing and alignment were performed with Veracyte or in the MMG GBIM facility. We thank the cells and animal imaging platform and the animal phenotyping core platform (MMG). Flow Cytometry was performed on the Amu-Flow Cytometry platform (Manon Richaud) and AMUTICYT platform (Stéphane Robert). We thank Lionel Spinelli and the CIML bioinformatics platform for expert advice on bioinformatics analyses. We thank Robert Kelly and Anabela Bensimon-Brito for careful reading of the manuscript.

## Author contributions

L.A., F.L., S.Z designed the experiments. L.A. and C. Choquet performed most of the biological experiments. C. Chevalier performed bioinformatic analysis for all the single-cell RNA-seq data. L.A. performed HCR and FACS experiments. C. Choquet performed RNAscope on gastruloids and mouse embryos. N.N. performed the immunofluorescence on Mef2c-Cre mouse embryos. L.A. and N.N rederived Mef2c-Cre/tdTomato mouse ESCs. A.G. provided technical support. F.L., S.Z. and A.B. wrote the manuscript. All authors read and approved the final manuscript.

## References

1. Nomaru, H. et al. Single cell multi-omic analysis identifies a Tbx1-dependent multilineage primed population in murine cardiopharyngeal mesoderm. Nat Commun 12, 6645 (2021).

2. Buckingham, M., Meilhac, S. & Zaffran, S. Building the mammalian heart from two sources of myocardial cells. Nat Rev Genet 6, 826–835 (2005).

3. Meilhac, S. M., Esner, M., Kelly, R. G., Nicolas, J.-F. & Buckingham, M. E. The Clonal Origin of Myocardial Cells in Different Regions of the Embryonic Mouse Heart. Developmental Cell 6, 685–698 (2004).

4. Kelly, R. G., Brown, N. A. & Buckingham, M. E. The Arterial Pole of the Mouse Heart Forms from Fgf10-Expressing Cells in Pharyngeal Mesoderm. Developmental Cell 1, 435–440 (2001).

5. Zaffran, S., Kelly, R. G., Meilhac, S. M., Buckingham, M. E. & Brown, N. A. Right Ventricular Myocardium Derives From the Anterior Heart Field. Circulation Research 95, 261– 268 (2004).

6. Waldo, K. L. et al. Conotruncal myocardium arises from a secondary heart field. Development 128, 3179–3188 (2001).

7. Mjaatvedt, C. H. et al. The Outflow Tract of the Heart Is Recruited from a Novel Heart-Forming Field. Developmental Biology 238, 97–109 (2001).

8. Lescroart, F., Dumas, C. E., Adachi, N. & Kelly, R. G. Emergence of heart and branchiomeric muscles in cardiopharyngeal mesoderm. Experimental Cell Research 410, 112931 (2022).

9. Diogo, R. et al. A new heart for a new head in vertebrate cardiopharyngeal evolution. Nature 520, 466–473 (2015).

10. Tzahor, E. Heart and craniofacial muscle development: A new developmental theme of distinct myogenic fields. Developmental Biology 327, 273–279 (2009).

11. Vyas, B., Nandkishore, N. & Sambasivan, R. Vertebrate cranial mesoderm: developmental trajectory and evolutionary origin. Cell. Mol. Life Sci. 77, 1933–1945 (2020).

12. Lescroart, F. et al. Clonal analysis reveals common lineage relationships between head muscles and second heart field derivatives in the mouse embryo. Development 137, 3269– 3279 (2010).

13. Lescroart, F. et al. Clonal analysis reveals a common origin between nonsomite-derived neck muscles and heart myocardium. Proc Natl Acad Sci USA 112, 1446–1451 (2015).

14. Lescroart, F. et al. Early lineage restriction in temporally distinct populations of Mesp1 progenitors during mammalian heart development. Nat Cell Biol 16, 829–840 (2014).

15. Stolfi, A. et al. Early Chordate Origins of the Vertebrate Second Heart Field. Science 329, 565–568 (2010).

16. Grimaldi, A. & Tajbakhsh, S. Diversity in cranial muscles: Origins and developmental programs. Current Opinion in Cell Biology 73, 110–116 (2021).

17. Tajbakhsh, S., Rocancourt, D., Cossu, G. & Buckingham, M. Redefining the Genetic Hierarchies Controlling Skeletal Myogenesis: Pax-3 and Myf-5 Act Upstream of MyoD. Cell 89, 127–138 (1997).

18. Theis, S. et al. The occipital lateral plate mesoderm is a novel source for vertebrate neck musculature. Development 137, 2961–2971 (2010).

19. Harel, I. et al. Distinct Origins and Genetic Programs of Head Muscle Satellite Cells. Developmental Cell 16, 822–832 (2009).

20. Harel, I. et al. Pharyngeal mesoderm regulatory network controls cardiac and head muscle morphogenesis. Proceedings of the National Academy of Sciences 109, 18839–18844 (2012).

21. Sambasivan, R. et al. Distinct Regulatory Cascades Govern Extraocular and Pharyngeal Arch Muscle Progenitor Cell Fates. Developmental Cell 16, 810–821 (2009).

22. Kelly, R. G., Jerome-Majewska, L. A. & Papaioannou, V. E. The del22q11.2 candidate gene Tbx1 regulates branchiomeric myogenesis. Human Molecular Genetics 13, 2829–2840 (2004).

23. Razy-Krajka, F. et al. Collier/OLF/EBF-Dependent Transcriptional Dynamics Control Pharyngeal Muscle Specification from Primed Cardiopharyngeal Progenitors. Developmental Cell 29, 263–276 (2014).

24. Wang, W. et al. A single-cell transcriptional roadmap for cardiopharyngeal fate diversification. Nat Cell Biol 21, 674–686 (2019).

25. Chabab, S. et al. Uncovering the Number and Clonal Dynamics of Mesp1 Progenitors during Heart Morphogenesis. Cell Reports 14, 1–10 (2016).

26. Murry, C. E. & Keller, G. Differentiation of Embryonic Stem Cells to Clinically Relevant Populations: Lessons from Embryonic Development. Cell 132, 661–680 (2008).

27. Kattman, S. J., Adler, E. D. & Keller, G. M. Specification of Multipotential Cardiovascular Progenitor Cells During Embryonic Stem Cell Differentiation and Embryonic Development. Trends in Cardiovascular Medicine 17, 240–246 (2007).

28. Chan, S. S.-K. et al. Development of Bipotent Cardiac/Skeletal Myogenic Progenitors from MESP1+ Mesoderm. Stem Cell Reports 6, 26–34 (2016).

29. Nandkishore, N., Vyas, B., Javali, A., Ghosh, S. & Sambasivan, R. Divergent early mesoderm specification underlies distinct head and trunk muscle programmes in vertebrates. Development 145, dev160945 (2018).

30. Turner, D. A. et al. Anteroposterior polarity and elongation in the absence of extraembryonic tissues and spatially localised signalling in *Gastruloids*, mammalian embryonic organoids. Development dev.150391 (2017) doi:10.1242/dev.150391.

31. van den Brink, S. C., et al. Symmetry breaking, germ layer specification and axial organisation in aggregates of mouse embryonic stem cells. Development 141, 4231–4242 (2014).

32. Beccari, L. et al. Multi-axial self-organization properties of mouse embryonic stem cells into gastruloids. Nature 562, 272–276 (2018).

33. Olmsted, Z. T. & Paluh, J. L. A combined human gastruloid model of cardiogenesis and neurogenesis. iScience 25, 104486 (2022).

34. Rossi, G. et al. Capturing Cardiogenesis in Gastruloids. Cell Stem Cell 28, 230–240.e6 (2021).

35. van den Brink, S. C., et al. Single-cell and spatial transcriptomics reveal somitogenesis in gastruloids. Nature 582, 405–409 (2020).

36. Veenvliet, J. V. et al. Mouse embryonic stem cells self-organize into trunk-like structures with neural tube and somites. Science (2020) doi:10.1126/science.aba4937.

37. Pijuan-Sala, B. et al. A single-cell molecular map of mouse gastrulation and early organogenesis. Nature 566, 490–495 (2019).

38. Song, M., et al. GATA4/5/6 family transcription factors are conserved determinants of cardiac versus pharyngeal mesoderm fate. 2020.12.01.406140 https://www.biorxiv.org/content/10.1101/2020.12.01.406140v1 (2020) doi:10.1101/2020.12.01.406140.

39. Swedlund, B. & Lescroart, F. Cardiopharyngeal Progenitor Specification: Multiple Roads to the Heart and Head Muscles. Cold Spring Harb Perspect Biol 12, a036731 (2020).

40. Tyser, R. C. V. et al. Characterization of a common progenitor pool of the epicardium and myocardium. Science 371, eabb2986 (2021).

41. Zhang, Q. et al. Unveiling Complexity and Multipotentiality of Early Heart Fields. Circ Res 129, 474–487 (2021).

42. Dupuis, L. E. et al. Altered versican cleavage in ADAMTS5 deficient mice; a novel etiology of myxomatous valve disease. Dev Biol 357, 152–164 (2011).

43. Gurdziel, K., Vogt, K. R., Walton, K. D., Schneider, G. K. & Gumucio, D. L. Transcriptome of the inner circular smooth muscle of the developing mouse intestine: Evidence for regulation of visceral smooth muscle genes by the hedgehog target gene, cJun. Developmental Dynamics 245, 614–626 (2016).

44. Liang, D. et al. Cellular and molecular landscape of mammalian sinoatrial node revealed by single-cell RNA sequencing. Nat Commun 12, 287 (2021).

45. Ola, R., Lefebvre, S., Braunewell, K.-H., Sainio, K. & Sariola, H. The expression of Visinin-like 1 during mouse embryonic development. Gene Expression Patterns 12, 53–62 (2012).

46. van Eif, V. et al. Transcriptome analysis of mouse and human sinoatrial node cells reveals a conserved genetic program. Development dev.173161 (2019) doi:10.1242/dev.173161.

47. DeLaughter, D. M. et al. Single-Cell Resolution of Temporal Gene Expression during Heart Development. Developmental Cell 39, 480–490 (2016).

48. Gonzalez, D. M. et al. Dissecting mechanisms of chamber-specific cardiac differentiation and its perturbation following retinoic acid exposure. Development dev.200557 (2022) doi:10.1242/dev.200557.

49. Li, G. et al. Transcriptomic Profiling Maps Anatomically Patterned Subpopulations among Single Embryonic Cardiac Cells. Developmental Cell 39, 491–507 (2016).

50. Verzi, M. P., McCulley, D. J., De Val, S., Dodou, E. & Black, B. L. The right ventricle, outflow tract, and ventricular septum comprise a restricted expression domain within the secondary/anterior heart field. Developmental Biology 287, 134–145 (2005).

51. Adachi, N., Bilio, M., Baldini, A. & Kelly, R. G. Cardiopharyngeal mesoderm origins of musculoskeletal and connective tissues in the mammalian pharynx. Development 147, dev185256 (2020).

52. De Bono, C. et al. T-box genes and retinoic acid signaling regulate the segregation of arterial and venous pole progenitor cells in the murine second heart field. Human Molecular Genetics 27, 3747–3760 (2018).

53. Comai, G. et al. A distinct cardiopharyngeal mesoderm genetic hierarchy establishes antero-posterior patterning of esophagus striated muscle. eLife 8, e47460 (2019).

54. Prunotto, C. et al. Analysis of Mlc-lacZ Met mutants highlights the essential function of Met for migratory precursors of hypaxial muscles and reveals a role for Met in the development of hyoid arch-derived facial muscles. Developmental Dynamics 231, 582–591 (2004).

55. Ding, J., Sharon, N. & Bar-Joseph, Z. Temporal modelling using single-cell transcriptomics. Nat Rev Genet 23, 355–368 (2022).

56. Lange, M. et al. CellRank for directed single-cell fate mapping. Nat Methods 19, 159– 170 (2022).

57. Farrell, J. A. et al. Single-cell reconstruction of developmental trajectories during zebrafish embryogenesis. Science 360, eaar3131 (2018).

58. Carmona, R. et al. The embryonic epicardium: an essential element of cardiac development. Journal of Cellular and Molecular Medicine 14, 2066–2072 (2010).

59. Wagner, D. E. & Klein, A. M. Lineage tracing meets single-cell omics: opportunities and challenges. Nat Rev Genet 21, 410–427 (2020).

60. Ng, S. Y., Wong, C. K. & Tsang, S. Y. Differential gene expressions in atrial and ventricular myocytes: insights into the road of applying embryonic stem cell-derived cardiomyocytes for future therapies. American Journal of Physiology-Cell Physiology 299, C1234–C1249 (2010).

61. Schmidt, C. et al. Multi-chamber cardioids unravel human heart development and cardiac defects. 2022.07.14.499699 Preprint at https://doi.org/10.1101/2022.07.14.499699 (2022).

62. Kidokoro, H., Yonei-Tamura, S., Tamura, K., Schoenwolf, G. C. & Saijoh, Y. The heart tube forms and elongates through dynamic cell rearrangement coordinated with foregut extension. Development dev.152488 (2018) doi:10.1242/dev.152488.

63. Nascone, N. & Mercola, M. An inductive role for the endoderm in Xenopus cardiogenesis. Development 121, 515–523 (1995).

64. Etchevers, H. C., Dupin, E. & Le Douarin, N. M. The diverse neural crest: from embryology to human pathology. Development 146, dev169821 (2019).

65. Keyte, A. & Hutson, M. R. The neural crest in cardiac congenital anomalies. Differentiation 84, 25–40 (2012).

66. Le Douarin, N. M., Creuzet, S., Couly, G. & Dupin, E. Neural crest cell plasticity and its limits. Development 131, 4637–4650 (2004).

67. Rinon, A. et al. Cranial neural crest cells regulate head muscle patterning and differentiation during vertebrate embryogenesis. Development 134, 3065–3075 (2007).

68. Gopalakrishnan, S. et al. A Cranial Mesoderm Origin for Esophagus Striated Muscles. Developmental Cell 34, 694–704 (2015).

69. Sanaki-Matsumiya, M. et al. Periodic formation of epithelial somites from human pluripotent stem cells. Nat Commun 13, 2325 (2022).

70. Budjan, C. et al. Paraxial mesoderm organoids model development of human somites. eLife 11, e68925 (2022).

71. Miao, Y. et al. Reconstruction and deconstruction of human somitogenesis in vitro. Nature 614, 500–508 (2023).

72. Madisen, L. et al. A robust and high-throughput Cre reporting and characterization system for the whole mouse brain. Nat Neurosci 13, 133–140 (2010).

73. Iacovino, M., Roth, M. E. & Kyba, M. Rapid Genetic Modification of Mouse Embryonic Stem Cells by Inducible Cassette Exchange Recombination. in Gene Function Analysis (ed. Ochs, M. F.) 339–351 (Humana Press, 2014). doi:10.1007/978-1-62703-721-1_16.

74. Hao, Y. et al. Integrated analysis of multimodal single-cell data. Cell 184, 3573–3587.e29 (2021).

75. McInnes, L., Healy, J. & Melville, J. UMAP: Uniform Manifold Approximation and Projection for Dimension Reduction. Preprint at https://doi.org/10.48550/arXiv.1802.03426 (2020).

76. Traag, V. A., Waltman, L. & van Eck, N. J. From Louvain to Leiden: guaranteeing well-connected communities. Sci Rep 9, 5233 (2019).

77. McGinnis, C. S., Murrow, L. M. & Gartner, Z. J. DoubletFinder: Doublet Detection in Single-Cell RNA Sequencing Data Using Artificial Nearest Neighbors. Cell Syst 8, 329–337.e4 (2019).

78. Virshup, I., Rybakov, S., Theis, F. J., Angerer, P. & Wolf, F. A. anndata: Annotated data. 2021.12.16.473007 Preprint at https://doi.org/10.1101/2021.12.16.473007 (2021).

79. Wolf, F. A., Angerer, P. & Theis, F. J. SCANPY: large-scale single-cell gene expression data analysis. Genome Biol 19, 15 (2018).

80. Schiebinger, G. et al. Optimal-Transport Analysis of Single-Cell Gene Expression Identifies Developmental Trajectories in Reprogramming. Cell 176, 928–943.e22 (2019).

81. Reuter, B., Weber, M., Fackeldey, K., Röblitz, S. & Garcia, M. E. Generalized Markov State Modeling Method for Nonequilibrium Biomolecular Dynamics: Exemplified on Amyloid β Conformational Dynamics Driven by an Oscillating Electric Field. J Chem Theory Comput 14, 3579–3594 (2018).

82. Brandts, J. Matlab code for sorting real Schur forms. Numerical Linear Algebra with Applications 9, 249–261 (2002).

