## Supplementary materials for "Cardiopharyngeal Mesoderm specification into cardiac and skeletal muscle lineages in gastruloids"

### Supplementary figure legends

**Supplementary Figure S1. Similar expression kinetics with 2 different cell lines.** Expression profiles of *Mesp1* (**a**), *Isl1* (**b**)and *Tcf21* (**c**) throughout the culture of gastruloids with Zx1 (blue) and R1 (red) cell lines as measured by quantitative RT-PCR. Results are normalized on the expression of *TBP*. Fold changes are represented over expression day 0. (Mean with standard error of mean (SEM).

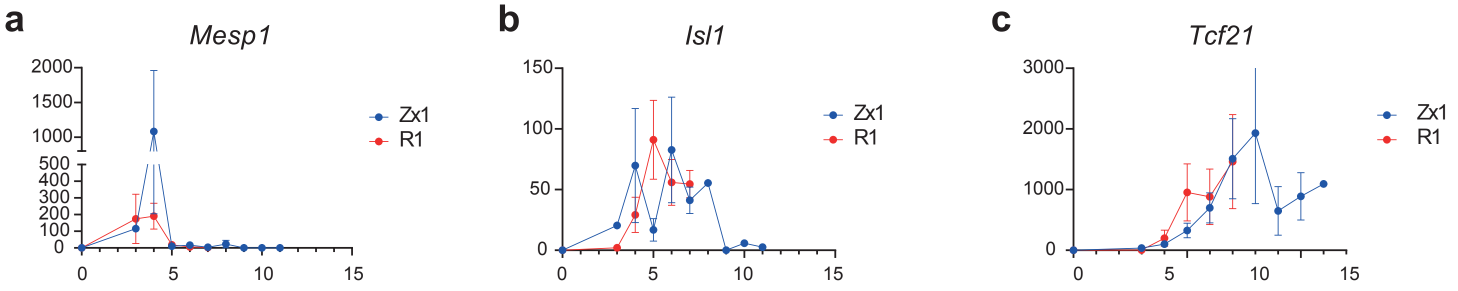


**Supplementary Figure S1**

Argiro et al.

**Supplementary Figure S2. Analysis of single-cell gastruloids data integrated with the embryonic atlas.** UMAP representations of the mouse embryonic cell atlas integrated with gastruloids' single cell datasets at day 4 (**a-c**), day 5 (**d-f**), day 6 (**g-i**) and day 11 (**j-l**). Embryonic stages are represented in **a**, **d**, **g** and **j**. Legends are found below **j**. Cell types are represented for the cells of the embryonic atlas in **b**, **e**, **h** and **k**, while cell types annotation transfer from the atlas in gastruloids' cells are shown in **c**, **f**, **i** and **l**. Legends are found below **k-l**.

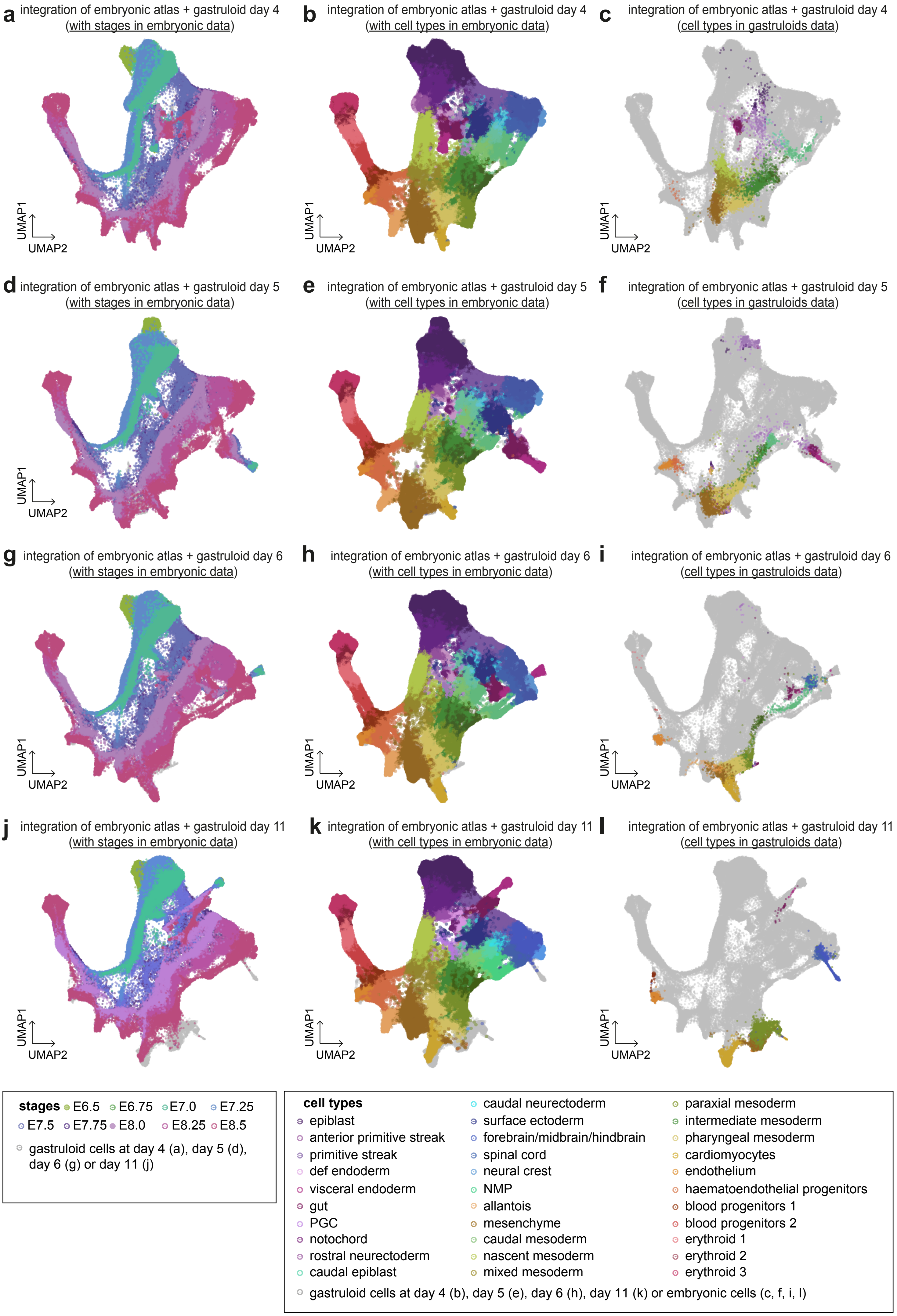


**Supplementary Figure S2**

Argiro et al.

**Supplementary Figure S3. Predictive cell types from the embryonic cell atlas in gastruloids at day 4.** UMAP representations of gastruloids' single cell datasets at day 4 with predicted cell types from the embryonic cell atlas.

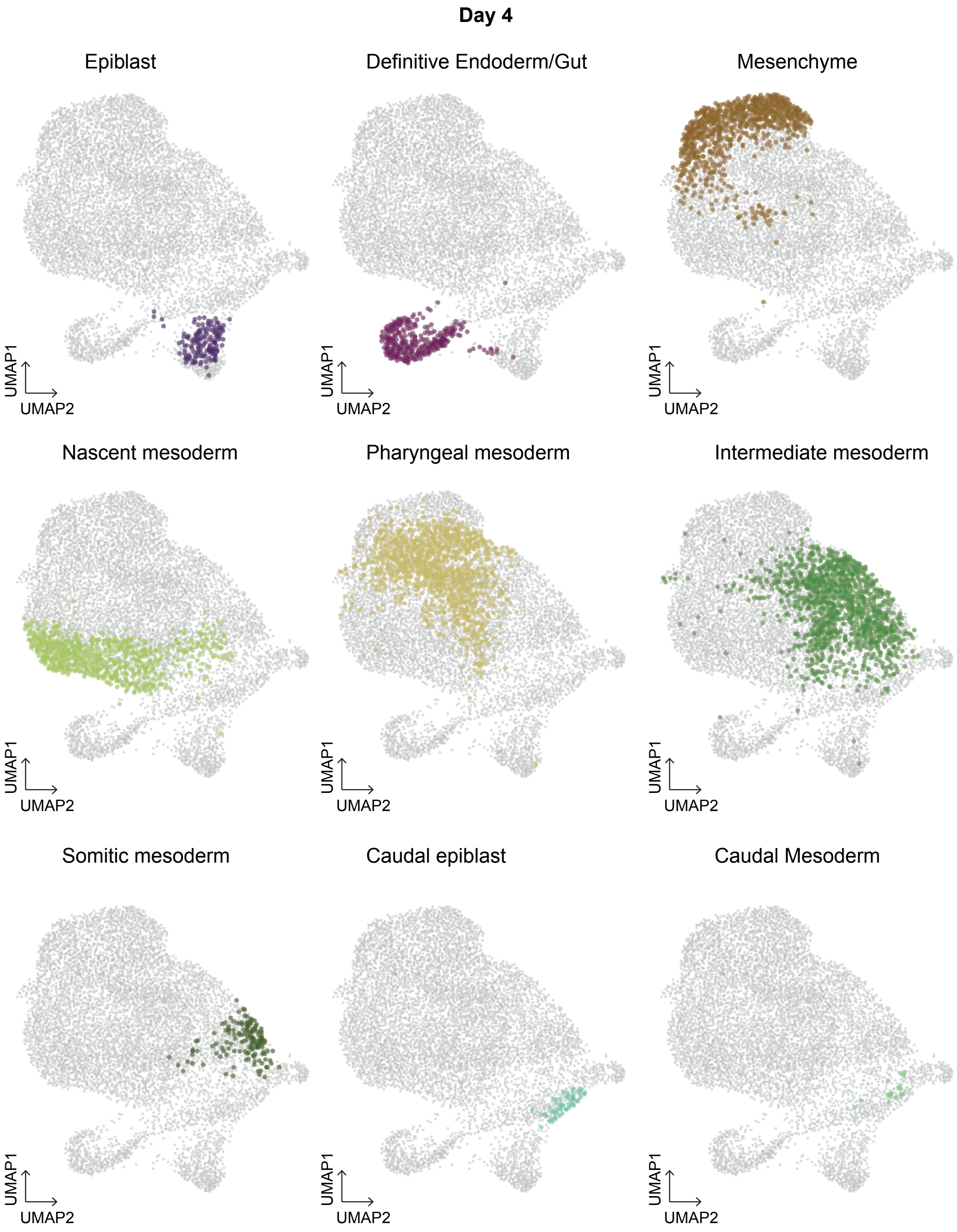


**Supplementary Figure S3**

Argiro et al.

**Supplementary Figure S4. Expression of key markers in single-cell RNAseq of gastruloids at day 4.** FeaturePlots showing expression of *Gata6*, *Hand1*, *T*, and *Mesp2* in gastruloids at day 4. Scale bars represent expression levels. Numbers represent Leiden clusters.

Day 4

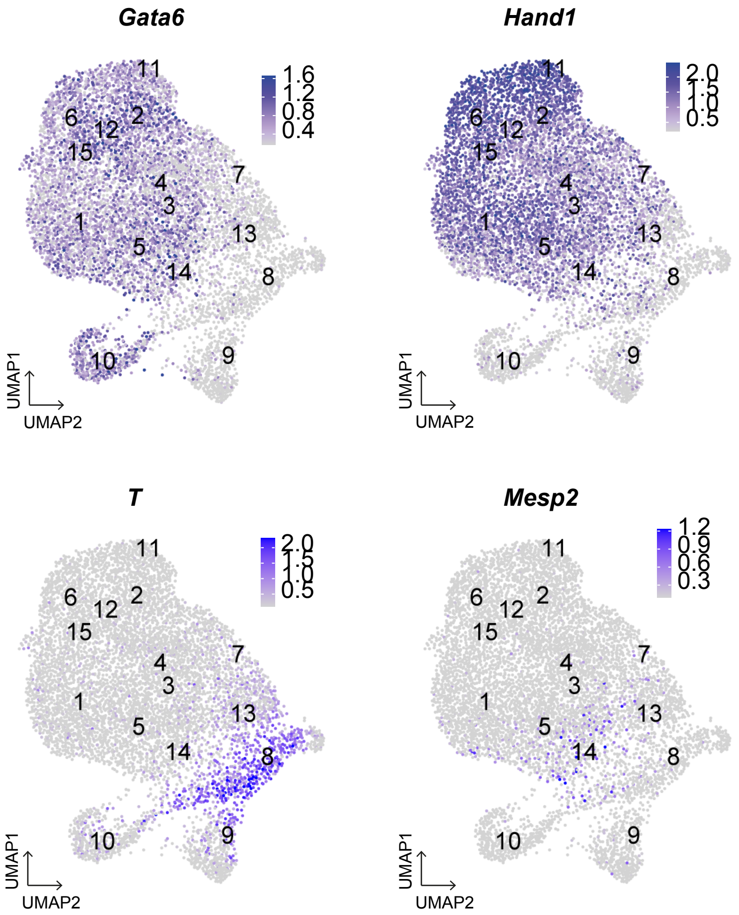


**Supplementary Figure S4**

Argiro et al.

**Supplementary Figure S5. Predictive cell types from the embryonic cell atlas in gastruloids at day 5.** UMAP representations of gastruloids' single cell datasets at day 5 with predicted cell types from the embryonic cell atlas.

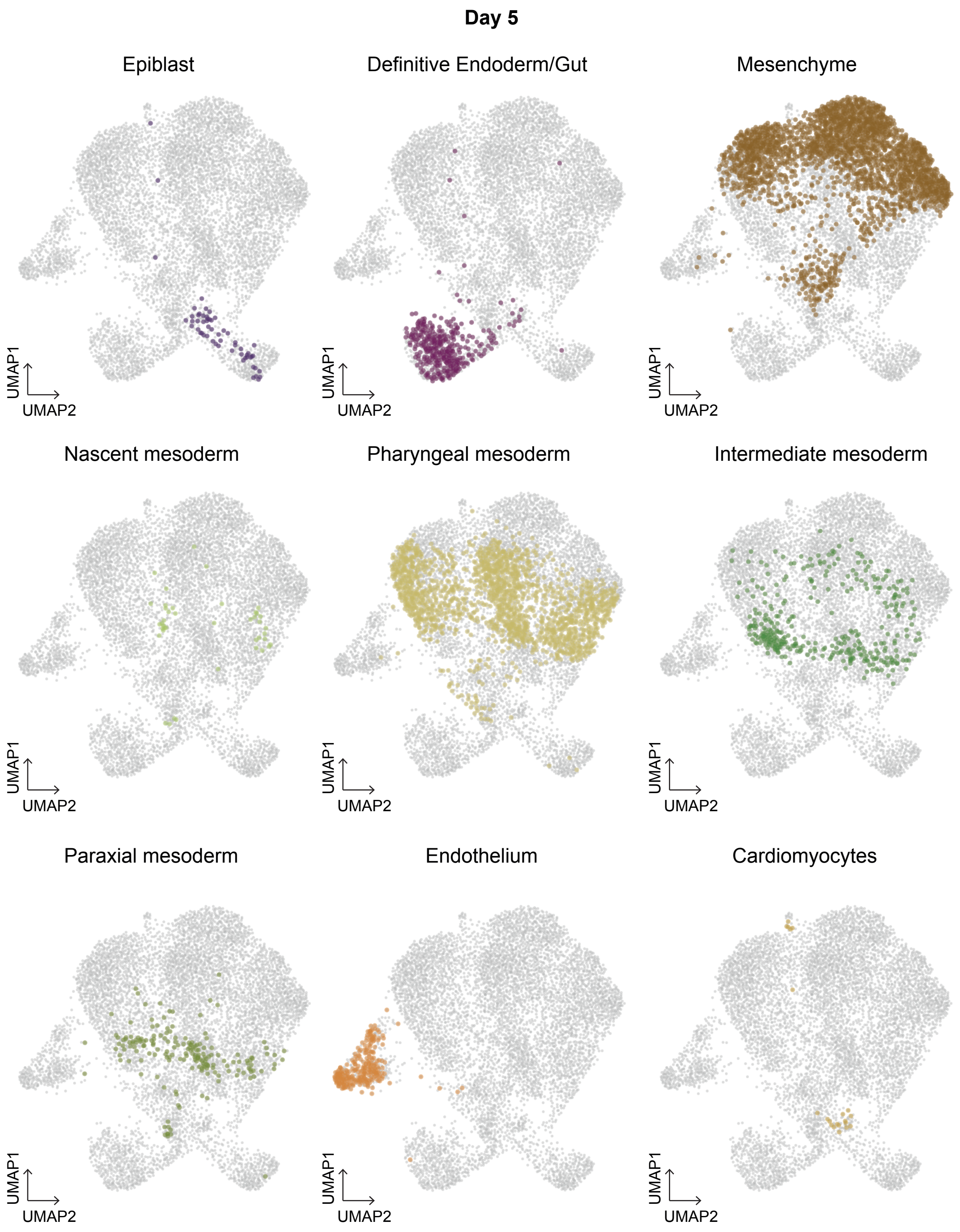


**Supplementary Figure S5**

Argiro et al.

**Supplementary Figure S6. Expression of key markers in single-cell RNAseq of gastruloids at day 5.** FeaturePlots showing expression of *Mesp1, Gata6*, *Hand1* and *Tbx1* in gastruloids at day 5. Scale bars represent expression levels. Numbers represent Leiden clusters.

**Day 5**

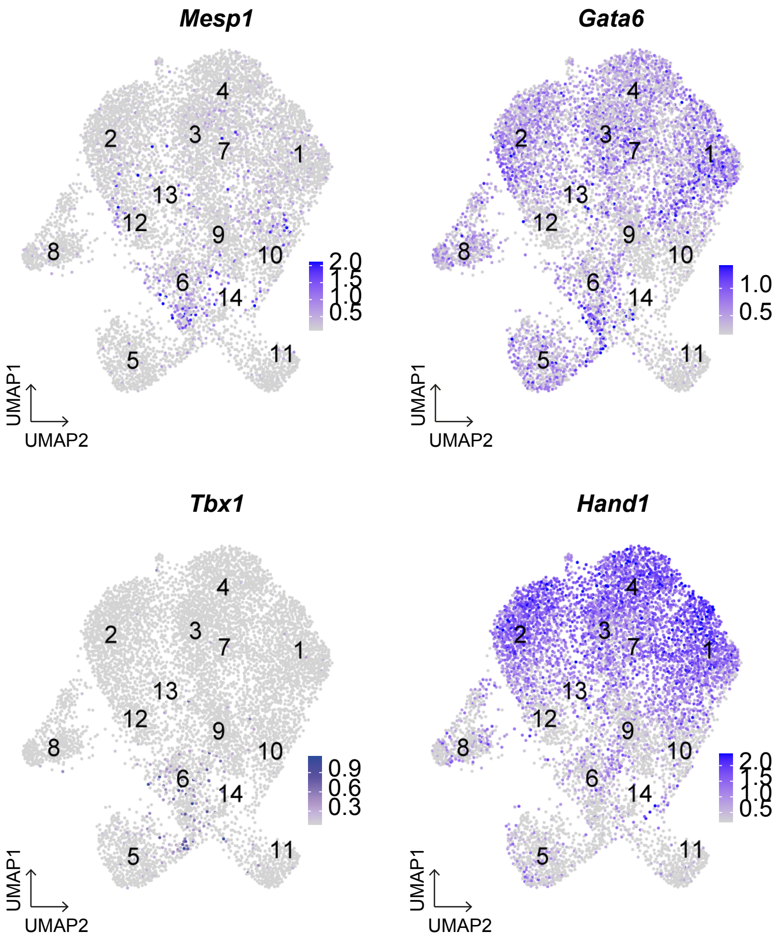


**Supplementary Figure S6**

Argiro et al.

**Supplementary Figure S7. Predictive cell types from the embryonic cell atlas in gastruloids at day 6.** UMAP representations of gastruloids' single cell datasets at day 6 with predicted cell types from the embryonic cell atlas.

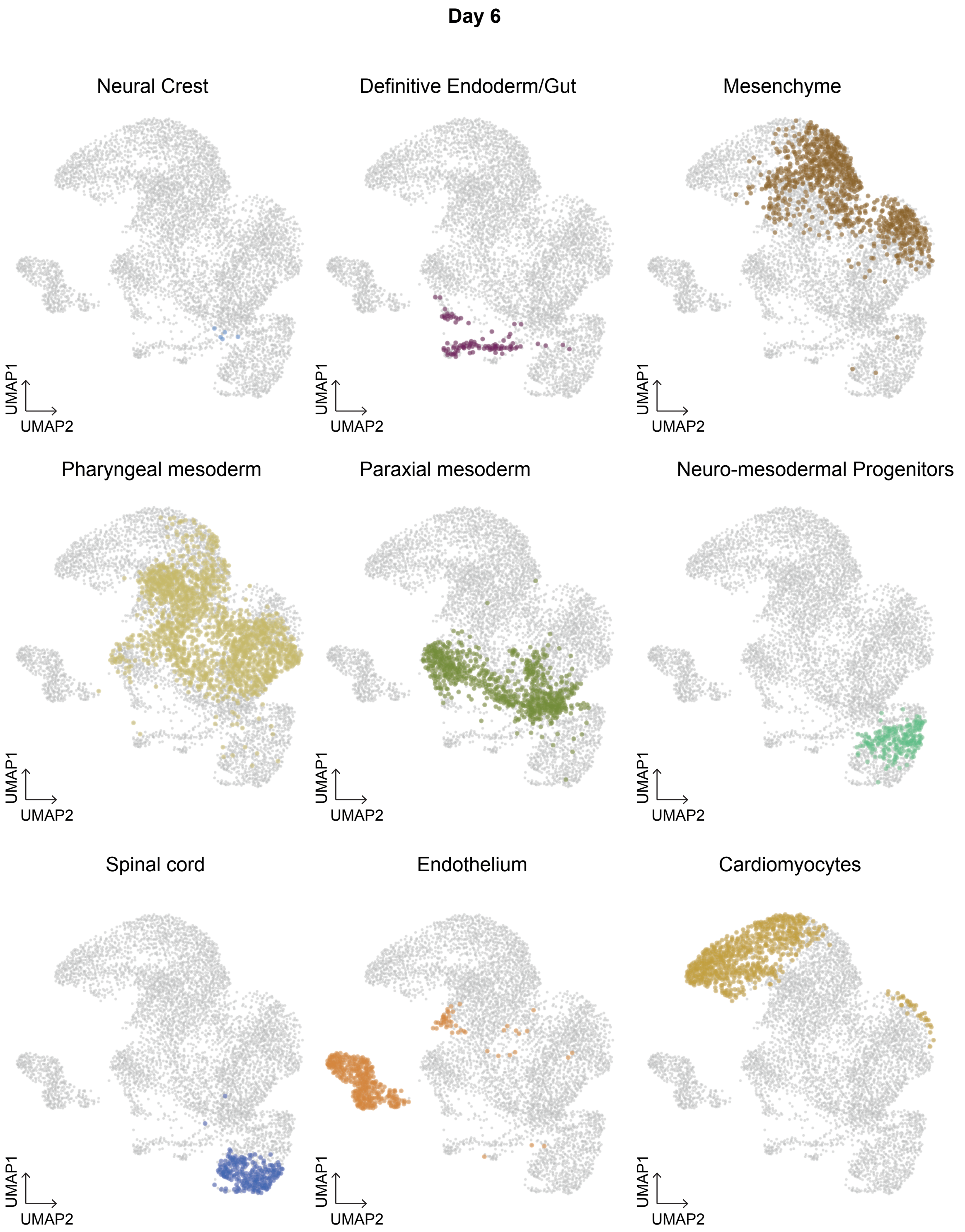


**Supplementary Figure S7**

**Supplementary Figure S8. Expression of key markers in single-cell RNAseq of gastruloids at day 6.** FeaturePlots showing expression of *Gata4*, *Gata6*, *Hand1*, *Mab21l2*, *Mef2c*, *Isl1*, *Pax3* and *Tbx1* in gastruloids at day 6. Scale bars represent expression levels. Numbers represent Leiden clusters.

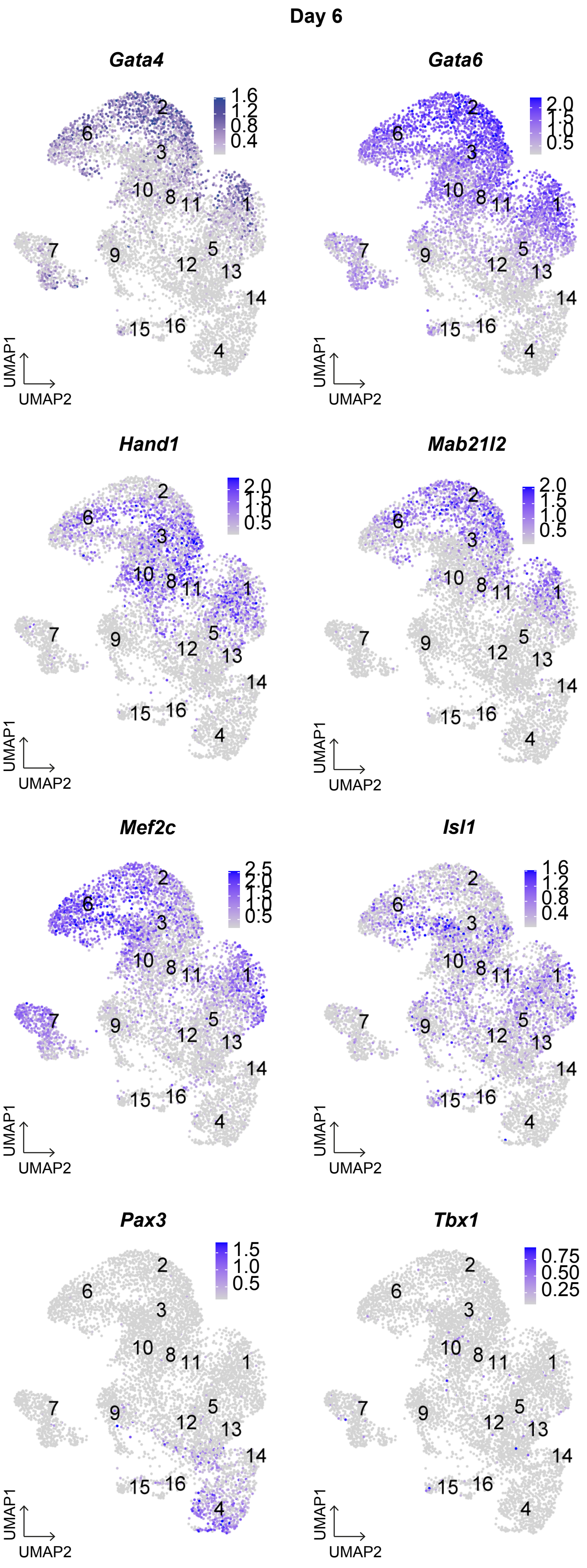


**Supplementary Figure S8**

Argiro et al.

**Supplementary Figure S9. Expression of key markers in single-cell RNAseq of gastruloids at day 11.** FeaturePlots showing expression of *Tbx1*, *Isl1*, *Tcf21*, *Mab21l2*, *Wt1*, *Tbx18*, *Pax3* and *Meox1* in gastruloids at day 6. Scale bars represent expression levels.

**Day 11**

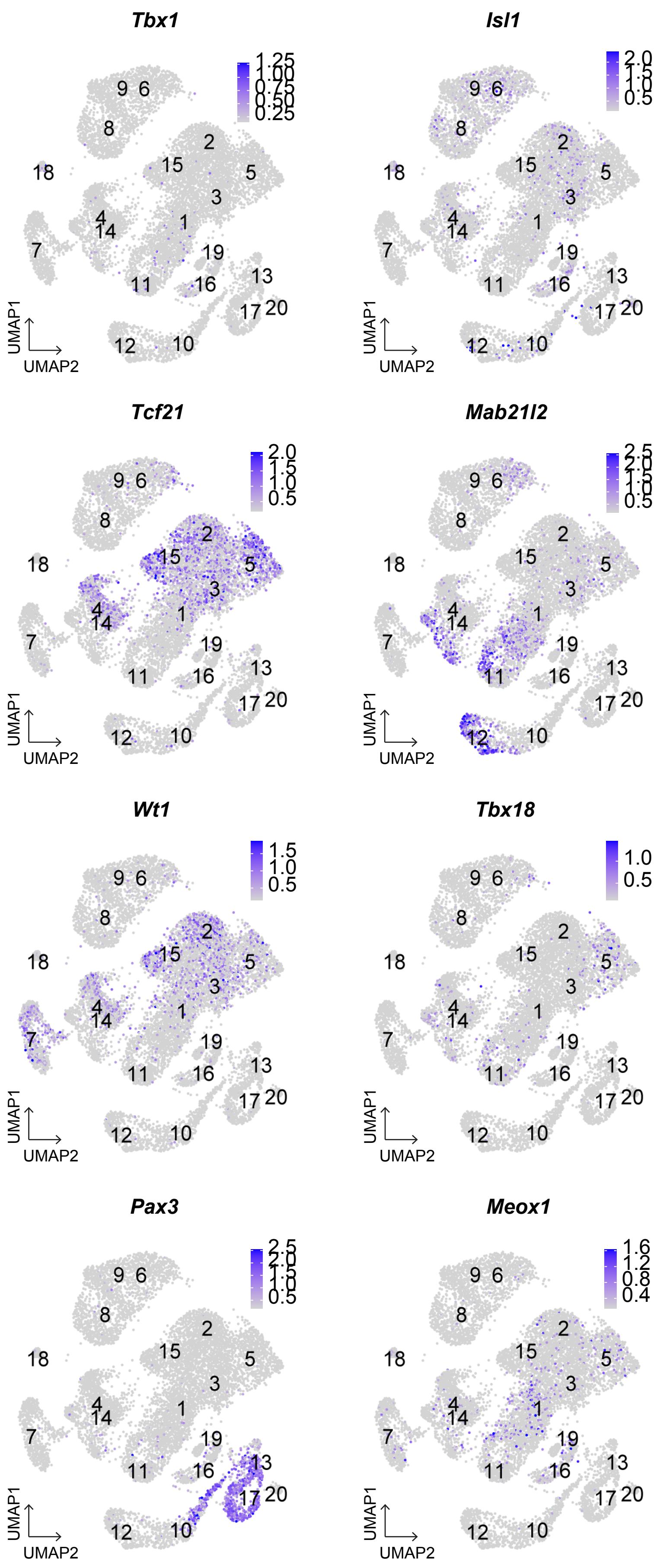


**Supplementary Figure S9**

**Supplementary Figure S10. Expression of key markers across the URD trajectory.** Expression of *Gata6, Hand1, Mab21l2, Nkx2-5, Tbx18, Wt1, Lhx2, Ebf3, Tbx1, MyoR, Myf5 and Pax3* across URD trajectory. Scale bars represent expression levels.

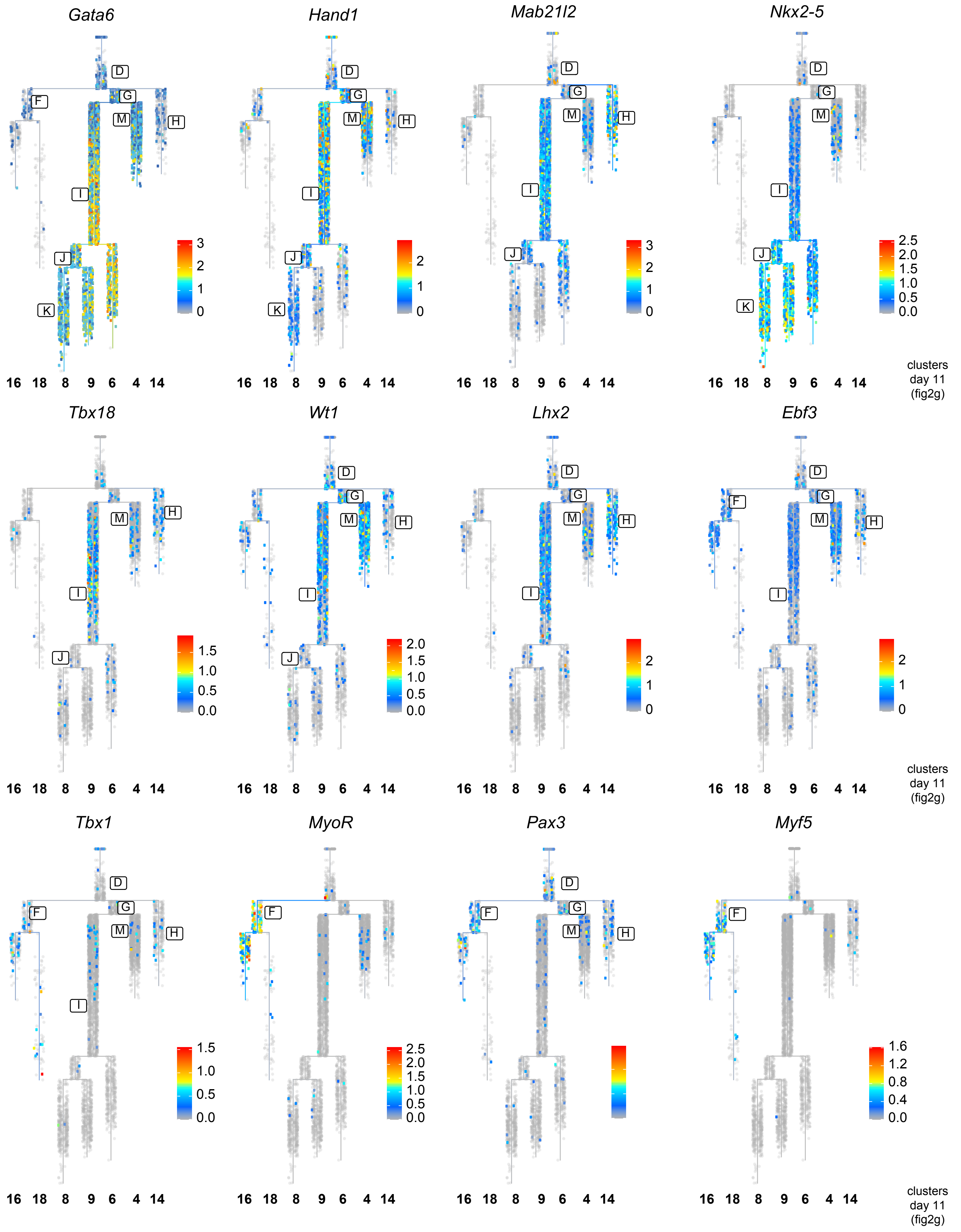


**Supplementary Figure S10**

Argiro et al.

**<u>Supplementary Table S1:</u> Differential gene expression analysis at day 4**

| Cluster | Gene | avg_logFC | pct.1 | pct.2 | p_val | p_val_adj | most expressed celltype | pct.celltype |
|---|---|---|---|---|---|---|---|---|
| 1 | Eomes | 1.44062310567834 | 0.91 | 0.225 | 0 | 0 | Mixed mesoderm | 52.83 |
| 1 | Mixl1 | 0.835397458873914 | 0.643 | 0.145 | 1.35513753715201e-288 | 2.61568647421081e-284 | Mixed mesoderm | 52.83 |
| 1 | Kdr | 0.851842228955024 | 0.902 | 0.397 | 1.10245003775088e-273 | 2.12794906286675e-269 | Mixed mesoderm | 52.83 |
| 1 | Fn1 | 0.889860122228518 | 0.999 | 0.935 | 1.42493860350167e-225 | 2.75041649247892e-221 | Mixed mesoderm | 52.83 |
| 1 | S100a10 | 1.0782240936129 | 0.985 | 0.771 | 3.03478034316966e-209 | 5.85773301838608e-205 | Mixed mesoderm | 52.83 |
| 1 | Tdgf1 | 0.823745269451282 | 0.732 | 0.257 | 1.87010618980983e-187 | 3.60967896757094e-183 | Mixed mesoderm | 52.83 |
| 1 | Asb4 | 0.996503960749489 | 0.881 | 0.438 | 2.46672342607014e-186 | 4.76126955700058e-182 | Mixed mesoderm | 52.83 |
| 1 | Cyp26a1 | 0.722592407000632 | 0.762 | 0.336 | 8.76373480987582e-176 | 1.69157609300223e-171 | Mixed mesoderm | 52.83 |
| 1 | Vstm2b | 0.733228517196797 | 0.725 | 0.333 | 1.1650952008005e-170 | 2.24886675658513e-166 | Mixed mesoderm | 52.83 |
| 1 | Myl7 | 0.940052511423953 | 0.719 | 0.334 | 2.23394341905128e-162 | 4.31195758745279e-158 | Mixed mesoderm | 52.83 |
| 1 | Smad1 | 0.55770988298489 | 0.917 | 0.681 | 1.18548399920086e-154 | 2.28822121525749e-150 | Mixed mesoderm | 52.83 |
| 1 | Olfm1 | 0.548230328789921 | 0.783 | 0.43 | 7.49776022993039e-149 | 1.44721767958116e-144 | Mixed mesoderm | 52.83 |
| 1 | Gas1 | 0.558610223354859 | 0.838 | 0.509 | 1.30451109193454e-145 | 2.51796730965206e-141 | Mixed mesoderm | 52.83 |
| 1 | Mesp1 | 0.862266795151961 | 0.929 | 0.62 | 1.0455006157652e-141 | 2.01802528854999e-137 | Mixed mesoderm | 52.83 |
| 1 | Hmga2 | 0.573807788046865 | 0.999 | 0.924 | 1.31906564042568e-123 | 2.54606049914964e-119 | Mixed mesoderm | 52.83 |
| 1 | Vldlr | 0.519285271504902 | 0.767 | 0.494 | 3.92603482551782e-101 | 7.57803242021449e-97 | Mixed mesoderm | 52.83 |
| 1 | Pmp22 | 0.547744356097404 | 0.911 | 0.593 | 5.28924950270053e-99 | 1.02093093901126e-94 | Mixed mesoderm | 52.83 |
| 1 | Tbx3 | 0.560522706469912 | 0.889 | 0.668 | 4.24994361482167e-97 | 8.20324116532879e-93 | Mixed mesoderm | 52.83 |
| 1 | Phlda2 | 0.646191090995447 | 0.977 | 0.835 | 2.24514407423875e-77 | 4.33357709209563e-73 | Mixed mesoderm | 52.83 |
| 1 | Lhx1 | 0.737615187598106 | 0.8 | 0.622 | 2.04279393396095e-62 | 3.94300085133143e-58 | Mixed mesoderm | 52.83 |
| 2 | Twist1 | 1.11763736518495 | 0.951 | 0.631 | 5.1019151307526e-184 | 9.84771658537866e-180 | Pharyngeal mesoderm | 75.33 |
| 2 | Csrp2 | 0.740372667954095 | 0.999 | 0.944 | 1.92202061261125e-137 | 3.70988418646223e-133 | Pharyngeal mesoderm | 75.33 |
| 2 | Hand2 | 0.677529840869601 | 0.924 | 0.471 | 1.99897471707049e-159 | 3.85842099888945e-155 | Pharyngeal mesoderm | 75.33 |
| 2 | Hmga2 | 0.645317199314846 | 1 | 0.927 | 2.42888364931247e-129 | 4.68823121990292e-125 | Pharyngeal mesoderm | 75.33 |
| 2 | Igf2 | 0.636653076519623 | 0.97 | 0.667 | 1.00903071096234e-127 | 1.94763107829951e-123 | Pharyngeal mesoderm | 75.33 |
| 2 | Cgnl1 | 0.581903989744196 | 0.952 | 0.58 | 5.49613252131284e-133 | 1.06086349926381e-128 | Pharyngeal mesoderm | 75.33 |
| 2 | Prrx2 | 0.543034681694713 | 0.878 | 0.485 | 5.80746456303852e-109 | 1.12095680995769e-104 | Pharyngeal mesoderm | 75.33 |
| 2 | Gata6 | 0.53668087449583 | 0.933 | 0.62 | 4.55770526079371e-93 | 8.79728269438402e-89 | Pharyngeal mesoderm | 75.33 |
| 2 | 1810058I24 | 0.531901975266017 | 0.999 | 0.969 | 2.09685830177094e-128 | 4.04735589407827e-124 | Pharyngeal mesoderm | 75.33 |
| 2 | Prrx1 | 0.52325218536575 | 0.776 | 0.246 | 7.53060984060432e-188 | 1.45355831143345e-183 | Pharyngeal mesoderm | 75.33 |
| 2 | Hist1h1b | 0.517730638248395 | 0.896 | 0.719 | 4.49344698291694e-41 | 8.67325136642629e-37 | Pharyngeal mesoderm | 75.33 |
| 2 | Cd24a | 0.501744335928284 | 0.997 | 0.901 | 3.38643330293132e-108 | 6.53649356131802e-104 | Pharyngeal mesoderm | 75.33 |
| 2 | Hoxb1 | 0.498786582874071 | 0.941 | 0.55 | 1.63250479022774e-99 | 3.15106074609759e-95 | Pharyngeal mesoderm | 75.33 |
| 2 | Slc38a4 | 0.495270424734017 | 0.973 | 0.671 | 1.02887209472731e-105 | 1.98592891724265e-101 | Pharyngeal mesoderm | 75.33 |
| 2 | Fth1 | 0.492139436345933 | 1 | 1 | 4.1104497539932e-109 | 7.93399011515768e-105 | Pharyngeal mesoderm | 75.33 |
| 2 | Uaca | 0.470054910868221 | 0.909 | 0.458 | 1.3689493279971e-131 | 2.64234599290001e-127 | Pharyngeal mesoderm | 75.33 |
| 2 | Mest | 0.46736111690169 | 0.774 | 0.517 | 7.6591867730802e-50 | 1.47837623093994e-45 | Pharyngeal mesoderm | 75.33 |
| 2 | Tuba1a | 0.462060438357908 | 0.997 | 0.948 | 2.42161347256734e-80 | 4.67419832474947e-76 | Pharyngeal mesoderm | 75.33 |
| 2 | Nrp1 | 0.449395038920022 | 0.96 | 0.604 | 1.50684343715875e-97 | 2.90850920240382e-93 | Pharyngeal mesoderm | 75.33 |
| 2 | Id2 | 0.443754618712582 | 0.927 | 0.687 | 1.34250002410478e-63 | 2.59129354652705e-59 | Pharyngeal mesoderm | 75.33 |
| 3 | Ube2c | 1.03032167846094 | 1 | 0.843 | 1.364349083498e-151 | 2.63346660096784e-147 | Mixed mesoderm | 35.55 |
| 3 | Cenpf | 0.99931868855719 | 0.997 | 0.835 | 4.19363110772496e-157 | 8.09454676413072e-153 | Mixed mesoderm | 35.55 |
| 3 | Mesp1 | 0.881872799075859 | 0.93 | 0.633 | 7.53458045582092e-93 | 1.45432471958255e-88 | Mixed mesoderm | 35.55 |
| 3 | Top2a | 0.823963221938589 | 0.995 | 0.867 | 3.49240712641985e-142 | 6.74104423541559e-138 | Mixed mesoderm | 35.55 |
| 3 | Nusap1 | 0.81425721860379 | 0.942 | 0.536 | 1.24725420254292e-149 | 2.40745006174835e-145 | Mixed mesoderm | 35.55 |
| 3 | Prc1 | 0.802614010230456 | 0.955 | 0.551 | 3.93195224188799e-152 | 7.5894542172922e-148 | Mixed mesoderm | 35.55 |
| 3 | Hmmr | 0.797011299206442 | 0.989 | 0.654 | 2.5119072018882e-160 | 4.8484832810846e-156 | Mixed mesoderm | 35.55 |
| 3 | Arl6ip1 | 0.760530800342425 | 0.998 | 0.896 | 3.70451866948879e-115 | 7.15046193584726e-111 | Mixed mesoderm | 35.55 |
| 3 | Egln1 | 0.747700466937284 | 0.984 | 0.805 | 3.05202682492437e-123 | 5.89102217746901e-119 | Mixed mesoderm | 35.55 |
| 3 | Cdk1 | 0.70342513098451 | 0.992 | 0.825 | 4.85187528925093e-126 | 9.36508968331214e-122 | Mixed mesoderm | 35.55 |
| 3 | Tpx2 | 0.657623623383024 | 0.989 | 0.766 | 1.17958041303628e-120 | 2.27682611324262e-116 | Mixed mesoderm | 35.55 |
| 3 | Cenpe | 0.654466064800494 | 0.979 | 0.728 | 6.08962955934413e-118 | 1.1754202975446e-113 | Mixed mesoderm | 35.55 |
| 3 | Mki67 | 0.645509130300431 | 0.981 | 0.775 | 9.0967474219457e-104 | 1.75585418738396e-99 | Mixed mesoderm | 35.55 |
| 3 | Ccnb1 | 0.619358413830368 | 0.984 | 0.749 | 1.69914719584139e-116 | 3.27969391741304e-112 | Mixed mesoderm | 35.55 |
| 3 | Higd1a | 0.608447900248967 | 1 | 0.987 | 1.00748923526064e-102 | 1.94465572190009e-98 | Mixed mesoderm | 35.55 |
| 3 | 2410006H1 | 0.592165925181121 | 1 | 0.994 | 9.45277555254604e-90 | 1.82457473715244e-85 | Mixed mesoderm | 35.55 |
| 3 | Pfkp | 0.563810817904391 | 0.825 | 0.483 | 6.66846782662578e-98 | 1.28714765989531e-93 | Mixed mesoderm | 35.55 |
| 3 | Bub3 | 0.560131693380085 | 0.987 | 0.871 | 8.22389759263096e-108 | 1.58737671332963e-103 | Mixed mesoderm | 35.55 |
| 3 | Fam162a | 0.558234264357764 | 1 | 0.998 | 1.95115493446136e-113 | 3.76611925449732e-109 | Mixed mesoderm | 35.55 |
| 3 | Mis18bp1 | 0.556711151271175 | 0.94 | 0.619 | 2.1819087976244e-112 | 4.21152036117462e-108 | Mixed mesoderm | 35.55 |
| 4 | Mest | 0.920263549728098 | 0.662 | 0.532 | 2.53057770698222e-37 | 4.88452109001709e-33 | Intermediate mesoderm | 48.08 |
| 4 | Crabp1 | 0.753039727481981 | 0.939 | 0.731 | 4.65939584285005e-86 | 8.99356585586916e-82 | Intermediate mesoderm | 48.08 |
| 4 | Hotairm1 | 0.730250133007812 | 0.777 | 0.4 | 2.1990352621688e-123 | 4.24457786303822e-119 | Intermediate mesoderm | 48.08 |
| 4 | Twist1 | 0.712915826381994 | 0.848 | 0.646 | 6.313737214927e-73 | 1.21867755722521e-68 | Intermediate mesoderm | 48.08 |
| 4 | Aldh1a2 | 0.672205561929295 | 0.697 | 0.39 | 5.53559815181628e-73 | 1.06848115526358e-68 | Intermediate mesoderm | 48.08 |
| 4 | Grb10 | 0.669458367233713 | 0.902 | 0.755 | 3.17268478754482e-73 | 6.12391617691901e-69 | Intermediate mesoderm | 48.08 |
| 4 | Id2 | 0.663345293954189 | 0.88 | 0.696 | 5.01601186078262e-64 | 9.68190609368262e-60 | Intermediate mesoderm | 48.08 |
| 4 | Osr1 | 0.605685680909679 | 0.538 | 0.164 | 6.45088952419274e-122 | 1.24515069595968e-117 | Intermediate mesoderm | 48.08 |
| 4 | Prrx1 | 0.584135715416316 | 0.615 | 0.27 | 4.19360754542239e-91 | 8.0945012841743e-87 | Intermediate mesoderm | 48.08 |
| 4 | Tgfb2 | 0.560755240672246 | 0.735 | 0.458 | 1.32213937590822e-70 | 2.55199342337805e-66 | Intermediate mesoderm | 48.08 |
| 4 | Mdk | 0.550654663698621 | 1 | 0.996 | 4.46526470709458e-115 | 8.61885393763396e-111 | Intermediate mesoderm | 48.08 |
| 4 | Gm45889 | 0.547494302580283 | 0.826 | 0.612 | 2.82155985178222e-63 | 5.44617482591003e-59 | Intermediate mesoderm | 48.08 |
| 4 | Meis2 | 0.529177445824579 | 0.958 | 0.742 | 2.22723270831292e-78 | 4.2990045735856e-74 | Intermediate mesoderm | 48.08 |
| 4 | Prrx2 | 0.524379875068127 | 0.754 | 0.503 | 6.93709709490764e-61 | 1.33899848125907e-56 | Intermediate mesoderm | 48.08 |
| 4 | Hoxb1 | 0.406671782639628 | 0.836 | 0.567 | 1.9904498539176e-50 | 3.84196630803176e-46 | Intermediate mesoderm | 48.08 |
| 4 | Cd24a | 0.404188704875072 | 0.963 | 0.906 | 7.48361084306853e-40 | 1.44448656492909e-35 | Intermediate mesoderm | 48.08 |
| 4 | Tpm1 | 0.402844715620192 | 0.986 | 0.967 | 3.10363658926515e-28 | 5.99063934459959e-24 | Intermediate mesoderm | 48.08 |
| 4 | Igf2 | 0.391101694678958 | 0.821 | 0.686 | 4.94227705653587e-27 | 9.53958317452553e-23 | Intermediate mesoderm | 48.08 |
| 4 | Serpinh1 | 0.38548591415978 | 0.96 | 0.908 | 3.91643857418891e-40 | 7.55950973589943e-36 | Intermediate mesoderm | 48.08 |
| 4 | Cdx2 | 0.38227009427551 | 0.704 | 0.422 | 1.5019852891855e-51 | 2.89913200518586e-47 | Intermediate mesoderm | 48.08 |
| 5 | Mesp1 | 0.807139248253679 | 0.908 | 0.641 | 9.56870411189077e-64 | 1.84695126767716e-59 | Mixed mesoderm | 33.88 |
| 5 | Egln1 | 0.78711001224939 | 0.963 | 0.81 | 1.8849404791483e-104 | 3.63831211285205e-100 | Mixed mesoderm | 33.88 |
| 5 | Hist1h2ap | 0.681229175930095 | 0.98 | 0.872 | 2.20492978861805e-61 | 4.25595547799055e-57 | Mixed mesoderm | 33.88 |
| 5 | Higd1a | 0.661478526464121 | 0.998 | 0.988 | 1.17525090099923e-87 | 2.26846928910872e-83 | Mixed mesoderm | 33.88 |
| 5 | Fam162a | 0.64539339525363 | 1 | 0.998 | 5.46660819447377e-112 | 1.05516471369733e-107 | Mixed mesoderm | 33.88 |
| 5 | Hist1h2ae | 0.625329317737423 | 0.918 | 0.73 | 3.39886756964956e-54 | 6.56049418293757e-50 | Mixed mesoderm | 33.88 |
| 5 | 2410006H1 | 0.622293625026636 | 0.996 | 0.994 | 3.12908235949593e-67 | 6.03975477029905e-63 | Mixed mesoderm | 33.88 |
| 5 | Anxa2 | 0.603026124132713 | 0.814 | 0.512 | 8.01414327435397e-67 | 1.5468899348158e-62 | Mixed mesoderm | 33.88 |
| 5 | Pgk1 | 0.563250315400613 | 1 | 1 | 1.59381752782928e-101 | 3.07638659221607e-97 | Mixed mesoderm | 33.88 |
| 5 | Bnip3 | 0.551601520273647 | 0.976 | 0.854 | 6.00655248488571e-66 | 1.15938476063264e-61 | Mixed mesoderm | 33.88 |
| 5 | Ddit4 | 0.518867421304449 | 0.898 | 0.636 | 3.70754145154506e-70 | 7.15629650977227e-66 | Mixed mesoderm | 33.88 |
| 5 | Pfkp | 0.500922610572004 | 0.784 | 0.494 | 7.43609788327359e-67 | 1.43531561342947e-62 | Mixed mesoderm | 33.88 |
| 5 | Ero1l | 0.485161686890963 | 0.814 | 0.608 | 3.57570522859756e-54 | 6.90182623223901e-50 | Mixed mesoderm | 33.88 |
| 5 | Ptgds | 0.478836614280738 | 0.382 | 0.201 | 3.86616536758324e-26 | 7.46247239250918e-22 | Mixed mesoderm | 33.88 |
| 5 | Bsg | 0.474900411052726 | 0.998 | 0.998 | 8.19990946344742e-72 | 1.58274652463462e-67 | Mixed mesoderm | 33.88 |
| 5 | Cfc1 | 0.44873665906057 | 0.91 | 0.778 | 1.57038015064585e-36 | 3.03114776677662e-32 | Mixed mesoderm | 33.88 |
| 5 | Pgm1 | 0.446111399005466 | 0.843 | 0.617 | 1.41621486244112e-61 | 2.73357792748384e-57 | Mixed mesoderm | 33.88 |
| 5 | Upp1 | 0.432703280406721 | 0.68 | 0.506 | 1.52534210845509e-27 | 2.94421533774001e-23 | Mixed mesoderm | 33.88 |
| 5 | Gpc3 | 0.42048296105082 | 0.996 | 0.958 | 5.84692110585602e-51 | 1.12857271185233e-46 | Mixed mesoderm | 33.88 |
| 5 | Bnip3l | 0.409859679877601 | 0.996 | 0.993 | 1.95655727013426e-62 | 3.77654684281315e-58 | Mixed mesoderm | 33.88 |
| 6 | Krt18 | 1.26057276034133 | 0.928 | 0.439 | 2.52725305460398e-137 | 4.87810384599661e-133 | Mesenchyme | 64.77 |
| 6 | Krt8 | 1.02573614223622 | 0.923 | 0.521 | 1.22787310313303e-106 | 2.37004066366737e-102 | Mesenchyme | 64.77 |
| 6 | Hand1 | 0.992619030795856 | 0.998 | 0.765 | 4.58267802151489e-132 | 8.84548511712805e-128 | Mesenchyme | 64.77 |

**<u>Supplementary Table S1:</u> Differential gene expression analysis at day 4**

| Cluster | Gene | avg_logFC | pct.1 | pct.2 | p_val | p_val_adj | most expressed celltype | pct.celltype |
|---|---|---|---|---|---|---|---|---|
| 6 | Hist1h1b | 0.897683135631013 | 0.989 | 0.719 | 1.53022399093882e-95 | 2.95363834731011e-91 | Mesenchyme | 64.77 |
| 6 | Phlda2 | 0.855420176827334 | 0.987 | 0.844 | 1.59369087401074e-62 | 3.07614212501552e-58 | Mesenchyme | 64.77 |
| 6 | Msx2 | 0.828055754412265 | 0.974 | 0.606 | 5.5029609906802e-134 | 1.06218153042109e-129 | Mesenchyme | 64.77 |
| 6 | Asb4 | 0.808526627406991 | 0.91 | 0.466 | 1.89246189726724e-97 | 3.65282995410523e-93 | Mesenchyme | 64.77 |
| 6 | Hist1h2ap | 0.760553667527556 | 1 | 0.872 | 9.34829087307681e-83 | 1.80440710432129e-78 | Mesenchyme | 64.77 |
| 6 | Stard8 | 0.730975955130054 | 0.895 | 0.399 | 1.42783568971658e-144 | 2.75600844829094e-140 | Mesenchyme | 64.77 |
| 6 | Pmp22 | 0.687247391413798 | 0.976 | 0.61 | 2.57283066866528e-90 | 4.96607775665773e-86 | Mesenchyme | 64.77 |
| 6 | Ppic | 0.676666514775501 | 0.996 | 0.83 | 1.11019228947048e-106 | 2.14289315713592e-102 | Mesenchyme | 64.77 |
| 6 | Peg10 | 0.643460422012505 | 0.934 | 0.642 | 1.8967119740718e-86 | 3.6610334523534e-82 | Mesenchyme | 64.77 |
| 6 | Tbx3 | 0.632348984330499 | 0.982 | 0.677 | 5.8550521804923e-93 | 1.13014217187862e-88 | Mesenchyme | 64.77 |
| 6 | Ahnak | 0.628790448428847 | 0.713 | 0.236 | 7.02174752927155e-128 | 1.35533770809999e-123 | Mesenchyme | 64.77 |
| 6 | Spin2c | 0.627358864090043 | 0.65 | 0.242 | 3.79046385263403e-98 | 7.3163533283542e-94 | Mesenchyme | 64.77 |
| 6 | Fth1 | 0.615413786079325 | 1 | 1 | 2.90251800021429e-105 | 5.60244024401362e-101 | Mesenchyme | 64.77 |
| 6 | Tuba1a | 0.609514093938616 | 0.993 | 0.95 | 1.01411344790805e-63 | 1.95744177715212e-59 | Mesenchyme | 64.77 |
| 6 | Csrp2 | 0.605961794525144 | 0.993 | 0.946 | 5.35343148418475e-54 | 1.03331934507734e-49 | Mesenchyme | 64.77 |
| 6 | Hist1h2ae | 0.579200267347545 | 0.982 | 0.727 | 9.83526753915084e-67 | 1.89840334040689e-62 | Mesenchyme | 64.77 |
| 6 | Sparc | 0.562514002836878 | 0.982 | 0.709 | 9.27735613726503e-87 | 1.7907152816149e-82 | Mesenchyme | 64.77 |
| 7 | Rspo3 | 0.973367810939389 | 0.789 | 0.526 | 7.56987151680118e-54 | 1.46113660017296e-49 | Intermediate mesoderm | 67.57 |
| 7 | Ube2c | 0.844211601863901 | 0.995 | 0.847 | 3.23725202040993e-87 | 6.24854384979524e-83 | Intermediate mesoderm | 67.57 |
| 7 | Cited1 | 0.748425222276093 | 0.741 | 0.371 | 1.20191639077329e-79 | 2.31993901747061e-75 | Intermediate mesoderm | 67.57 |
| 7 | Cenpf | 0.748361652507439 | 1 | 0.839 | 6.16447897767245e-83 | 1.18986773227034e-78 | Intermediate mesoderm | 67.57 |
| 7 | Hoxaas3 | 0.682054617692 | 0.553 | 0.166 | 2.40876317751983e-88 | 4.64939468524878e-84 | Intermediate mesoderm | 67.57 |
| 7 | Crabp1 | 0.680717994979656 | 0.921 | 0.737 | 7.59268973280892e-51 | 1.46554097222678e-46 | Intermediate mesoderm | 67.57 |
| 7 | Lef1 | 0.651845392844019 | 0.975 | 0.752 | 3.27213679124057e-95 | 6.31587843445256e-91 | Intermediate mesoderm | 67.57 |
| 7 | Top2a | 0.642535951077244 | 0.995 | 0.87 | 2.14088427916408e-71 | 4.13233483564251e-67 | Intermediate mesoderm | 67.57 |
| 7 | Osr1 | 0.632657935908967 | 0.556 | 0.171 | 1.45647753216219e-97 | 2.81129293257946e-93 | Intermediate mesoderm | 67.57 |
| 7 | Hoxd9 | 0.620088414725646 | 0.596 | 0.149 | 2.80085824641605e-135 | 5.40621658723225e-131 | Intermediate mesoderm | 67.57 |
| 7 | Prc1 | 0.617730909441037 | 0.932 | 0.564 | 7.572706555771e-81 | 1.46168381939492e-76 | Intermediate mesoderm | 67.57 |
| 7 | Meis2 | 0.617359521338226 | 0.975 | 0.745 | 1.12051664019109e-77 | 2.16282121889684e-73 | Intermediate mesoderm | 67.57 |
| 7 | Mest | 0.611504189465002 | 0.537 | 0.544 | 0.00467917065169102 | 1 | Intermediate mesoderm | 67.57 |
| 7 | Cks2 | 0.606264829890544 | 1 | 0.958 | 1.8969042732377e-85 | 3.66140462820341e-81 | Intermediate mesoderm | 67.57 |
| 7 | H2afx | 0.557519759644519 | 1 | 0.962 | 4.77096048103879e-58 | 9.20890792050107e-54 | Intermediate mesoderm | 67.57 |
| 7 | Rbp1 | 0.550860685177741 | 0.95 | 0.897 | 9.23852614550349e-41 | 1.78322031660508e-36 | Intermediate mesoderm | 67.57 |
| 7 | Cdk1 | 0.532285889311958 | 0.991 | 0.83 | 6.20441507635732e-67 | 1.19757619803849e-62 | Intermediate mesoderm | 67.57 |
| 7 | Cenpe | 0.519889111700905 | 0.959 | 0.736 | 8.35380618704643e-59 | 1.6124516702237e-54 | Intermediate mesoderm | 67.57 |
| 7 | Grb10 | 0.503196683704003 | 0.907 | 0.758 | 3.15400065902424e-39 | 6.08785207204859e-35 | Intermediate mesoderm | 67.57 |
| 7 | Arl6ip1 | 0.499659799487845 | 0.993 | 0.899 | 5.80192436353775e-51 | 1.11988744065006e-46 | Intermediate mesoderm | 67.57 |
| 8 | Fgf8 | 1.85308547032312 | 0.897 | 0.395 | 1.79636640425431e-189 | 3.46734643349166e-185 | PGC | 37.61 |
| 8 | Hoxaas3 | 1.78724376901377 | 0.608 | 0.163 | 3.58406270714972e-147 | 6.9179578373404e-143 | PGC | 37.61 |
| 8 | T | 1.69526335699707 | 0.846 | 0.189 | 3.96651337214248e-291 | 7.65616411090941e-287 | PGC | 37.61 |
| 8 | Rspo3 | 1.47028370564661 | 0.89 | 0.519 | 9.04573657289067e-129 | 1.74600807329936e-124 | PGC | 37.61 |
| 8 | Fst | 1.31322760342332 | 0.628 | 0.288 | 5.22270527658079e-86 | 1.00808657248562e-81 | PGC | 37.61 |
| 8 | S100a11 | 1.21273574460934 | 0.803 | 0.68 | 3.06977934345813e-57 | 5.92528808874288e-53 | PGC | 37.61 |
| 8 | Cdx2 | 1.19604552126688 | 0.872 | 0.417 | 1.86909430837099e-157 | 3.60772583401768e-153 | PGC | 37.61 |
| 8 | Grsf1 | 1.10695660375504 | 0.839 | 0.756 | 1.66045088129038e-56 | 3.20500229106669e-52 | PGC | 37.61 |
| 8 | Defa30 | 1.09277807245861 | 0.537 | 0.154 | 3.96372073437655e-108 | 7.65077376149362e-104 | PGC | 37.61 |
| 8 | Ccnd2 | 1.06764644667588 | 0.947 | 0.818 | 4.17893493429108e-88 | 8.06618021016863e-84 | PGC | 37.61 |
| 8 | Psme2 | 1.04986781432232 | 0.922 | 0.754 | 3.8172929126381e-108 | 7.36813877997406e-104 | PGC | 37.61 |
| 8 | Bex1 | 1.04938706650185 | 0.977 | 0.841 | 3.54125512446367e-124 | 6.83533064123977e-120 | PGC | 37.61 |
| 8 | Crabp2 | 1.01989403783045 | 0.711 | 0.283 | 3.03464198622432e-126 | 5.85746596181019e-122 | PGC | 37.61 |
| 8 | Pou5f1 | 0.930741472720602 | 0.713 | 0.33 | 3.85965172120109e-92 | 7.44989975226234e-88 | PGC | 37.61 |
| 8 | Bex4 | 0.914996153344115 | 0.952 | 0.747 | 5.12545460709885e-115 | 9.89315248262221e-111 | PGC | 37.61 |
| 8 | Id1 | 0.911474126075047 | 0.961 | 0.913 | 5.31678245942412e-42 | 1.02624535031804e-37 | PGC | 37.61 |
| 8 | Lefty1 | 0.906604816865165 | 0.261 | 0.035 | 1.82891365669333e-100 | 3.53016914014947e-96 | PGC | 37.61 |
| 8 | Rbp1 | 0.810293683000592 | 0.888 | 0.901 | 3.82522719446556e-36 | 7.38345353075742e-32 | PGC | 37.61 |
| 8 | Cdx1 | 0.767670410266845 | 0.534 | 0.148 | 5.93811977185655e-115 | 1.14617587836375e-110 | PGC | 37.61 |
| 8 | Cdkn1a | 0.749382073225113 | 0.817 | 0.501 | 2.72183648435142e-71 | 5.25368878209511e-67 | PGC | 37.61 |
| 9 | Dppa5a | 4.53063240825147 | 0.954 | 0.415 | 1.32420750779028e-253 | 2.55598533153681e-249 | PGC | 54.68 |
| 9 | Mt1 | 2.76669474524997 | 0.914 | 0.318 | 3.00580568550979e-232 | 5.801806134171e-228 | PGC | 54.68 |
| 9 | Chchd10 | 2.22451223044246 | 0.964 | 0.251 | 0 | 0 | PGC | 54.68 |
| 9 | Mt2 | 2.22347019479736 | 0.827 | 0.129 | 0 | 0 | PGC | 54.68 |
| 9 | Mkrn1 | 2.17836344433081 | 0.969 | 0.693 | 1.35947195968487e-200 | 2.62405277658373e-196 | PGC | 54.68 |
| 9 | Pou5f1 | 2.02946651678121 | 0.993 | 0.312 | 0 | 0 | PGC | 54.68 |
| 9 | Tdh | 1.7351977616664 | 0.609 | 0.015 | 0 | 0 | PGC | 54.68 |
| 9 | Klf2 | 1.71863934905581 | 0.705 | 0.038 | 0 | 0 | PGC | 54.68 |
| 9 | Utf1 | 1.70942197069585 | 0.815 | 0.021 | 0 | 0 | PGC | 54.68 |
| 9 | Zfp42 | 1.58274393797051 | 0.734 | 0.067 | 0 | 0 | PGC | 54.68 |
| 9 | L1td1 | 1.55075231801741 | 0.916 | 0.121 | 0 | 0 | PGC | 54.68 |
| 9 | Rhox5 | 1.52102069538338 | 0.614 | 0.286 | 1.60948153410291e-71 | 3.10662125712544e-67 | PGC | 54.68 |
| 9 | Mylpf | 1.51046185777508 | 0.525 | 0.083 | 1.88377609625937e-199 | 3.63606462099984e-195 | PGC | 54.68 |
| 9 | Sox2 | 1.50832944139149 | 0.823 | 0.104 | 0 | 0 | PGC | 54.68 |
| 9 | Ckb | 1.45152849817888 | 0.847 | 0.262 | 1.54345779963528e-208 | 2.97918224485601e-204 | PGC | 54.68 |
| 9 | Hspb1 | 1.43406528460168 | 0.76 | 0.127 | 3.62664272132256e-295 | 7.0001457806968e-291 | PGC | 54.68 |
| 9 | Epcam | 1.2696468193446 | 0.88 | 0.25 | 1.28862843771373e-217 | 2.48731061047503e-213 | PGC | 54.68 |
| 9 | Dppa3 | 1.22376973781235 | 0.47 | 0.009 | 0 | 0 | PGC | 54.68 |
| 9 | Gpx4 | 1.15389610682374 | 0.995 | 0.997 | 3.58992390552037e-71 | 6.92927112243543e-67 | PGC | 54.68 |
| 9 | Bst2 | 1.14128637795101 | 0.875 | 0.241 | 3.81683977617372e-234 | 7.36726413597052e-230 | PGC | 54.68 |
| 10 | Emb | 2.88634931374905 | 0.993 | 0.687 | 3.84241841811576e-236 | 7.41663603064703e-232 | Gut | 77.31 |
| 10 | Spink1 | 2.62316101187881 | 0.591 | 0.042 | 0 | 0 | Gut | 77.31 |
| 10 | Slc2a3 | 2.393245328508 | 0.993 | 0.614 | 3.16056449055667e-231 | 6.10052157967249e-227 | Gut | 77.31 |
| 10 | Trh | 2.38499232384539 | 0.843 | 0.13 | 0 | 0 | Gut | 77.31 |
| 10 | Krt18 | 2.06179346857294 | 0.985 | 0.439 | 1.2578288998245e-218 | 2.42786134244125e-214 | Gut | 77.31 |
| 10 | Apoe | 1.87887934273153 | 0.998 | 0.827 | 2.28225342933149e-187 | 4.40520556929563e-183 | Gut | 77.31 |
| 10 | Epcam | 1.81278719142737 | 0.948 | 0.247 | 3.50570785364057e-294 | 6.76671729909702e-290 | Gut | 77.31 |
| 10 | Krt8 | 1.76033221864969 | 0.985 | 0.52 | 8.35911366480063e-196 | 1.61347611957982e-191 | Gut | 77.31 |
| 10 | Slc16a1 | 1.666930569827 | 0.95 | 0.629 | 2.18399949753854e-197 | 4.21555583014889e-193 | Gut | 77.31 |
| 10 | Sox17 | 1.58043648117905 | 0.873 | 0.025 | 0 | 0 | Gut | 77.31 |
| 10 | Cldn7 | 1.3714125031865 | 0.86 | 0.057 | 0 | 0 | Gut | 77.31 |
| 10 | Flt1 | 1.29345734179863 | 0.83 | 0.114 | 0 | 0 | Gut | 77.31 |
| 10 | Cldn6 | 1.28336055039319 | 0.87 | 0.054 | 0 | 0 | Gut | 77.31 |
| 10 | Bex4 | 1.2783588749459 | 0.983 | 0.746 | 5.40539817911971e-160 | 1.04334995653369e-155 | Gut | 77.31 |
| 10 | Car2 | 1.23831658362356 | 0.88 | 0.39 | 7.89669996209367e-174 | 1.52422102668332e-169 | Gut | 77.31 |
| 10 | Amot | 1.2209392467811 | 0.88 | 0.621 | 1.02021692802217e-108 | 1.96922271446839e-104 | Gut | 77.31 |
| 10 | Dnmt3b | 1.21672737328651 | 0.915 | 0.762 | 6.79735575188264e-111 | 1.31202560722839e-106 | Gut | 77.31 |
| 10 | Espn | 1.20471084896807 | 0.833 | 0.216 | 2.9879903142015e-216 | 5.76741890447173e-212 | Gut | 77.31 |
| 10 | Car4 | 1.19245484978661 | 0.599 | 0.149 | 8.27850782520138e-148 | 1.59791758042037e-143 | Gut | 77.31 |
| 10 | Pcbd1 | 1.17567116252147 | 0.81 | 0.153 | 3.18261992034627e-305 | 6.14309297025238e-301 | Gut | |
| 11 | Krt8 | 1.21929618019482 | 0.967 | 0.524 | 9.07905944263893e-122 | 1.75244005361817e-117 | Mesenchyme | 92.24 |
| 11 | Hand1 | 1.07638906074976 | 1 | 0.768 | 1.30404742886158e-129 | 2.51707234718863e-125 | Mesenchyme | 92.24 |
| 11 | Ube2c | 1.02770897855968 | 1 | 0.849 | 1.88725144978115e-98 | 3.64277274836757e-94 | Mesenchyme | 92.24 |
| 11 | Ahnak | 1.00433399516518 | 0.898 | 0.232 | 4.71761455240471e-211 | 9.10593960905158e-207 | Mesenchyme | 92.24 |
| 11 | Arl6ip1 | 0.901296283235516 | 1 | 0.9 | 7.98727140293537e-106 | 1.54170312619459e-101 | Mesenchyme | 92.24 |
| 11 | Asb4 | 0.901012756757259 | 0.975 | 0.469 | 3.08393224841825e-105 | 5.95260602589691e-101 | Mesenchyme | 92.24 |

**<u>Supplementary Table S1:</u> Differential gene expression analysis at day 4**

| Cluster | Gene | avg_logFC | pct.1 | pct.2 | p_val | p_val_adj | most expressed celltype | pct.celltype |
|---|---|---|---|---|---|---|---|---|
| 11 | Stard8 | 0.852252524279046 | 0.95 | 0.403 | 2.01746782320581e-149 | 3.89411639235186e-145 | Mesenchyme | 92.24 |
| 11 | Tuba1a | 0.824899151395243 | 1 | 0.95 | 9.67463534425162e-97 | 1.86739811414745e-92 | Mesenchyme | 92.24 |
| 11 | Pmp22 | 0.819371782334138 | 0.992 | 0.614 | 1.03796680141214e-100 | 2.00348352008572e-96 | Mesenchyme | 92.24 |
| 11 | Peg10 | 0.804577248876936 | 0.978 | 0.644 | 7.95174331185541e-92 | 1.53484549405433e-87 | Mesenchyme | 92.24 |
| 11 | Spin2c | 0.794801438695191 | 0.801 | 0.24 | 2.46761352852062e-151 | 4.7629876327505e-147 | Mesenchyme | 92.24 |
| 11 | Csrp2 | 0.791155949414126 | 1 | 0.946 | 2.36962730626749e-76 | 4.57385462655752e-72 | Mesenchyme | 92.24 |
| 11 | Nusap1 | 0.783757011435185 | 0.978 | 0.55 | 5.5669313529859e-106 | 1.07452908975334e-101 | Mesenchyme | 92.24 |
| 11 | Lgals1 | 0.770740994881067 | 0.961 | 0.681 | 1.43127818182105e-70 | 2.76265314655098e-66 | Mesenchyme | 92.24 |
| 11 | Cenpf | 0.761697514081321 | 1 | 0.842 | 1.22687446866211e-79 | 2.3681130994116e-75 | Mesenchyme | 92.24 |
| 11 | Msx2 | 0.728859757416437 | 0.986 | 0.611 | 2.5324324414299e-102 | 4.88810109844799e-98 | Mesenchyme | 92.24 |
| 11 | Map1b | 0.708151893283211 | 0.898 | 0.605 | 1.21120546191124e-65 | 2.33786878258108e-61 | Mesenchyme | 92.24 |
| 11 | Prc1 | 0.706668368370212 | 0.97 | 0.566 | 3.678568491133e-88 | 7.10037290158491e-84 | Mesenchyme | 92.24 |
| 11 | Fth1 | 0.687771395204057 | 1 | 1 | 8.60973924700062e-100 | 1.66185186945606e-95 | Mesenchyme | 92.24 |
| 12 | Csrp2 | 0.771136672120798 | 0.991 | 0.947 | 7.21340688481035e-60 | 1.39233179690609e-55 | Pharyngeal mesoderm | 49.55 |
| 12 | Krt18 | 0.676016958833061 | 0.767 | 0.457 | 6.65053907751162e-36 | 1.28368705274129e-31 | Pharyngeal mesoderm | 49.55 |
| 12 | Tuba1a | 0.660815734337742 | 0.985 | 0.951 | 1.67279295161857e-47 | 3.22882495521417e-43 | Pharyngeal mesoderm | 49.55 |
| 12 | Krt8 | 0.622881324001965 | 0.749 | 0.538 | 1.16180806303084e-26 | 2.24252192326212e-22 | Pharyngeal mesoderm | 49.55 |
| 12 | Lgals1 | 0.54175074197207 | 0.798 | 0.69 | 2.45588773368872e-15 | 4.74035450356596e-11 | Pharyngeal mesoderm | 49.55 |
| 12 | Phlda2 | 0.533730148434748 | 0.958 | 0.848 | 1.47728309752868e-20 | 2.85145183484986e-16 | Pharyngeal mesoderm | 49.55 |
| 12 | Fth1 | 0.527206973230507 | 1 | 1 | 7.83000787156876e-57 | 1.5113481193702e-52 | Pharyngeal mesoderm | 49.55 |
| 12 | Peg10 | 0.526041991144991 | 0.834 | 0.653 | 2.21217690547189e-32 | 4.26994386294184e-28 | Pharyngeal mesoderm | 49.55 |
| 12 | Vim | 0.519572186169791 | 0.994 | 0.959 | 1.01594716391051e-36 | 1.96098121578007e-32 | Pharyngeal mesoderm | 49.55 |
| 12 | Hand1 | 0.500811701652871 | 0.964 | 0.771 | 3.29457020495888e-31 | 6.35917940961162e-27 | Pharyngeal mesoderm | 49.55 |
| 12 | Tmsb4x | 0.492913763655255 | 0.915 | 0.817 | 1.15874483739377e-20 | 2.23660928513745e-16 | Pharyngeal mesoderm | 49.55 |
| 12 | 1810058I24 | 0.490269577899521 | 0.997 | 0.971 | 3.63921030287789e-44 | 7.02440372661491e-40 | Pharyngeal mesoderm | 49.55 |
| 12 | Hspa8 | 0.48100161625235 | 1 | 1 | 2.20294001617839e-66 | 4.25211481922754e-62 | Pharyngeal mesoderm | 49.55 |
| 12 | Ahnak | 0.476122396016489 | 0.474 | 0.258 | 2.40512832765469e-25 | 4.64237869803908e-21 | Pharyngeal mesoderm | 49.55 |
| 12 | Nrp1 | 0.475247399237542 | 0.855 | 0.629 | 1.21963811818967e-30 | 2.3541454957297e-26 | Pharyngeal mesoderm | 49.55 |
| 12 | Hspa5 | 0.469661958803515 | 0.976 | 0.952 | 2.92293368748133e-30 | 5.64184660357646e-26 | Pharyngeal mesoderm | 49.55 |
| 12 | Cd24a | 0.436659533476986 | 0.982 | 0.907 | 1.96990158796171e-34 | 3.80230404508369e-30 | Pharyngeal mesoderm | 49.55 |
| 12 | Msx2 | 0.43475157238609 | 0.861 | 0.619 | 1.42553723010956e-31 | 2.75157196155748e-27 | Pharyngeal mesoderm | 49.55 |
| 12 | Igf2 | 0.433145814514003 | 0.9 | 0.687 | 7.80353548088642e-26 | 1.5062384185207e-21 | Pharyngeal mesoderm | 49.55 |
| 12 | Pmp22 | 0.430658124387288 | 0.867 | 0.623 | 5.54795669137748e-28 | 1.07086660056968e-23 | Pharyngeal mesoderm | 49.55 |
| 13 | Rspo3 | 1.46920076555662 | 0.943 | 0.524 | 2.31628383089195e-112 | 4.47089105038764e-108 | Intermediate mesoderm | 73.33 |
| 13 | Cited1 | 1.4583616490247 | 0.887 | 0.372 | 3.68439772837351e-135 | 7.11162449530655e-131 | Intermediate mesoderm | 73.33 |
| 13 | Hoxaas3 | 1.35135058733163 | 0.663 | 0.17 | 6.93528080735546e-114 | 1.33864790143575e-109 | Intermediate mesoderm | 73.33 |
| 13 | Rbp1 | 1.16697794235881 | 0.993 | 0.896 | 3.23676287823053e-106 | 6.24759970756057e-102 | Intermediate mesoderm | 73.33 |
| 13 | Smc6 | 0.944709686709157 | 0.987 | 0.954 | 8.70224081483587e-79 | 1.67970652207962e-74 | Intermediate mesoderm | 73.33 |
| 13 | Ifitm1 | 0.886673773864465 | 1 | 0.98 | 2.08775288793303e-77 | 4.02978062428833e-73 | Intermediate mesoderm | 73.33 |
| 13 | Cdx2 | 0.700357370234703 | 0.83 | 0.428 | 2.51961891563361e-67 | 4.86336843095599e-63 | Intermediate mesoderm | 73.33 |
| 13 | Fgf8 | 0.664457960301175 | 0.76 | 0.412 | 8.64779104468352e-45 | 1.66919662744481e-40 | Intermediate mesoderm | 73.33 |
| 13 | Hoxb1 | 0.6570264400626 | 0.913 | 0.575 | 1.93088039773922e-54 | 3.72698534371624e-50 | Intermediate mesoderm | 73.33 |
| 13 | Lef1 | 0.649845896848203 | 0.94 | 0.759 | 1.54372928567113e-56 | 2.97970626720242e-52 | Intermediate mesoderm | 73.33 |
| 13 | S100a11 | 0.648743899614164 | 0.797 | 0.682 | 4.07268459876386e-29 | 7.86109581253399e-25 | Intermediate mesoderm | 73.33 |
| 13 | Hoxb5os | 0.646536517129774 | 0.447 | 0.1 | 7.12689446143773e-80 | 1.37563316894671e-75 | Intermediate mesoderm | 73.33 |
| 13 | Aldh1a2 | 0.623755412150774 | 0.637 | 0.406 | 1.30564212017432e-24 | 2.52015042036046e-20 | Intermediate mesoderm | 73.33 |
| 13 | Sms | 0.604270884310501 | 0.99 | 0.928 | 4.64013075173222e-54 | 8.95638037699352e-50 | Intermediate mesoderm | 73.33 |
| 13 | Dll3 | 0.598967863925068 | 0.737 | 0.321 | 2.26745878410271e-71 | 4.37664894507504e-67 | Intermediate mesoderm | 73.33 |
| 13 | mt-Cytb | 0.571840853631568 | 1 | 0.997 | 4.95921063022305e-44 | 9.57226835845653e-40 | Intermediate mesoderm | 73.33 |
| 13 | Cdx1 | 0.553207523613281 | 0.537 | 0.156 | 4.69166647842978e-75 | 9.05585463666516e-71 | Intermediate mesoderm | 73.33 |
| 13 | Rnf213 | 0.541715718549728 | 0.633 | 0.207 | 6.06250305910095e-84 | 1.17018434046767e-79 | Intermediate mesoderm | 73.33 |
| 13 | mt-Nd4 | 0.534801366685639 | 1 | 0.997 | 5.61684801555477e-40 | 1.08416400396238e-35 | Intermediate mesoderm | 73.33 |
| 13 | Crabp1 | 0.534532320411808 | 0.88 | 0.743 | 5.13074275450867e-22 | 9.90335966475263e-18 | Intermediate mesoderm | 73.33 |
| 14 | Ptgds | 1.8607946824372 | 0.707 | 0.201 | 3.65036797306118e-80 | 7.04594026160269e-76 | Nascent mesoderm | 36.78 |
| 14 | Anxa2 | 1.18344603421434 | 0.868 | 0.525 | 2.51320246750512e-52 | 4.85098340277838e-48 | Nascent mesoderm | 36.78 |
| 14 | Mesp1 | 1.14098851855936 | 0.868 | 0.655 | 3.64332414121993e-30 | 7.03234425738271e-26 | Nascent mesoderm | 36.78 |
| 14 | 2410006H1 | 0.939084058197934 | 1 | 0.994 | 7.1155674477719e-52 | 1.37344682876893e-47 | Nascent mesoderm | 36.78 |
| 14 | Egln1 | 0.891530891308081 | 0.925 | 0.819 | 9.08760451358558e-38 | 1.75408942321229e-33 | Nascent mesoderm | 36.78 |
| 14 | Fam162a | 0.854236412592001 | 1 | 0.998 | 2.31658727124856e-57 | 4.47147675096397e-53 | Nascent mesoderm | 36.78 |
| 14 | Bsg | 0.820336968733183 | 1 | 0.998 | 4.50610670515911e-52 | 8.69768716229811e-48 | Nascent mesoderm | 36.78 |
| 14 | Ddit4 | 0.82016120347857 | 0.897 | 0.649 | 3.4908329051237e-42 | 6.73800567346977e-38 | Nascent mesoderm | 36.78 |
| 14 | Aldoa | 0.806136439573427 | 1 | 1 | 1.94188680470358e-69 | 3.74822991043885e-65 | Nascent mesoderm | 36.78 |
| 14 | Pgk1 | 0.804015246616566 | 1 | 1 | 9.97553452519965e-57 | 1.92547767405404e-52 | Nascent mesoderm | 36.78 |
| 14 | Ero1l | 0.759868162594702 | 0.753 | 0.62 | 2.62480388354299e-23 | 5.06639645601468e-19 | Nascent mesoderm | 36.78 |
| 14 | Aldh1a2 | 0.738628646876574 | 0.586 | 0.412 | 3.47768622584669e-12 | 6.71262995312928e-8 | Nascent mesoderm | 36.78 |
| 14 | Bnip3 | 0.710403711599033 | 0.954 | 0.861 | 2.53626088915184e-33 | 4.89549076824088e-29 | Nascent mesoderm | 36.78 |
| 14 | Cox6b2 | 0.693167369782268 | 0.81 | 0.602 | 1.22324089432125e-29 | 2.36109957421888e-25 | Nascent mesoderm | 36.78 |
| 14 | Gpi1 | 0.683095933716275 | 1 | 0.991 | 3.45738359004429e-44 | 6.6734418055035e-40 | Nascent mesoderm | 36.78 |
| 14 | Pfkp | 0.656964208351039 | 0.718 | 0.509 | 8.65034261876044e-23 | 1.66968913227314e-18 | Nascent mesoderm | 36.78 |
| 14 | P4ha2 | 0.65408213426921 | 0.718 | 0.424 | 5.02151468971071e-37 | 9.69252765407961e-33 | Nascent mesoderm | 36.78 |
| 14 | Ier3 | 0.645792403952964 | 0.42 | 0.148 | 6.2778129622556e-28 | 1.21174345797458e-23 | Nascent mesoderm | 36.78 |
| 14 | Mt1 | 0.639546881818428 | 0.58 | 0.349 | 7.28616604951923e-16 | 1.4063757708782e-11 | Nascent mesoderm | 36.78 |
| 14 | Adssl1 | 0.604792281917073 | 0.563 | 0.25 | 4.64442178111941e-34 | 8.96466292191669e-30 | Nascent mesoderm | 36.78 |
| 15 | Fth1 | 0.663300590716553 | 1 | 1 | 3.30998689162452e-24 | 6.38893669821366e-20 | Mixed mesoderm | 47.25 |
| 15 | Snrpg | 0.580558747387605 | 1 | 1 | 2.55356622417816e-40 | 4.92889352590868e-36 | Mixed mesoderm | 47.25 |
| 15 | Tmsb10 | 0.575451174663774 | 1 | 1 | 1.38887615076186e-25 | 2.68080874620054e-21 | Mixed mesoderm | 47.25 |
| 15 | Sec61b | 0.566689873775695 | 1 | 0.998 | 3.01427532141292e-32 | 5.81815422539122e-28 | Mixed mesoderm | 47.25 |
| 15 | Atp5md | 0.564022101031463 | 1 | 0.996 | 2.30641093937911e-30 | 4.45183439518956e-26 | Mixed mesoderm | 47.25 |
| 15 | Gng5 | 0.558799985568402 | 1 | 1 | 5.94888843536317e-32 | 1.1482544457938e-27 | Mixed mesoderm | 47.25 |
| 15 | Mrpl20 | 0.539096097038961 | 1 | 0.939 | 1.87321782488424e-25 | 3.61568504559156e-21 | Mixed mesoderm | 47.25 |
| 15 | Gm10076 | 0.529076695327217 | 1 | 1 | 1.898462776177e-29 | 3.66441285057685e-25 | Mixed mesoderm | 47.25 |
| 15 | Atp5g1 | 0.524542131849598 | 1 | 0.997 | 3.61554411621939e-31 | 6.97872325312667e-27 | Mixed mesoderm | 47.25 |
| 15 | Atp5l | 0.517988863503903 | 1 | 0.998 | 9.69591416892214e-29 | 1.87150535288535e-24 | Mixed mesoderm | 47.25 |
| 15 | Chchd2 | 0.517585952233046 | 1 | 1 | 9.9167505282354e-38 | 1.91413118696e-33 | Mixed mesoderm | 47.25 |
| 15 | Sec61g | 0.514840477745059 | 1 | 1 | 5.91920860338141e-31 | 1.14252564462468e-26 | Mixed mesoderm | 47.25 |
| 15 | Hspe1 | 0.509607653632378 | 1 | 1 | 6.48418903145161e-34 | 1.25157816685079e-29 | Mixed mesoderm | 47.25 |
| 15 | Tomm7 | 0.502868632046745 | 1 | 0.999 | 1.34930978644693e-28 | 2.60443774979987e-24 | Mixed mesoderm | 47.25 |
| 15 | Cox6b1 | 0.500786216208401 | 1 | 1 | 1.68789035235921e-33 | 3.25796595812375e-29 | Mixed mesoderm | 47.25 |
| 15 | Snrpe | 0.500306565155793 | 1 | 1 | 3.7011785460546e-38 | 7.14401482959458e-34 | Mixed mesoderm | 47.25 |
| 15 | H2afz | 0.489934002497108 | 1 | 1 | 2.5209849339515e-29 | 4.86600511951318e-25 | Mixed mesoderm | 47.25 |
| 15 | Tbca | 0.481927856938237 | 1 | 0.994 | 9.26980870040897e-23 | 1.78925847535294e-18 | Mixed mesoderm | 47.25 |
| 15 | Sem1 | 0.476069283559394 | 1 | 1 | 6.8001097155704e-31 | 1.3125571772994e-26 | Mixed mesoderm | 47.25 |
| 15 | Dbi | 0.467635071787252 | 1 | 0.994 | 1.89288804065138e-22 | 3.65365249606529e-18 | Mixed mesoderm | 47.25 |

**Supplementary Table S1:** Differential gene expression analysis at day 5

| Cluster | Gene | avg_logFC | pct.1 | pct.2 | p_val | p_val_adj | most expressed celltype | pct.celltype |
|---|---|---|---|---|---|---|---|---|
| 1 | Hand1 | 0.643146055280597 | 0.94 | 0.679 | 1.66194314351148e-104 | 3.40781441577029e-100 | Mesenchyme | 59.7 |
| 1 | Vim | 0.678787735110705 | 0.999 | 0.958 | 1.21473489501971e-102 | 2.49081390223792e-98 | Mesenchyme | 59.7 |
| 1 | Bambi | 0.510761184557094 | 0.974 | 0.844 | 4.0966344925943e-91 | 8.40014902706461e-87 | Mesenchyme | 59.7 |
| 1 | Peg3 | 0.573339574194454 | 0.996 | 0.928 | 2.24547815046212e-87 | 4.60435294752258e-83 | Mesenchyme | 59.7 |
| 1 | Pmp22 | 0.514114087171967 | 0.907 | 0.693 | 2.67622898752072e-76 | 5.48760753891123e-72 | Mesenchyme | 59.7 |
| 1 | Hand2 | 0.420920079674075 | 0.778 | 0.527 | 8.30052603883608e-73 | 1.70202286426334e-68 | Mesenchyme | 59.7 |
| 1 | Ddt | 0.379230297197591 | 0.971 | 0.933 | 4.11263149358204e-72 | 8.43295087758997e-68 | Mesenchyme | 59.7 |
| 1 | Csrp2 | 0.539876677645575 | 0.995 | 0.981 | 5.64263905535465e-72 | 1.15702313830047e-67 | Mesenchyme | 59.7 |
| 1 | Ptpn18 | 0.417332534476073 | 0.918 | 0.751 | 3.53296456068961e-69 | 7.24434383169404e-65 | Mesenchyme | 59.7 |
| 1 | Rnd2 | 0.359915462300879 | 0.801 | 0.612 | 1.44311308153297e-63 | 2.95910337368336e-59 | Mesenchyme | 59.7 |
| 1 | Lgals1 | 0.69541526483914 | 0.948 | 0.889 | 3.04851329584988e-63 | 6.25097651314017e-59 | Mesenchyme | 59.7 |
| 1 | Arhgap29 | 0.354970526238314 | 0.772 | 0.569 | 7.50456130124278e-63 | 1.53881029481983e-58 | Mesenchyme | 59.7 |
| 1 | Rnd3 | 0.348911000751058 | 0.726 | 0.525 | 9.58641115379151e-63 | 1.96569360708495e-58 | Mesenchyme | 59.7 |
| 1 | Prrx2 | 0.383098583687527 | 0.735 | 0.532 | 2.90856560018251e-57 | 5.96401376317423e-53 | Mesenchyme | 59.7 |
| 1 | Ppic | 0.378861629891647 | 0.972 | 0.897 | 7.9254461992281e-49 | 1.62511274315172e-44 | Mesenchyme | 59.7 |
| 1 | Rdx | 0.359298683405803 | 0.993 | 0.976 | 2.20895584931838e-38 | 4.52946396902734e-34 | Mesenchyme | 59.7 |
| 1 | Twist1 | 0.389962427234045 | 0.789 | 0.684 | 8.52135866241451e-32 | 1.74730459328809e-27 | Mesenchyme | 59.7 |
| 1 | Tmsb4x | 0.420732195097075 | 0.981 | 0.962 | 1.34724459860004e-30 | 2.76252504942939e-26 | Mesenchyme | 59.7 |
| 1 | Dlk1 | 0.370288917499112 | 0.956 | 0.88 | 1.14942962629453e-24 | 2.35690544871694e-20 | Mesenchyme | 59.7 |
| 1 | Hmgcs1 | 0.351211742796483 | 0.884 | 0.857 | 2.14221217699767e-23 | 4.39260606893373e-19 | Mesenchyme | 59.7 |
| 2 | Ube2c | 1.75120666217474 | 1 | 0.816 | 0 | 0 | Mesenchyme | 54.22 |
| 2 | Cenpf | 1.47498129266329 | 1 | 0.699 | 0 | 0 | Mesenchyme | 54.22 |
| 2 | Arl6ip1 | 1.37002680197825 | 1 | 0.964 | 0 | 0 | Mesenchyme | 54.22 |
| 2 | Nusap1 | 1.22189638101566 | 0.988 | 0.438 | 0 | 0 | Mesenchyme | 54.22 |
| 2 | Prc1 | 1.08734411662497 | 0.985 | 0.503 | 0 | 0 | Mesenchyme | 54.22 |
| 2 | Hmmr | 1.04907725200338 | 0.994 | 0.551 | 0 | 0 | Mesenchyme | 54.22 |
| 2 | Ccnb1 | 1.02523874548805 | 0.995 | 0.613 | 0 | 0 | Mesenchyme | 54.22 |
| 2 | Plk1 | 0.8028104120157 | 0.955 | 0.296 | 0 | 0 | Mesenchyme | 54.22 |
| 2 | Cks2 | 1.07954308094708 | 1 | 0.878 | 2.02446151964122e-304 | 4.15115834602432e-300 | Mesenchyme | 54.22 |
| 2 | Cenpa | 1.14085601769125 | 1 | 0.728 | 1.37434897296475e-291 | 2.81810256906422e-287 | Mesenchyme | 54.22 |
| 2 | Tpx2 | 0.927200384331888 | 0.993 | 0.636 | 1.90295230002644e-276 | 3.90200369120422e-272 | Mesenchyme | 54.22 |
| 2 | Cdc20 | 0.995134871667126 | 0.994 | 0.619 | 3.19599338014583e-265 | 6.55338442598902e-261 | Mesenchyme | 54.22 |
| 2 | Cdk1 | 0.99844489891483 | 0.996 | 0.785 | 7.12696026972529e-264 | 1.46138320330717e-259 | Mesenchyme | 54.22 |
| 2 | Cdca8 | 0.80330718603372 | 0.999 | 0.828 | 5.7063158589588e-249 | 1.1700800668795e-244 | Mesenchyme | 54.22 |
| 2 | Tubb4b | 0.956637320063706 | 1 | 0.946 | 6.49474977212627e-243 | 1.33174844077449e-238 | Mesenchyme | 54.22 |
| 2 | Jpt1 | 0.813237817708341 | 1 | 0.973 | 1.57375995276438e-241 | 3.22699478314336e-237 | Mesenchyme | 54.22 |
| 2 | Top2a | 1.04035873323279 | 0.995 | 0.802 | 7.58561974068428e-240 | 1.55543132782731e-235 | Mesenchyme | 54.22 |
| 2 | Cenpe | 0.756252176741111 | 0.983 | 0.631 | 4.35688043345318e-228 | 8.93378332879575e-224 | Mesenchyme | 54.22 |
| 2 | Mki67 | 0.786222440754101 | 0.977 | 0.574 | 6.19573310424481e-224 | 1.2704350730254e-219 | Mesenchyme | 54.22 |
| 2 | H2afx | 0.887610794713665 | 1 | 0.939 | 2.10954302531314e-220 | 4.3256179734046e-216 | Mesenchyme | 54.22 |
| 3 | Hist1h1b | 0.965171097402129 | 0.943 | 0.499 | 5.84835013276969e-212 | 1.19920419472442e-207 | Pharyngeal mesoderm | 48.2 |
| 3 | Hist1h2ap | 0.876208174842507 | 0.983 | 0.657 | 3.00831413111465e-195 | 6.16854812585058e-191 | Pharyngeal mesoderm | 48.2 |
| 3 | Pcna | 0.54824987258717 | 0.998 | 0.935 | 1.86598796224325e-158 | 3.82620831657978e-154 | Pharyngeal mesoderm | 48.2 |
| 3 | Tyms | 0.574990261601469 | 0.995 | 0.888 | 6.46639649302851e-147 | 1.3259346008955e-142 | Pharyngeal mesoderm | 48.2 |
| 3 | Hist1h2ae | 0.505302970182744 | 0.82 | 0.435 | 3.97461422613024e-137 | 8.14994647068006e-133 | Pharyngeal mesoderm | 48.2 |
| 3 | Fgf15 | 0.458105023716603 | 0.642 | 0.254 | 4.49937913737469e-132 | 9.22597692118679e-128 | Pharyngeal mesoderm | 48.2 |
| 3 | Hist1h1a | 0.433555834472409 | 0.822 | 0.443 | 3.22164699426163e-128 | 6.60598716173348e-124 | Pharyngeal mesoderm | 48.2 |
| 3 | Rrm2 | 0.590189132028055 | 0.994 | 0.815 | 9.59444286453863e-120 | 1.96734050937365e-115 | Pharyngeal mesoderm | 48.2 |
| 3 | Pclaf | 0.5922358209853243 | 1 | 0.884 | 5.24761556129223e-115 | 1.07602357084297e-110 | Pharyngeal mesoderm | 48.2 |
| 3 | Nasp | 0.488176621262125 | 1 | 0.975 | 2.73613134014249e-112 | 5.61043731296218e-108 | Pharyngeal mesoderm | 48.2 |
| 3 | Tubb5 | 0.523821703653539 | 1 | 0.987 | 2.78349394504539e-101 | 5.70755433431556e-97 | Pharyngeal mesoderm | 48.2 |
| 3 | Dhfr | 0.438993898259365 | 0.989 | 0.866 | 1.64856476323994e-100 | 3.3803820470235e-96 | Pharyngeal mesoderm | 48.2 |
| 3 | Dut | 0.40737307455459 | 1 | 0.973 | 1.42275628574363e-97 | 2.91736176391732e-93 | Pharyngeal mesoderm | 48.2 |
| 3 | Igfbp4 | 0.515746688231718 | 0.996 | 0.818 | 1.58958666064054e-97 | 3.25944744764342e-93 | Pharyngeal mesoderm | 48.2 |
| 3 | Fgf3 | 0.40996255844013 | 0.655 | 0.322 | 2.52030533658351e-86 | 5.16788609266448e-82 | Pharyngeal mesoderm | 48.2 |
| 3 | Hist1h1e | 0.413178139416011 | 0.897 | 0.621 | 1.9268116176777e-80 | 3.95092722204811e-76 | Pharyngeal mesoderm | 48.2 |
| 3 | Rdx | 0.460158058835659 | 0.999 | 0.976 | 9.06453627574669e-61 | 1.85868316334186e-56 | Pharyngeal mesoderm | 48.2 |
| 3 | Ccnd1 | 0.418837857340698 | 0.978 | 0.888 | 1.25581958358242e-56 | 2.57505805613575e-52 | Pharyngeal mesoderm | 48.2 |
| 3 | Vim | 0.480044739446126 | 1 | 0.959 | 1.1079395815301e-52 | 2.27183011192748e-48 | Pharyngeal mesoderm | 48.2 |
| 3 | Dlk1 | 0.5627991629693 | 0.973 | 0.88 | 4.22486031228666e-52 | 8.6630760703438e-48 | Pharyngeal mesoderm | 48.2 |
| 4 | Hist1h2ap | 0.779630005135194 | 0.97 | 0.661 | 4.64174500120357e-162 | 9.51789812496792e-158 | Mesenchyme | 84.26 |
| 4 | Hist1h1b | 0.741303589369115 | 0.903 | 0.507 | 2.68399406657945e-131 | 5.50352983352117e-127 | Mesenchyme | 84.26 |
| 4 | Csrp2 | 0.738215726867399 | 1 | 0.98 | 2.31694820487733e-122 | 4.75090229410096e-118 | Mesenchyme | 84.26 |
| 4 | Tmem88 | 0.503280925705453 | 0.921 | 0.632 | 2.68087613501901e-116 | 5.49713651485649e-112 | Mesenchyme | 84.26 |
| 4 | Bex3 | 0.50948205509033 | 1 | 0.996 | 2.10773185513675e-114 | 4.3219041689579e-110 | Mesenchyme | 84.26 |
| 4 | Hand1 | 0.754479276846555 | 0.966 | 0.686 | 8.17708683019855e-114 | 1.67671165453221e-109 | Mesenchyme | 84.26 |
| 4 | Tceal9 | 0.592835686975574 | 1 | 0.988 | 5.04931270040706e-112 | 1.03536156921847e-107 | Mesenchyme | 84.26 |
| 4 | Tmem108 | 0.459962792508285 | 0.792 | 0.414 | 3.83235828695939e-110 | 7.85825066741022e-106 | Mesenchyme | 84.26 |
| 4 | Pmp22 | 0.608412326887972 | 0.962 | 0.694 | 1.73160960169175e-100 | 3.55066548826894e-96 | Mesenchyme | 84.26 |
| 4 | Ppic | 0.545425820182308 | 0.995 | 0.897 | 2.95962597641027e-100 | 6.06871306462926e-96 | Mesenchyme | 84.26 |
| 4 | Krt8 | 0.677258068646933 | 0.987 | 0.871 | 1.44683233986154e-95 | 2.96672971288608e-91 | Mesenchyme | 84.26 |
| 4 | Krt18 | 0.542829601761896 | 0.971 | 0.749 | 2.82714383302666e-93 | 5.79705842962118e-89 | Mesenchyme | 84.26 |
| 4 | Pclaf | 0.516563608328427 | 0.997 | 0.885 | 1.78222530594219e-89 | 3.65445298983446e-85 | Mesenchyme | 84.26 |
| 4 | Msx2 | 0.450711898731102 | 0.858 | 0.532 | 1.78162945832608e-87 | 3.65323120429763e-83 | Mesenchyme | 84.26 |
| 4 | Atg3 | 0.485479941551821 | 0.946 | 0.818 | 5.51612448245833e-82 | 1.13108132512808e-77 | Mesenchyme | 84.26 |
| 4 | Csrp1 | 0.460470131764285 | 0.948 | 0.819 | 9.85508219957753e-80 | 2.02078460502337e-75 | Mesenchyme | 84.26 |
| 4 | Sparc | 0.425971353369461 | 0.991 | 0.895 | 6.27646690253253e-74 | 1.28698953836429e-69 | Mesenchyme | 84.26 |
| 4 | Tuba1a | 0.552832357718107 | 0.995 | 0.936 | 9.34027097309138e-71 | 1.91522256303239e-66 | Mesenchyme | 84.26 |
| 4 | Tpm1 | 0.486163120222137 | 0.997 | 0.98 | 5.44005667271186e-68 | 1.11548362073957e-63 | Mesenchyme | 84.26 |
| 4 | Mest | 0.638731242287334 | 0.837 | 0.706 | 1.27081229300644e-46 | 2.6058006068097e-42 | Mesenchyme | 84.26 |
| 5 | Spink1 | 3.33455701552471 | 0.785 | 0.053 | 0 | 0 | Gut | 57.65 |
| 5 | Emb | 3.07684765206537 | 0.993 | 0.74 | 0 | 0 | Gut | 57.65 |
| 5 | Cldn7 | 2.0544900196624 | 0.977 | 0.112 | 0 | 0 | Gut | 57.65 |
| 5 | Epcam | 1.83817615493938 | 0.982 | 0.388 | 0 | 0 | Gut | 57.65 |
| 5 | Pcbd1 | 1.54981164659401 | 0.97 | 0.509 | 0 | 0 | Gut | 57.65 |
| 5 | Car4 | 1.45164738434548 | 0.753 | 0.151 | 0 | 0 | Gut | 57.65 |
| 5 | Igfbp2 | 1.22129738621831 | 0.835 | 0.194 | 0 | 0 | Gut | 57.65 |
| 5 | Cldn6 | 1.21841571437115 | 0.897 | 0.05 | 0 | 0 | Gut | 57.65 |
| 5 | Pdpn | 1.11306207628771 | 0.811 | 0.106 | 0 | 0 | Gut | 57.65 |
| 5 | Ttr | 1.09964072547085 | 0.319 | 0.007 | 0 | 0 | Gut | 57.65 |
| 5 | Cystm1 | 1.21052928173868 | 0.972 | 0.433 | 5.79321000792037e-305 | 1.18789771212407e-300 | Gut | 57.65 |
| 5 | Slc2a3 | 2.18719317171388 | 0.995 | 0.59 | 6.43631130384685e-305 | 1.3197656328538e-300 | Gut | 57.65 |
| 5 | Spint2 | 1.47371842422008 | 0.975 | 0.535 | 4.00457777888026e-299 | 8.21138673559877e-295 | Gut | 57.65 |
| 5 | Tnfrsf12a | 1.19687153283284 | 0.742 | 0.142 | 1.20675000199707e-293 | 2.47444087909498e-289 | Gut | 57.65 |
| 5 | Slc16a1 | 1.30174327720441 | 0.95 | 0.491 | 5.929965272501e-262 | 1.21593937912633e-257 | Gut | 57.65 |
| 5 | Krt18 | 1.99633450986091 | 1 | 0.754 | 2.48854557291733e-256 | 5.10276269726699e-252 | Gut | 57.65 |

**Supplementary Table S1:** **Differential gene expression analysis at day 5**

| Cluster | Gene | avg_logFC | pct.1 | pct.2 | p_val | p_val_adj | most expressed celltype | pct.celltype |
|---|---|---|---|---|---|---|---|---|
| 5 | Cyba | 1.45725021968267 | 0.993 | 0.846 | 4.0968743389113e-232 | 8.40064083193761e-228 | Gut | 57.65 |
| 5 | Fth1 | 1.16077216089278 | 1 | 0.999 | 9.83958113614163e-159 | 2.01760611196584e-154 | Gut | 57.65 |
| 5 | Krt8 | 1.24458447898734 | 0.998 | 0.874 | 1.40604872721916e-156 | 2.88310291516288e-152 | Gut | 57.65 |
| 5 | Apoe | 1.49359019584649 | 0.993 | 0.907 | 5.45905566709243e-119 | 1.1193793645373e-114 | Gut | 57.65 |
| 6 | Krt15 | 1.1763671502117 | 0.495 | 0.052 | 7.10743002875186e-275 | 1.45737852739557e-270 | PGC | 34.61 |
| 6 | Perp | 1.25817623952587 | 0.909 | 0.357 | 3.80116833629804e-235 | 7.79429567357914e-231 | PGC | 34.61 |
| 6 | Actc1 | 1.2840461278112 | 0.425 | 0.043 | 1.40992397046879e-234 | 2.89104910144625e-230 | PGC | 34.61 |
| 6 | Rgcc | 1.79264950535741 | 0.519 | 0.095 | 2.19185894855991e-185 | 4.49440677402208e-181 | PGC | 34.61 |
| 6 | Fabp3 | 1.15298214921054 | 0.855 | 0.361 | 3.45916641536632e-180 | 7.09302073470863e-176 | PGC | 34.61 |
| 6 | Krt19 | 1.05066860129419 | 0.722 | 0.227 | 4.60786361748964e-174 | 9.4484243476625e-170 | PGC | 34.61 |
| 6 | Pgk1 | 1.03977850820481 | 1 | 0.997 | 6.78698049501135e-173 | 1.39167035050208e-168 | PGC | 34.61 |
| 6 | Crip2 | 1.2959871502852 | 0.936 | 0.59 | 1.1578835021751e-164 | 2.37424012121003e-160 | PGC | 34.61 |
| 6 | Txnip | 1.32253940803854 | 0.996 | 0.924 | 2.31596391285358e-140 | 4.74888400330626e-136 | PGC | 34.61 |
| 6 | Cdkn1a | 1.19073323972597 | 0.992 | 0.926 | 4.77927565648805e-140 | 9.79990473362874e-136 | PGC | 34.61 |
| 6 | Ddit4 | 0.921758640655392 | 0.944 | 0.711 | 9.81541950499774e-140 | 2.01265176949979e-135 | PGC | 34.61 |
| 6 | H1f0 | 1.58921906836839 | 0.885 | 0.594 | 3.63502829820821e-130 | 7.45362552547594e-126 | PGC | 34.61 |
| 6 | Ifi30 | 0.995160321065834 | 0.958 | 0.767 | 7.17049427425053e-130 | 1.47030985093507e-125 | PGC | 34.61 |
| 6 | Btg2 | 1.01879820517099 | 0.903 | 0.639 | 1.7516282450397e-124 | 3.59171371645391e-120 | PGC | 34.61 |
| 6 | Arrdc4 | 0.897294979056655 | 0.976 | 0.879 | 3.50224508033995e-120 | 7.18135353723706e-116 | PGC | 34.61 |
| 6 | 1500009L1 | 0.86823732637216 | 0.825 | 0.5 | 4.37242557807867e-114 | 8.96565864785032e-110 | PGC | 34.61 |
| 6 | Mdm2 | 1.058103482475 | 0.891 | 0.731 | 6.83841040173275e-111 | 1.4022160528753e-106 | PGC | 34.61 |
| 6 | S100a6 | 0.949296384477133 | 0.793 | 0.468 | 1.06474625608869e-97 | 2.18326219810986e-93 | PGC | 34.61 |
| 6 | AC160336. | 1.44692990953536 | 0.99 | 0.952 | 1.69066469663879e-95 | 3.46670796045784e-91 | PGC | 34.61 |
| 6 | Gfod2 | 0.929020675838863 | 0.489 | 0.342 | 2.05154930420491e-25 | 4.20670184827218e-21 | PGC | 34.61 |
| 7 | Ccnb2 | 0.873364341315035 | 0.992 | 0.73 | 5.27664948747395e-130 | 1.08197697740653e-125 | Mesenchyme | 36.75 |
| 7 | Cenpa | 1.01388707627452 | 1 | 0.747 | 1.00915883143108e-118 | 2.06928018384943e-114 | Mesenchyme | 36.75 |
| 7 | Cdc20 | 0.971445301867838 | 0.971 | 0.647 | 1.08394738478001e-111 | 2.22263411249142e-107 | Mesenchyme | 36.75 |
| 7 | Ube2s | 0.701578451632604 | 1 | 0.981 | 1.76080128265477e-97 | 3.61052303008361e-93 | Mesenchyme | 36.75 |
| 7 | Hmgb3 | 0.647120642810191 | 1 | 0.955 | 1.86348638649518e-94 | 3.82107883550837e-90 | Mesenchyme | 36.75 |
| 7 | Hist1h2bc | 0.647304189108544 | 0.722 | 0.289 | 2.55464954408765e-94 | 5.23830889015173e-90 | Mesenchyme | 36.75 |
| 7 | Dynll1 | 0.598487678799685 | 1 | 0.997 | 1.92199006046717e-92 | 3.94104061898794e-88 | Mesenchyme | 36.75 |
| 7 | Cdkn3 | 0.523295351059618 | 0.774 | 0.379 | 4.08145151547601e-84 | 8.36901633248357e-80 | Mesenchyme | 36.75 |
| 7 | Cdca8 | 0.649760276048885 | 0.992 | 0.841 | 3.53106543724181e-77 | 7.24044967906434e-73 | Mesenchyme | 36.75 |
| 7 | Tpx2 | 0.678055943209226 | 0.921 | 0.666 | 1.95915838490385e-70 | 4.01725426824535e-66 | Mesenchyme | 36.75 |
| 7 | H2afz | 0.531116600352985 | 1 | 0.996 | 3.21689187987425e-68 | 6.59623679968215e-64 | Mesenchyme | 36.75 |
| 7 | Ptms | 0.559488460172578 | 0.995 | 0.914 | 1.02993201986489e-62 | 2.11187560673295e-58 | Mesenchyme | 36.75 |
| 7 | Nasp | 0.533612262790625 | 1 | 0.976 | 1.70221470524342e-61 | 3.49039125310162e-57 | Mesenchyme | 36.75 |
| 7 | Ccna2 | 0.555453895688032 | 0.95 | 0.718 | 5.50341616318641e-57 | 1.12847548426137e-52 | Mesenchyme | 36.75 |
| 7 | Hsp90b1 | 0.486378150231446 | 1 | 0.99 | 7.25656267591042e-55 | 1.48795817669543e-50 | Mesenchyme | 36.75 |
| 7 | Pttg1 | 0.813090083990763 | 0.672 | 0.44 | 3.15465761329387e-46 | 6.46862543605908e-42 | Mesenchyme | 36.75 |
| 7 | Hmmr | 0.494530256877465 | 0.856 | 0.591 | 2.90276825316142e-41 | 5.95212630310748e-37 | Mesenchyme | 36.75 |
| 7 | Cd24a | 0.491092066025204 | 0.987 | 0.92 | 1.44312306476992e-37 | 2.95912384431071e-33 | Mesenchyme | 36.75 |
| 7 | Cks2 | 0.561945970869909 | 0.966 | 0.889 | 6.34123943886703e-37 | 1.30027114693968e-32 | Mesenchyme | 36.75 |
| 7 | Tuba1a | 0.517492500096597 | 0.995 | 0.939 | 2.06605328451997e-34 | 4.23644225990819e-30 | Mesenchyme | 36.75 |
| 8 | Ctla2a | 2.78830185944797 | 0.695 | 0.036 | 0 | 0 | Endothelium | 67.86 |
| 8 | Kdr | 1.69157744035261 | 0.989 | 0.184 | 0 | 0 | Endothelium | 67.86 |
| 8 | Etv2 | 1.46832105451552 | 0.904 | 0.146 | 0 | 0 | Endothelium | 67.86 |
| 8 | Flt1 | 1.13027616036896 | 0.904 | 0.118 | 0 | 0 | Endothelium | 67.86 |
| 8 | Hhex | 1.1099621452752 | 0.882 | 0.111 | 0 | 0 | Endothelium | 67.86 |
| 8 | Dusp2 | 0.919168974069204 | 0.777 | 0.103 | 0 | 0 | Endothelium | 67.86 |
| 8 | Ecscr | 1.21013241754139 | 0.816 | 0.139 | 2.03268226049415e-299 | 4.16801497514326e-295 | Endothelium | 67.86 |
| 8 | Lmo2 | 1.32289573957691 | 0.827 | 0.14 | 1.53185324816393e-289 | 3.14106508536014e-285 | Endothelium | 67.86 |
| 8 | Ramp2 | 1.91968313263899 | 0.978 | 0.314 | 5.37704444694112e-277 | 1.10256296384528e-272 | Endothelium | 67.86 |
| 8 | Egfl7 | 2.06453577332738 | 1 | 0.795 | 1.43857748630907e-208 | 2.94980313567675e-204 | Endothelium | 67.86 |
| 8 | Inka1 | 0.877068366841961 | 0.931 | 0.344 | 3.23821452087211e-195 | 6.63995887504827e-191 | Endothelium | 67.86 |
| 8 | F2r | 0.961018758512067 | 0.898 | 0.358 | 2.04355711488415e-176 | 4.19031386406995e-172 | Endothelium | 67.86 |
| 8 | Gadd45g | 1.62489874237252 | 0.984 | 0.627 | 9.05309821014283e-171 | 1.85633778798979e-166 | Endothelium | 67.86 |
| 8 | Clic1 | 0.895662925302318 | 1 | 0.989 | 1.00027433841327e-153 | 2.05106253091641e-149 | Endothelium | 67.86 |
| 8 | Itm2a | 1.18717086223139 | 0.981 | 0.686 | 3.30720767343543e-151 | 6.78142933437935e-147 | Endothelium | 67.86 |
| 8 | Creg1 | 0.964094662299 | 0.992 | 0.815 | 3.10853361825835e-141 | 6.37404818423875e-137 | Endothelium | 67.86 |
| 8 | Igfbp4 | 1.26244785450386 | 1 | 0.83 | 7.95415256354769e-137 | 1.63099898315545e-132 | Endothelium | 67.86 |
| 8 | Arpc3 | 0.940140805265325 | 1 | 0.972 | 7.88818944748261e-121 | 1.61747324620631e-116 | Endothelium | 67.86 |
| 8 | Phlda1 | 0.932222482787121 | 0.945 | 0.606 | 6.24003892924462e-107 | 1.27951998244161e-102 | Endothelium | 67.86 |
| 8 | Rspo3 | 0.938144353097125 | 0.766 | 0.313 | 7.57455501408733e-100 | 1.55316250563861e-95 | Endothelium | 67.86 |
| 9 | Hoxb5os | 0.790808539141403 | 0.642 | 0.117 | 3.49687021621556e-182 | 7.17033237835001e-178 | Pharyngeal mesoderm | 36.9 |
| 9 | Hoxaas3 | 1.51596564481989 | 0.758 | 0.186 | 2.69253393375798e-173 | 5.52104083117074e-169 | Pharyngeal mesoderm | 36.9 |
| 9 | Hoxc8 | 0.801495470195416 | 0.524 | 0.087 | 1.40245444149133e-152 | 2.87573283227797e-148 | Pharyngeal mesoderm | 36.9 |
| 9 | Hoxc9 | 0.575863940971921 | 0.442 | 0.083 | 4.07399008069598e-111 | 8.35371666046711e-107 | Pharyngeal mesoderm | 36.9 |
| 9 | Pclaf | 0.878971188519889 | 1 | 0.892 | 1.61133372523937e-106 | 3.30403980360334e-102 | Pharyngeal mesoderm | 36.9 |
| 9 | Hoxb8 | 0.682909327423344 | 0.701 | 0.25 | 3.22403022648431e-99 | 6.61087397940609e-95 | Pharyngeal mesoderm | 36.9 |
| 9 | Mdk | 0.675743417621734 | 1 | 0.997 | 8.52533153025049e-98 | 1.74811923027786e-93 | Pharyngeal mesoderm | 36.9 |
| 9 | Hoxd9 | 0.49799438673517 | 0.654 | 0.22 | 1.76962770424848e-90 | 3.6286216075615e-86 | Pharyngeal mesoderm | 36.9 |
| 9 | Hist1h2ap | 0.790034004809602 | 0.961 | 0.681 | 1.83121690247168e-80 | 3.75491025851818e-76 | Pharyngeal mesoderm | 36.9 |
| 9 | Dut | 0.538335023682877 | 1 | 0.975 | 4.04785757758072e-76 | 8.30013196282926e-72 | Pharyngeal mesoderm | 36.9 |
| 9 | H2afy | 0.573103948606653 | 0.994 | 0.943 | 7.75832520320772e-73 | 1.59084458291774e-68 | Pharyngeal mesoderm | 36.9 |
| 9 | Zfp503 | 0.519726168161546 | 0.783 | 0.392 | 2.64859709450203e-69 | 5.43094834227641e-65 | Pharyngeal mesoderm | 36.9 |
| 9 | Sms | 0.545479427255438 | 0.986 | 0.862 | 1.96380962416346e-64 | 4.02679163434717e-60 | Pharyngeal mesoderm | 36.9 |
| 9 | Rrm2 | 0.617339457661052 | 0.983 | 0.828 | 2.49296254864456e-58 | 5.11181970599568e-54 | Pharyngeal mesoderm | 36.9 |
| 9 | H2afz | 0.505289934535084 | 1 | 0.996 | 7.01678216740881e-58 | 1.43879118342718e-53 | Pharyngeal mesoderm | 36.9 |
| 9 | Hist1h1e | 0.563944248932786 | 0.893 | 0.641 | 1.44913717356967e-52 | 2.97145577440461e-48 | Pharyngeal mesoderm | 36.9 |
| 9 | Hist1h1b | 0.679778517509906 | 0.842 | 0.536 | 8.46663496871458e-51 | 1.73608350033492e-46 | Pharyngeal mesoderm | 36.9 |
| 9 | Gm45889 | 0.526881685730851 | 0.749 | 0.482 | 2.37900727656698e-40 | 4.8781544206006e-36 | Pharyngeal mesoderm | 36.9 |
| 9 | Fgf8 | 0.503049482971556 | 0.318 | 0.115 | 3.16840056396272e-32 | 6.49680535640555e-28 | Pharyngeal mesoderm | 36.9 |
| 9 | Rspo3 | 0.664053884561668 | 0.552 | 0.325 | 2.70517468267111e-25 | 5.54696068681712e-21 | Pharyngeal mesoderm | 36.9 |
| 10 | Hoxaas3 | 1.10653207667131 | 0.62 | 0.195 | 4.82692886550416e-92 | 9.89761763871628e-88 | Pharyngeal mesoderm | 38.86 |
| 10 | Hoxc8 | 0.674581910079545 | 0.44 | 0.093 | 6.38515298404808e-92 | 1.30927561937906e-87 | Pharyngeal mesoderm | 38.86 |
| 10 | Hoxb5os | 0.592036261996283 | 0.506 | 0.126 | 2.88008462389999e-91 | 5.90561352130693e-87 | Pharyngeal mesoderm | 38.86 |
| 10 | Hoxc6 | 0.412532703415374 | 0.416 | 0.086 | 1.04477231936785e-89 | 2.14230564086379e-85 | Pharyngeal mesoderm | 38.86 |
| 10 | Mdk | 0.655225573234302 | 1 | 0.997 | 8.54979467355963e-75 | 1.7531353978134e-70 | Pharyngeal mesoderm | 38.86 |
| 10 | Ccnd2 | 0.983397065959205 | 0.892 | 0.626 | 5.36375221677089e-72 | 1.09983739204887e-67 | Pharyngeal mesoderm | 38.86 |
| 10 | Hoxc9 | 0.421246770049294 | 0.355 | 0.089 | 1.22323847831854e-58 | 2.50825049979216e-54 | Pharyngeal mesoderm | 38.86 |
| 10 | Mllt3 | 0.476650356290447 | 0.614 | 0.304 | 9.9068773403241e-53 | 2.03140519863346e-48 | Pharyngeal mesoderm | 38.86 |
| 10 | Hoxb8 | 0.498735767585113 | 0.554 | 0.259 | 4.26101970356644e-46 | 8.73722090216298e-42 | Pharyngeal mesoderm | 38.86 |
| 10 | Crabp2 | 0.550049578608136 | 0.569 | 0.284 | 4.33510212634150e-43 | 8.88912691006326e-39 | Pharyngeal mesoderm | 38.86 |
| 10 | H2afy | 0.40355930991234 | 0.982 | 0.944 | 9.38274925701788e-37 | 1.92393273515152e-32 | Pharyngeal mesoderm | 38.86 |
| 10 | Fgf8 | 0.575082316283222 | 0.325 | 0.115 | 8.02632581525903e-33 | 1.64579810841886e-28 | Pharyngeal mesoderm | 38.86 |

**Supplementary Table S1:** **Differential gene expression analysis at day 5**

| Cluster | Gene | avg_logFC | pct.1 | pct.2 | p_val | p_val_adj | most expressed celltype | pct.celltype |
|---|---|---|---|---|---|---|---|---|
| 10 | Plgrkt | 0.396429587380007 | 0.834 | 0.703 | 3.14938755145023e-31 | 6.4578191742487e-27 | Pharyngeal mesoderm | 38.86 |
| 10 | Gsta4 | 0.401195417468862 | 0.973 | 0.93 | 8.56223646640536e-30 | 1.75568658743642e-25 | Pharyngeal mesoderm | 38.86 |
| 10 | Cdx2 | 0.425980516469865 | 0.572 | 0.357 | 3.4461226666355e-25 | 7.06627452793609e-21 | Pharyngeal mesoderm | 38.86 |
| 10 | Ptn | 0.399840563341853 | 0.461 | 0.296 | 2.08038383200909e-17 | 4.26582704753465e-13 | Pharyngeal mesoderm | 38.86 |
| 10 | Id1 | 0.410703231807076 | 0.967 | 0.935 | 5.45873497989983e-15 | 1.11931360762846e-10 | Pharyngeal mesoderm | 38.86 |
| 10 | Sct | 0.653463015039336 | 0.557 | 0.449 | 5.96582165499735e-12 | 1.22329173035721e-7 | Pharyngeal mesoderm | 38.86 |
| 10 | Rspo3 | 0.546930846704853 | 0.431 | 0.332 | 7.66988400884823e-8 | 0.00157270971601433 | Pharyngeal mesoderm | 38.86 |
| 10 | Mt1 | 0.421155661360876 | 0.916 | 0.892 | 0.0000396277770444482 | 0.81256756829641 | Pharyngeal mesoderm | 38.86 |
| 11 | Dppa5a | 5.42799780094868 | 0.988 | 0.16 | 0 | 0 | PGC | 89.51 |
| 11 | Mylpf | 2.07607998590899 | 0.725 | 0.062 | 0 | 0 | PGC | 89.51 |
| 11 | Utf1 | 2.03940687837349 | 0.935 | 0.022 | 0 | 0 | PGC | 89.51 |
| 11 | Pou5f1 | 1.97668806964849 | 0.963 | 0.106 | 0 | 0 | PGC | 89.51 |
| 11 | Klf2 | 1.91199739255995 | 0.781 | 0.033 | 0 | 0 | PGC | 89.51 |
| 11 | Tdh | 1.90352680248043 | 0.765 | 0.016 | 0 | 0 | PGC | 89.51 |
| 11 | Zfp42 | 1.8807540239626 | 0.88 | 0.05 | 0 | 0 | PGC | 89.51 |
| 11 | Ooep | 1.69595513136172 | 0.756 | 0.018 | 0 | 0 | PGC | 89.51 |
| 11 | Tdgf1 | 1.59160305203295 | 0.765 | 0.029 | 0 | 0 | PGC | 89.51 |
| 11 | Sox2 | 1.51849844599654 | 0.83 | 0.061 | 0 | 0 | PGC | 89.51 |
| 11 | Hspb1 | 2.24837036304512 | 0.963 | 0.341 | 3.06004773130488e-212 | 6.27462787304066e-208 | PGC | 89.51 |
| 11 | Chchd10 | 2.0547960784819 | 0.994 | 0.477 | 1.22806417241174e-207 | 2.51814558553028e-203 | PGC | 89.51 |
| 11 | Ckb | 1.70368201339729 | 0.954 | 0.345 | 2.22288821320133e-203 | 4.55803228116934e-199 | PGC | 89.51 |
| 11 | Mkrn1 | 2.50353469020417 | 0.994 | 0.755 | 9.8064251968719e-185 | 2.01080748661858e-180 | PGC | 89.51 |
| 11 | Sycp3 | 1.78773681644295 | 0.981 | 0.421 | 8.33050970619295e-176 | 1.70817101525486e-171 | PGC | 89.51 |
| 11 | Mt2 | 1.9345188486484 | 1 | 0.899 | 8.94517031411309e-156 | 1.83420717290889e-151 | PGC | 89.51 |
| 11 | Mt1 | 2.23723908609351 | 0.994 | 0.888 | 1.36589315568793e-144 | 2.80076391573809e-140 | PGC | 89.51 |
| 11 | Sdc4 | 1.59608157297066 | 0.944 | 0.519 | 1.04158994317797e-132 | 2.13578017848642e-128 | PGC | 89.51 |
| 11 | Gpx4 | 1.6551269323295 | 0.997 | 0.996 | 2.84022997830241e-121 | 5.82389157050909e-117 | PGC | 89.51 |
| 11 | Fabp3 | 1.73376656633566 | 0.852 | 0.374 | 1.81034475081371e-109 | 3.71211191154351e-105 | PGC | 89.51 |
| 12 | Ube2c | 1.63467855959385 | 1 | 0.833 | 7.03747687433514e-104 | 1.44303463308242e-99 | Intermediate mesoderm | 42.35 |
| 12 | Plk1 | 0.714285594571383 | 0.929 | 0.357 | 1.9227462132138e-103 | 3.9425911101949e-99 | Intermediate mesoderm | 42.35 |
| 12 | Hoxaas3 | 1.59844837708175 | 0.706 | 0.197 | 6.79213611716232e-101 | 1.39272751082413e-96 | Intermediate mesoderm | 42.35 |
| 12 | Nusap1 | 0.962100309002118 | 0.976 | 0.489 | 8.1761711489674e-98 | 1.67652389409577e-93 | Intermediate mesoderm | 42.35 |
| 12 | Cks2 | 1.04584980170295 | 1 | 0.889 | 1.61118272418469e-95 | 3.30373017594071e-91 | Intermediate mesoderm | 42.35 |
| 12 | Hoxc8 | 0.749267723089312 | 0.498 | 0.094 | 1.72284493512027e-95 | 3.53269353946411e-91 | Intermediate mesoderm | 42.35 |
| 12 | H2afx | 1.09557209545948 | 1 | 0.944 | 1.8054621327375e-92 | 3.70210010317825e-88 | Intermediate mesoderm | 42.35 |
| 12 | Cenpf | 1.19222076012478 | 1 | 0.726 | 2.93833224124336e-92 | 6.02505026067001e-88 | Intermediate mesoderm | 42.35 |
| 12 | Prc1 | 0.932082301348272 | 0.976 | 0.547 | 1.07275257446307e-90 | 2.19967915393652e-86 | Intermediate mesoderm | 42.35 |
| 12 | Ccnb1 | 0.938263482309749 | 0.988 | 0.648 | 3.35602745565629e-90 | 6.88153429782323e-86 | Intermediate mesoderm | 42.35 |
| 12 | Arl6ip1 | 1.07741657963623 | 1 | 0.967 | 5.75121030809488e-87 | 1.17928567367485e-82 | Intermediate mesoderm | 42.35 |
| 12 | Cenpa | 1.08061767800555 | 0.996 | 0.753 | 7.2709750481687e-86 | 1.49091343362699e-81 | Intermediate mesoderm | 42.35 |
| 12 | Tpx2 | 0.853864795344137 | 0.992 | 0.668 | 1.19031662306826e-85 | 2.44074423560146e-81 | Intermediate mesoderm | 42.35 |
| 12 | Hmmr | 0.865439613587769 | 0.988 | 0.591 | 5.47505601883939e-81 | 1.12266023666302e-76 | Intermediate mesoderm | 42.35 |
| 12 | Cdc20 | 0.927713810500256 | 0.996 | 0.653 | 7.7194435686759e-81 | 1.58287190375699e-76 | Intermediate mesoderm | 42.35 |
| 12 | Tubb4b | 0.978484641687011 | 1 | 0.951 | 2.14727101781212e-78 | 4.40297922202376e-74 | Intermediate mesoderm | 42.35 |
| 12 | Cdca8 | 0.714771994388668 | 0.996 | 0.844 | 2.71691911419383e-69 | 5.57104264365445e-65 | Intermediate mesoderm | 42.35 |
| 12 | Top2a | 0.927293617511549 | 0.988 | 0.82 | 2.23048939067585e-66 | 4.57361849558084e-62 | Intermediate mesoderm | 42.35 |
| 12 | Cdk1 | 0.864966554793533 | 0.973 | 0.805 | 2.7172896803134e-66 | 5.57180248948262e-62 | Intermediate mesoderm | 42.35 |
| 12 | Rspo3 | 0.738729204545253 | 0.58 | 0.328 | 1.32875930545569e-22 | 2.72462095583689e-18 | Intermediate mesoderm | 42.35 |
| 13 | mt-Co1 | 0.92800329282587 | 1 | 0.992 | 9.77471801051796e-66 | 2.00430592805671e-61 | Pharyngeal mesoderm | 37.42 |
| 13 | mt-Nd4 | 0.823085708251861 | 1 | 0.986 | 6.42561768370199e-53 | 1.31757290604309e-48 | Pharyngeal mesoderm | 37.42 |
| 13 | mt-Nd3 | 0.907628132619707 | 0.945 | 0.973 | 4.28010416677539e-48 | 8.77635359397294e-44 | Pharyngeal mesoderm | 37.42 |
| 13 | Prtg | 0.980514639131922 | 0.975 | 0.915 | 7.97830715364831e-46 | 1.63595188185559e-41 | Pharyngeal mesoderm | 37.42 |
| 13 | mt-Co2 | 0.668055059168927 | 1 | 0.991 | 1.44156572908644e-43 | 2.95593052749175e-39 | Pharyngeal mesoderm | 37.42 |
| 13 | mt-Atp6 | 0.694167348974372 | 1 | 0.992 | 9.29409501838973e-42 | 1.90575418352081e-37 | Pharyngeal mesoderm | 37.42 |
| 13 | Igf2 | 1.634025892713 | 0.982 | 0.869 | 1.31183727321825e-40 | 2.68992232873402e-36 | Pharyngeal mesoderm | 37.42 |
| 13 | mt-Nd1 | 0.7429623583282 | 0.988 | 0.983 | 2.74172195168507e-38 | 5.62190086193023e-34 | Pharyngeal mesoderm | 37.42 |
| 13 | mt-Nd2 | 0.663781889204966 | 0.988 | 0.984 | 2.80257509934913e-34 | 5.74668024121539e-30 | Pharyngeal mesoderm | 37.42 |
| 13 | Cbx3 | 0.752648860591101 | 0.957 | 0.969 | 5.76079136815589e-33 | 1.18125027004036e-28 | Pharyngeal mesoderm | 37.42 |
| 13 | Gm42418 | 1.74424210125918 | 1 | 1 | 1.06941254359787e-32 | 2.19283042064742e-28 | Pharyngeal mesoderm | 37.42 |
| 13 | Mid1 | 0.839126840639589 | 0.926 | 0.917 | 1.07428763130311e-29 | 2.20282678798702e-25 | Pharyngeal mesoderm | 37.42 |
| 13 | P4hb | 0.626805288903078 | 0.914 | 0.93 | 3.17821994647155e-29 | 6.51694000023991e-25 | Pharyngeal mesoderm | 37.42 |
| 13 | mt-Nd5 | 0.70080140346396 | 0.969 | 0.97 | 2.47598204968106e-28 | 5.07700119287102e-24 | Pharyngeal mesoderm | 37.42 |
| 13 | Igf2r | 0.763550423516226 | 0.926 | 0.807 | 1.1904700232629e-27 | 2.44105878270058e-23 | Pharyngeal mesoderm | 37.42 |
| 13 | Actb | 0.662064535240027 | 1 | 1 | 7.00336552806336e-24 | 1.43604010152939e-19 | Pharyngeal mesoderm | 37.42 |
| 13 | Kdm6b | 0.62715955483725 | 0.791 | 0.687 | 7.90153044208961e-19 | 1.62020881715047e-14 | Pharyngeal mesoderm | 37.42 |
| 13 | Meg3 | 0.70565662040163 | 0.969 | 0.929 | 2.97720460179727e-14 | 6.1047580359853e-10 | Pharyngeal mesoderm | 37.42 |
| 13 | Cgnl1 | 0.673216518487828 | 0.804 | 0.662 | 4.58262567475808e-13 | 9.39667394609143e-9 | Pharyngeal mesoderm | 37.42 |
| 13 | Golga4 | 0.648743340244673 | 0.663 | 0.632 | 0.000453404651691483 | 1 | Pharyngeal mesoderm | 37.42 |
| 14 | Ftl1-ps1 | 1.04439302202671 | 1 | 1 | 2.03711949552703e-33 | 4.17711352557817e-29 | Epiblast | 28.57 |
| 14 | Eif1 | 0.901320263334421 | 1 | 0.999 | 9.56718381460702e-26 | 1.96175104118517e-21 | Epiblast | 28.57 |
| 14 | Pgk1 | 1.04018485839799 | 0.988 | 0.997 | 1.25810542140048e-21 | 2.57974516658168e-17 | Epiblast | 28.57 |
| 14 | Crip2 | 1.36605909560446 | 0.798 | 0.614 | 1.12988410368682e-16 | 2.31682735460983e-12 | Epiblast | 28.57 |
| 14 | Tagln2 | 0.975736003088177 | 0.94 | 0.962 | 2.92251707452433e-16 | 5.99262126131215e-12 | Epiblast | 28.57 |
| 14 | Krt19 | 0.833013318917176 | 0.548 | 0.26 | 4.14767863350865e-14 | 8.50481503800949e-10 | Epiblast | 28.57 |
| 14 | Arrdc4 | 0.941474709971645 | 0.857 | 0.886 | 1.64036324410803e-13 | 3.36356483204352e-9 | Epiblast | 28.57 |
| 14 | AY036118 | 1.05859781162685 | 0.81 | 0.88 | 8.20433295627839e-12 | 1.68229847268488e-7 | Epiblast | 28.57 |
| 14 | Chchd10 | 0.906854057388367 | 0.69 | 0.499 | 3.2322014833359e-11 | 6.62762914158025e-7 | Epiblast | 28.57 |
| 14 | Anxa2 | 0.834645367516332 | 0.798 | 0.81 | 1.34338866470186e-10 | 0.00000275461845697117 | Epiblast | 28.57 |
| 14 | Fabp3 | 0.972167681132297 | 0.56 | 0.395 | 3.4908153618234e-8 | 0.000715791689941888 | Epiblast | 28.57 |
| 14 | Mt2 | 0.878060997643271 | 0.869 | 0.905 | 9.24422378047437e-8 | 0.00189552808618627 | Epiblast | 28.57 |
| 14 | Krt18 | 0.794330348485114 | 0.833 | 0.773 | 1.314888102943e-7 | 0.00269617805508462 | Epiblast | 28.57 |
| 14 | Txnip | 0.830798296697746 | 0.833 | 0.93 | 9.88637448518541e-7 | 0.0202720108818727 | Epiblast | 28.57 |
| 14 | Ube2c | 0.955864163926293 | 0.774 | 0.84 | 0.0000044154374503522 | 0.0905385449194719 | Epiblast | 28.57 |
| 14 | Krt8 | 0.862275199464936 | 0.845 | 0.885 | 0.0000124953514139606 | 0.256217180743262 | Epiblast | 28.57 |
| 14 | Mt1 | 1.01876586073939 | 0.833 | 0.894 | 0.0000268331909940431 | 0.550214581332855 | Epiblast | 28.57 |
| 14 | Rgcc | 1.15605294107889 | 0.25 | 0.125 | 0.000101819039984329 | 1 | Epiblast | 28.57 |
| 14 | Cystm1 | 0.874600166485528 | 0.536 | 0.477 | 0.0000703306387609154 | 1 | Epiblast | 28.57 |
| 14 | Crip1 | 0.829637723847908 | 0.524 | 0.463 | 0.000407770543008201 | 1 | Epiblast | 28.57 |

**Supplementary Table S1: Differential gene expression analysis at day 6**

| Cluster | Gene | avg_logFC | pct.1 | pct.2 | p_val | p_val_adj | most expressed celltype | pct.celltype |
|---|---|---|---|---|---|---|---|---|
| 1 | Nusap1 | 1.50958987258951 | 0.922 | 0.303 | 1.50486497322293e-260 | 2.93870031970974e-256 | Pharyngeal mesoderm | 45.67 |
| 1 | Top2a | 1.84847188249285 | 0.995 | 0.543 | 1.63924296593653e-243 | 3.20111366388086e-239 | Pharyngeal mesoderm | 45.67 |
| 1 | Prc1 | 1.33594606400337 | 0.889 | 0.296 | 1.1979653790464e-225 | 2.33938679220181e-221 | Pharyngeal mesoderm | 45.67 |
| 1 | Hmmr | 1.14901141604919 | 0.922 | 0.336 | 4.85530057880293e-215 | 9.48143097028637e-211 | Pharyngeal mesoderm | 45.67 |
| 1 | Cdk1 | 1.45582170103437 | 0.988 | 0.598 | 8.58872271727768e-215 | 1.67720577222999e-210 | Pharyngeal mesoderm | 45.67 |
| 1 | Mki67 | 1.34730634713087 | 0.958 | 0.429 | 7.16091629235965e-209 | 1.39838373357199e-204 | Pharyngeal mesoderm | 45.67 |
| 1 | Cenpf | 1.78438651847419 | 0.976 | 0.448 | 7.75185255113497e-207 | 1.51378176618564e-202 | Pharyngeal mesoderm | 45.67 |
| 1 | Smc2 | 1.10925583468028 | 0.997 | 0.766 | 1.8158109446107e-206 | 3.54591561263578e-202 | Pharyngeal mesoderm | 45.67 |
| 1 | Hmgb2 | 1.25744358500879 | 1 | 0.992 | 5.16489197101831e-204 | 1.00860010410046e-199 | Pharyngeal mesoderm | 45.67 |
| 1 | Smc4 | 1.13619932666435 | 0.991 | 0.693 | 5.69893503700093e-198 | 1.11288803402554e-193 | Pharyngeal mesoderm | 45.67 |
| 1 | Cenpe | 1.16000148730839 | 0.938 | 0.398 | 1.56380578585553e-194 | 3.05379993861868e-190 | Pharyngeal mesoderm | 45.67 |
| 1 | Birc5 | 1.20354359824945 | 0.998 | 0.617 | 3.13504825022583e-188 | 6.12212222304101e-184 | Pharyngeal mesoderm | 45.67 |
| 1 | Tpx2 | 1.22747937752608 | 0.969 | 0.5 | 1.67703030570831e-181 | 3.27490478098718e-177 | Pharyngeal mesoderm | 45.67 |
| 1 | Cks2 | 1.30402326467235 | 0.997 | 0.869 | 5.12757446132609e-174 | 1.00131274080776e-169 | Pharyngeal mesoderm | 45.67 |
| 1 | Ccnb1 | 1.16793389482729 | 0.946 | 0.453 | 3.54171295062237e-172 | 6.91625704997537e-168 | Pharyngeal mesoderm | 45.67 |
| 1 | Ube2c | 1.81875960848927 | 0.981 | 0.663 | 5.7427631860675e-166 | 1.12144679497526e-161 | Pharyngeal mesoderm | 45.67 |
| 1 | Arl6ip1 | 1.66323613510142 | 0.991 | 0.9 | 3.80672463273536e-152 | 7.43377186280562e-148 | Pharyngeal mesoderm | 45.67 |
| 1 | H2afx | 1.19387099383656 | 0.995 | 0.887 | 3.44112605810515e-141 | 6.71983096626774e-137 | Pharyngeal mesoderm | 45.67 |
| 1 | Hist1h1b | 1.22942264519142 | 0.879 | 0.458 | 4.31928318865381e-123 | 8.43469621080316e-119 | Pharyngeal mesoderm | 45.67 |
| 1 | Hist1h2ap | 1.1558179382636 | 0.912 | 0.521 | 1.27881566484278e-122 | 2.49727123030498e-118 | Pharyngeal mesoderm | 45.67 |
| 2 | Sfrp5 | 1.69730847699807 | 0.88 | 0.157 | 0 | 0 | Cardiomyocytes | 46.8 |
| 2 | Alcam | 1.35050321960589 | 0.784 | 0.186 | 5.27233725650959e-240 | 1.02958201945119e-235 | Cardiomyocytes | 46.8 |
| 2 | Gata4 | 0.961157335013805 | 0.902 | 0.318 | 3.64322084055323e-214 | 7.11448165743235e-210 | Cardiomyocytes | 46.8 |
| 2 | Mab21l2 | 1.12184245666932 | 0.816 | 0.217 | 1.32897585253982e-208 | 2.59522404483976e-204 | Cardiomyocytes | 46.8 |
| 2 | Vsnl1 | 0.983735978514078 | 0.61 | 0.107 | 7.76204554218105e-208 | 1.51577225347712e-203 | Cardiomyocytes | 46.8 |
| 2 | Sfrp1 | 1.53036493880296 | 0.99 | 0.653 | 1.07199286484521e-199 | 2.09338766646973e-195 | Cardiomyocytes | 46.8 |
| 2 | Igfbp5 | 1.7808758349628 | 0.992 | 0.717 | 4.82000113095594e-199 | 9.41249820853076e-195 | Cardiomyocytes | 46.8 |
| 2 | Gata6 | 1.30753668681038 | 0.998 | 0.7 | 8.71490251843617e-196 | 1.70184616380021e-191 | Cardiomyocytes | 46.8 |
| 2 | Map1b | 1.60073979486057 | 1 | 0.901 | 3.85219593918961e-195 | 7.52256823004948e-191 | Cardiomyocytes | 46.8 |
| 2 | Shox2 | 1.06304988193193 | 0.49 | 0.08 | 7.23928233963163e-172 | 1.41368705528326e-167 | Cardiomyocytes | 46.8 |
| 2 | Flrt3 | 1.1205818217176 | 0.882 | 0.483 | 1.22161318551306e-155 | 2.3855662286699e-151 | Cardiomyocytes | 46.8 |
| 2 | Tgfb2 | 1.08167798120078 | 0.986 | 0.758 | 2.98372155913849e-137 | 5.82661146068565e-133 | Cardiomyocytes | 46.8 |
| 2 | Asb4 | 1.2286842797425 | 0.96 | 0.619 | 3.47921372853416e-135 | 6.79420856908151e-131 | Cardiomyocytes | 46.8 |
| 2 | Lbh | 0.969046389076393 | 0.884 | 0.494 | 1.27257768470606e-131 | 2.48508970269399e-127 | Cardiomyocytes | 46.8 |
| 2 | Dsp | 1.07804955997428 | 0.836 | 0.38 | 4.29182534185394e-126 | 8.38107652757238e-122 | Cardiomyocytes | 46.8 |
| 2 | Nr2f1 | 1.23502125292889 | 0.916 | 0.601 | 1.60930559888551e-125 | 3.14265197350363e-121 | Cardiomyocytes | 46.8 |
| 2 | Maged2 | 1.03734261767075 | 0.988 | 0.924 | 3.62422486291494e-122 | 7.07738631230029e-118 | Cardiomyocytes | 46.8 |
| 2 | Tnni1 | 0.940804308450369 | 0.97 | 0.626 | 4.50977507648295e-110 | 8.80668876935591e-106 | Cardiomyocytes | 46.8 |
| 2 | Prnp | 0.946592567152215 | 0.886 | 0.549 | 5.54619843579653e-108 | 1.08306163054235e-103 | Cardiomyocytes | 46.8 |
| 2 | Bmp4 | 0.953906206264246 | 0.73 | 0.387 | 7.18913119972095e-88 | 1.40389354068151e-83 | Cardiomyocytes | 46.8 |
| 3 | Gnas | 0.985286019564628 | 1 | 0.999 | 1.97433113020421e-163 | 3.85547383106278e-159 | Pharyngeal mesoderm | 62.14 |
| 3 | Mdk | 0.71765388666016 | 1 | 0.999 | 1.46573572725422e-149 | 2.86228872818204e-145 | Pharyngeal mesoderm | 62.14 |
| 3 | Ntrk2 | 0.98753573709074 | 0.806 | 0.303 | 4.19854392921848e-146 | 8.19891658497785e-142 | Pharyngeal mesoderm | 62.14 |
| 3 | Alx1 | 0.957247421021133 | 0.92 | 0.487 | 1.81515857640404e-130 | 3.54464166800181e-126 | Pharyngeal mesoderm | 62.14 |
| 3 | Rgs5 | 1.36644134461925 | 0.846 | 0.425 | 3.79983201603268e-120 | 7.42031196090862e-116 | Pharyngeal mesoderm | 62.14 |
| 3 | Mfap4 | 0.794439422668385 | 0.938 | 0.635 | 9.66096379935669e-109 | 1.88659301073837e-104 | Pharyngeal mesoderm | 62.14 |
| 3 | Dlk1 | 1.0265804189724 | 1 | 0.83 | 5.24860864024021e-93 | 1.02494829526611e-88 | Pharyngeal mesoderm | 62.14 |
| 3 | Hand2 | 0.913575265607955 | 0.971 | 0.661 | 3.83059308564147e-92 | 7.48038217764067e-88 | Pharyngeal mesoderm | 62.14 |
| 3 | Hand2os1 | 0.815506188795439 | 0.898 | 0.541 | 2.28672386333505e-80 | 4.46551436032068e-76 | Pharyngeal mesoderm | 62.14 |
| 3 | Meis2 | 0.754244710482499 | 0.989 | 0.899 | 5.32234982214723e-79 | 1.03934847326891e-74 | Pharyngeal mesoderm | 62.14 |
| 3 | Gpc3 | 0.753909494264933 | 1 | 0.921 | 2.45707757300705e-78 | 4.79818108456817e-74 | Pharyngeal mesoderm | 62.14 |
| 3 | Armh4 | 0.749032755996355 | 0.739 | 0.394 | 1.07837193807408e-75 | 2.10584472067106e-71 | Pharyngeal mesoderm | 62.14 |
| 3 | Itm2a | 0.858439591572019 | 0.947 | 0.761 | 3.18921634592482e-68 | 6.22790168032198e-64 | Pharyngeal mesoderm | 62.14 |
| 3 | Mpped2 | 0.672813147792115 | 0.891 | 0.711 | 1.15297608433428e-59 | 2.25153169748798e-55 | Pharyngeal mesoderm | 62.14 |
| 3 | Col9a1 | 0.662295813690952 | 0.673 | 0.382 | 4.45114819049104e-59 | 8.69220218639089e-55 | Pharyngeal mesoderm | 62.14 |
| 3 | Tac2 | 0.819460301903548 | 0.461 | 0.178 | 1.24936035782174e-57 | 2.43975090675429e-53 | Pharyngeal mesoderm | 62.14 |
| 3 | Phox2a | 0.803897789388211 | 0.343 | 0.107 | 3.77019698736088e-53 | 7.36244067691832e-49 | Pharyngeal mesoderm | 62.14 |
| 3 | Mafb | 0.722071512641183 | 0.639 | 0.381 | 2.54353456127853e-51 | 4.96701429126472e-47 | Pharyngeal mesoderm | 62.14 |
| 3 | Meg3 | 0.717932213684206 | 0.982 | 0.938 | 5.97923592834152e-50 | 1.16762519208653e-45 | Pharyngeal mesoderm | 62.14 |
| 3 | Rarres2 | 0.672103477016426 | 0.528 | 0.327 | 5.31615484201959e-30 | 1.03813871754959e-25 | Pharyngeal mesoderm | 62.14 |
| 4 | Sox2 | 1.77923637183326 | 0.963 | 0.074 | 0 | 0 | Spinal cord | 54.76 |
| 4 | Fam181b | 1.17875762124832 | 0.742 | 0.096 | 0 | 0 | Spinal cord | 54.76 |
| 4 | Hes5 | 1.15864638328004 | 0.684 | 0.034 | 0 | 0 | Spinal cord | 54.76 |
| 4 | Pantr1 | 1.12054894788762 | 0.696 | 0.048 | 0 | 0 | Spinal cord | 54.76 |
| 4 | Crabp2 | 2.32191476156905 | 0.993 | 0.589 | 1.25335738045482e-230 | 2.44755629255218e-226 | Spinal cord | 54.76 |
| 4 | Fabp7 | 1.7444438345646 | 0.638 | 0.096 | 1.54990775294478e-230 | 3.02665985995057e-226 | Spinal cord | 54.76 |
| 4 | Hoxb9 | 1.12682882437737 | 0.937 | 0.261 | 5.55528850602943e-210 | 1.08483673945743e-205 | Spinal cord | 54.76 |
| 4 | Hoxb5os | 1.29726766021597 | 0.961 | 0.351 | 3.2949207245739e-202 | 6.43432119094792e-198 | Spinal cord | 54.76 |
| 4 | Ccnd1 | 1.67151420575612 | 0.977 | 0.483 | 6.66053106529455e-201 | 1.30066850643072e-196 | Spinal cord | 54.76 |
| 4 | Hoxc10 | 1.23450312298205 | 0.601 | 0.1 | 1.45914636342598e-194 | 2.84942101849825e-190 | Spinal cord | 54.76 |
| 4 | Sfrp2 | 1.30958143711102 | 0.919 | 0.366 | 3.68075359667007e-185 | 7.18777562357732e-181 | Spinal cord | 54.76 |
| 4 | Mkrn1 | 0.992341013189995 | 0.935 | 0.558 | 6.03938829229806e-158 | 1.17937174571997e-153 | Spinal cord | 54.76 |
| 4 | Tubb2b | 1.5502385045806 | 0.991 | 0.695 | 1.63082617351916e-150 | 3.18467735164822e-146 | Spinal cord | 54.76 |
| 4 | Dek | 1.00783055355707 | 0.986 | 0.925 | 6.44926357167414e-146 | 1.25941219027653e-141 | Spinal cord | 54.76 |
| 4 | Phgdh | 0.97817461618206 | 0.984 | 0.697 | 6.36056915523834e-130 | 1.24209194463494e-125 | Spinal cord | 54.76 |
| 4 | Rrm2 | 1.02866868742629 | 0.97 | 0.619 | 1.9920431146719e-125 | 3.89006179433128e-121 | Spinal cord | 54.76 |
| 4 | Cdca8 | 1.01926649578505 | 0.949 | 0.603 | 7.56773863330777e-109 | 1.47782800031234e-104 | Spinal cord | 54.76 |
| 4 | Ccnd2 | 1.34752855036988 | 0.961 | 0.796 | 2.74746166331286e-90 | 5.36524313611736e-86 | Spinal cord | 54.76 |
| 4 | Crabp1 | 1.39659932466318 | 0.731 | 0.459 | 3.43342951821005e-39 | 6.70480116316059e-35 | Spinal cord | 54.76 |
| 4 | Ckb | 0.991165021871008 | 0.803 | 0.564 | 7.38243567853357e-29 | 1.44164203930403e-24 | Spinal cord | 54.76 |
| 5 | Nusap1 | 1.29682670952784 | 0.985 | 0.32 | 2.02234039658973e-208 | 3.94922632646043e-204 | Pharyngeal mesoderm | 49.5 |
| 5 | Prc1 | 1.18366064602001 | 0.978 | 0.31 | 3.68591817931126e-204 | 7.19786102055903e-200 | Pharyngeal mesoderm | 49.5 |
| 5 | Hmmr | 1.11980550516148 | 0.99 | 0.352 | 4.7436437268886e-193 | 9.26338746986806e-189 | Pharyngeal mesoderm | 49.5 |
| 5 | Aurkb | 0.99248931721549 | 0.968 | 0.355 | 7.50053411269543e-180 | 1.46470430152716e-175 | Pharyngeal mesoderm | 49.5 |
| 5 | Ccnb1 | 1.29288318390819 | 0.995 | 0.466 | 2.88882652546484e-178 | 5.64130043892775e-174 | Pharyngeal mesoderm | 49.5 |
| 5 | Cenpf | 1.5451993776537 | 0.995 | 0.465 | 1.50546531205837e-173 | 2.93987266138758e-169 | Pharyngeal mesoderm | 49.5 |
| 5 | Cenpe | 1.11512491530364 | 0.98 | 0.414 | 1.77554479557805e-172 | 3.46728387680482e-168 | Pharyngeal mesoderm | 49.5 |
| 5 | Ube2c | 1.88982748880623 | 1 | 0.673 | 8.85123858629558e-171 | 1.7284698711318e-166 | Pharyngeal mesoderm | 49.5 |
| 5 | Tpx2 | 1.21006981600148 | 0.995 | 0.515 | 1.41161727679162e-157 | 2.75660621811868e-153 | Pharyngeal mesoderm | 49.5 |
| 5 | Cdk1 | 1.31400187237429 | 0.998 | 0.611 | 8.47866170881125e-157 | 1.65571305849666e-152 | Pharyngeal mesoderm | 49.5 |
| 5 | Top2a | 1.49454104573175 | 0.995 | 0.56 | 8.73460073400932e-157 | 1.70569283133734e-152 | Pharyngeal mesoderm | 49.5 |
| 5 | Cks2 | 1.30553177410847 | 1 | 0.874 | 1.06594363048409e-153 | 2.08157472160933e-149 | Pharyngeal mesoderm | 49.5 |
| 5 | Cdc20 | 1.18487783765552 | 0.978 | 0.462 | 1.89642957607043e-147 | 3.70334767615034e-143 | Pharyngeal mesoderm | 49.5 |
| 5 | Mki67 | 1.0485173781429 | 0.995 | 0.445 | 8.86320734959456e-147 | 1.73080713122883e-142 | Pharyngeal mesoderm | 49.5 |
| 5 | Arl6ip1 | 1.2976486284101 | 0.995 | 0.903 | 2.35315710774748e-137 | 4.59524520000927e-133 | Pharyngeal mesoderm | 49.5 |
| 5 | Smc4 | 0.980336290440971 | 0.995 | 0.704 | 9.03206404570498e-136 | 1.76378146684527e-131 | Pharyngeal mesoderm | 49.5 |
| 5 | Birc5 | 1.05577761644545 | 1 | 0.631 | 1.12850164530642e-131 | 2.20373801295438e-127 | Pharyngeal mesoderm | 49.5 |
| 5 | H2afx | 1.09982459154277 | 1 | 0.891 | 1.88180240997958e-119 | 3.67478374620813e-115 | Pharyngeal mesoderm | 49.5 |
| 5 | Cenpa | 1.05616066304989 | 1 | 0.728 | 3.03281283966094e-109 | 5.92247691328987e-105 | Pharyngeal mesoderm | 49.5 |
| 5 | Tubb4b | 1.07512928067453 | 1 | 0.846 | 1.07760897936312e-107 | 2.10435481490031e-103 | Pharyngeal mesoderm | 49.5 |
| 6 | Tnnt2 | 4.40641454642149 | 1 | 0.32 | 0 | 0 | Cardiomyocytes | 100 |
| 6 | Tnnc1 | 3.71253752564115 | 0.901 | 0.19 | 0 | 0 | Cardiomyocytes | 100 |
| 6 | Sh3bgr | 3.14304216863572 | 0.967 | 0.212 | 0 | 0 | Cardiomyocytes | 100 |
| 6 | Myh7 | 2.42657845424798 | 0.779 | 0.057 | 0 | 0 | Cardiomyocytes | 100 |
| 6 | Myh6 | 2.05597451909249 | 0.698 | 0.05 | 0 | 0 | Cardiomyocytes | 100 |
| 6 | Fbxl22 | 1.9686185796954 | 0.865 | 0.115 | 0 | 0 | Cardiomyocytes | 100 |
| 6 | Ttn | 3.89136296633733 | 0.99 | 0.288 | 2.49607536790738e-305 | 4.87433597844953e-301 | Cardiomyocytes | 100 |
| 6 | Nebl | 1.96546819754461 | 0.964 | 0.274 | 4.10599255983997e-283 | 8.01818227085549e-279 | Cardiomyocytes | 100 |

**Supplementary Table S1:** Differential gene expression analysis at day 6

| Cluster | Gene | avg_logFC | pct.1 | pct.2 | p_val | p_val_adj | most expressed celltype | pct.celltype |
|---|---|---|---|---|---|---|---|---|
| 6 | Pgam2 | 2.19127265348548 | 0.832 | 0.158 | 2.8133381433314e-280 | 5.49388672629755e-276 | Cardiomyocytes | 100 |
| 6 | Myl9 | 2.11426931133517 | 0.759 | 0.128 | 1.38156613714354e-270 | 2.69792235261391e-266 | Cardiomyocytes | 100 |
| 6 | Acta2 | 4.33659479028425 | 0.995 | 0.424 | 4.17541546980038e-268 | 8.15375132942618e-264 | Cardiomyocytes | 100 |
| 6 | Actc1 | 5.04176999074758 | 0.949 | 0.32 | 1.40189829225741e-264 | 2.73762698512027e-260 | Cardiomyocytes | 100 |
| 6 | Myl7 | 4.92360917123756 | 0.997 | 0.519 | 2.08249563672072e-247 | 4.06669747938823e-243 | Cardiomyocytes | 100 |
| 6 | Myl4 | 4.17029908726967 | 0.944 | 0.344 | 4.00767418279692e-243 | 7.82618614416583e-239 | Cardiomyocytes | 100 |
| 6 | Tnni1 | 2.77126245822487 | 0.997 | 0.631 | 3.09146826788107e-235 | 6.03701923351816e-231 | Cardiomyocytes | 100 |
| 6 | Tpm1 | 3.32328560989599 | 1 | 0.967 | 4.06688771312241e-229 | 7.94181832618544e-225 | Cardiomyocytes | 100 |
| 6 | Gyg | 2.88230803835281 | 1 | 0.739 | 7.50078159255847e-221 | 1.46475262939482e-216 | Cardiomyocytes | 100 |
| 6 | Myl3 | 3.48951976924835 | 0.642 | 0.109 | 1.1415215898982e-216 | 2.2291633607532e-212 | Cardiomyocytes | 100 |
| 6 | Ckb | 2.06587142713026 | 0.957 | 0.553 | 3.23505929259025e-168 | 6.31742378657024e-164 | Cardiomyocytes | 100 |
| 6 | Tnni3 | 1.93446968151809 | 0.452 | 0.067 | 2.26470492205904e-153 | 4.42251577179688e-149 | Cardiomyocytes | 100 |
| 7 | Ctla2a | 3.01117606476469 | 0.957 | 0.031 | 0 | 0 | Endothelium | 99.2 |
| 7 | Ecscr | 2.42297106045817 | 0.989 | 0.063 | 0 | 0 | Endothelium | 99.2 |
| 7 | Plvap | 2.24436794884652 | 0.989 | 0.154 | 0 | 0 | Endothelium | 99.2 |
| 7 | Cdh5 | 2.16690569851127 | 0.962 | 0.018 | 0 | 0 | Endothelium | 99.2 |
| 7 | Cldn5 | 2.12160827306506 | 0.874 | 0.015 | 0 | 0 | Endothelium | 99.2 |
| 7 | Kdr | 1.96065840746545 | 0.979 | 0.244 | 0 | 0 | Endothelium | 99.2 |
| 7 | Gmfg | 1.85060523585904 | 0.936 | 0.066 | 0 | 0 | Endothelium | 99.2 |
| 7 | Icam2 | 1.78002577199273 | 0.971 | 0.011 | 0 | 0 | Endothelium | 99.2 |
| 7 | Flt1 | 1.77337361040594 | 0.954 | 0.04 | 0 | 0 | Endothelium | 99.2 |
| 7 | Igf1 | 1.7363017209616 | 0.941 | 0.031 | 0 | 0 | Endothelium | 99.2 |
| 7 | Emcn | 1.71625638518595 | 0.893 | 0.038 | 0 | 0 | Endothelium | 99.2 |
| 7 | Esam | 1.56285151773414 | 0.976 | 0.014 | 0 | 0 | Endothelium | 99.2 |
| 7 | Cavin3 | 1.5962404012775 | 0.874 | 0.167 | 4.67070696812615e-292 | 9.12095656735674e-288 | Endothelium | 99.2 |
| 7 | Ramp2 | 2.98663844680421 | 1 | 0.485 | 1.336220820549e-249 | 2.60937201836809e-245 | Endothelium | 99.2 |
| 7 | Egfl7 | 2.19405041657776 | 1 | 0.657 | 9.67659829586807e-230 | 1.88964611521712e-225 | Endothelium | 99.2 |
| 7 | Hapln1 | 1.77001477143336 | 0.909 | 0.236 | 1.13498399958642e-224 | 2.21639675439235e-220 | Endothelium | 99.2 |
| 7 | Tagln2 | 2.07182677520706 | 0.995 | 0.552 | 2.49092166165435e-208 | 4.86427182087861e-204 | Endothelium | 99.2 |
| 7 | Vim | 1.68082329031116 | 1 | 0.982 | 6.49546655655356e-182 | 1.26843470916413e-177 | Endothelium | 99.2 |
| 7 | Col18a1 | 1.65423216535183 | 0.984 | 0.638 | 2.46722129502349e-175 | 4.81798974492188e-171 | Endothelium | 99.2 |
| 7 | Apoe | 2.11369391677664 | 0.995 | 0.758 | 5.3537538627971e-155 | 1.04548105432702e-150 | Endothelium | 99.2 |
| 8 | Rprm | 1.54329113934766 | 0.892 | 0.428 | 3.27126689101162e-120 | 6.38812998476749e-116 | Pharyngeal mesoderm | 82.1 |
| 8 | 2610528A1 | 1.58231525845948 | 0.767 | 0.276 | 7.71223867955453e-113 | 1.50604596934341e-108 | Pharyngeal mesoderm | 82.1 |
| 8 | Gm21814 | 0.590359788509692 | 0.648 | 0.281 | 3.88190754255599e-66 | 7.58058904910334e-62 | Pharyngeal mesoderm | 82.1 |
| 8 | Ifitm2 | 0.613157503786461 | 1 | 0.989 | 1.29332497862665e-63 | 2.52560501826212e-59 | Pharyngeal mesoderm | 82.1 |
| 8 | Peg3 | 0.847303198157004 | 0.986 | 0.967 | 1.20122522645829e-59 | 2.34575262222775e-55 | Pharyngeal mesoderm | 82.1 |
| 8 | Col3a1 | 0.58401006523185 | 0.773 | 0.381 | 4.34418792406971e-55 | 8.48333017812332e-51 | Pharyngeal mesoderm | 82.1 |
| 8 | Tshz2 | 0.649930867450891 | 0.918 | 0.731 | 2.94470440375062e-49 | 5.75041875964421e-45 | Pharyngeal mesoderm | 82.1 |
| 8 | Alx1 | 0.546827180002565 | 0.855 | 0.5 | 2.34476589732432e-48 | 4.57885884429493e-44 | Pharyngeal mesoderm | 82.1 |
| 8 | Igdcc3 | 0.539107423642701 | 0.918 | 0.728 | 2.06767380208798e-47 | 4.0377534007174e-43 | Pharyngeal mesoderm | 82.1 |
| 8 | Pbx1 | 0.521714285161151 | 0.969 | 0.932 | 3.32568479031414e-39 | 6.49439725852544e-35 | Pharyngeal mesoderm | 82.1 |
| 8 | Abcg2 | 0.545268411141236 | 0.912 | 0.806 | 6.29362145599777e-39 | 1.22901839792724e-34 | Pharyngeal mesoderm | 82.1 |
| 8 | Gm49708 | 0.536339367252293 | 0.972 | 0.844 | 6.00413911525952e-38 | 1.17248828642788e-33 | Pharyngeal mesoderm | 82.1 |
| 8 | Dlk1 | 0.661241531912728 | 0.983 | 0.835 | 3.64248203419475e-34 | 7.1130389163755e-30 | Pharyngeal mesoderm | 82.1 |
| 8 | Aldoc | 0.525946382301109 | 0.699 | 0.458 | 6.7941596837571e-34 | 1.32676350304409e-29 | Pharyngeal mesoderm | 82.1 |
| 8 | Crabp1 | 0.582431808120141 | 0.724 | 0.464 | 3.47853960386847e-30 | 6.79289213843434e-26 | Pharyngeal mesoderm | 82.1 |
| 8 | Resf1 | 0.485017656919163 | 0.943 | 0.883 | 7.30779936672808e-30 | 1.42706706033466e-25 | Pharyngeal mesoderm | 82.1 |
| 8 | Ptn | 0.955519911549686 | 0.946 | 0.845 | 1.54065800691384e-29 | 3.00859695590135e-25 | Pharyngeal mesoderm | 82.1 |
| 8 | Pdgfra | 0.488889832087378 | 0.767 | 0.519 | 4.53899313488397e-29 | 8.86374579380142e-25 | Pharyngeal mesoderm | 82.1 |
| 8 | Pmp22 | 0.64836272927172 | 0.707 | 0.508 | 5.76925863718799e-26 | 1.12662082667007e-21 | Pharyngeal mesoderm | 82.1 |
| 8 | Ifitm1 | 0.612126898208974 | 0.653 | 0.424 | 3.53078703468328e-24 | 6.89492092132951e-20 | Pharyngeal mesoderm | 82.1 |
| 9 | Osr1 | 0.834225471382845 | 0.605 | 0.172 | 5.30834060249831e-97 | 1.03661275285587e-92 | Paraxial mesoderm | 67.15 |
| 9 | Espn | 0.75810509352288 | 0.593 | 0.2 | 1.64328976671177e-81 | 3.20901625643474e-77 | Paraxial mesoderm | 67.15 |
| 9 | Hoxb8 | 0.66126353061541 | 0.741 | 0.345 | 1.30567550636929e-74 | 2.54972312883796e-70 | Paraxial mesoderm | 67.15 |
| 9 | Hoxb5os | 0.908363087188317 | 0.802 | 0.372 | 2.48434599265174e-74 | 4.85143085445031e-70 | Paraxial mesoderm | 67.15 |
| 9 | Meox1 | 0.795095134543899 | 0.645 | 0.251 | 1.12403337439023e-73 | 2.19501237350923e-69 | Paraxial mesoderm | 67.15 |
| 9 | Hoxb9 | 0.742896795842443 | 0.68 | 0.291 | 6.3385407228006e-60 | 1.2377902323485e-55 | Paraxial mesoderm | 67.15 |
| 9 | 2410006H1 | 0.669122519451757 | 1 | 0.971 | 2.10739251129995e-56 | 4.11531609606653e-52 | Paraxial mesoderm | 67.15 |
| 9 | Ncam1 | 0.812812253973863 | 0.892 | 0.667 | 2.48172352360176e-53 | 4.84630969688952e-49 | Paraxial mesoderm | 67.15 |
| 9 | Hoxc6 | 0.600886674117 | 0.651 | 0.306 | 1.04443911130038e-48 | 2.03958069654739e-44 | Paraxial mesoderm | 67.15 |
| 9 | Ifi30 | 0.663602774862053 | 0.73 | 0.489 | 8.35034832213646e-44 | 1.63065602034681e-39 | Paraxial mesoderm | 67.15 |
| 9 | Gas1 | 0.596138789760252 | 0.869 | 0.701 | 4.89219083485727e-40 | 9.55347026230927e-36 | Paraxial mesoderm | 67.15 |
| 9 | Arg1 | 0.669592845206136 | 0.526 | 0.247 | 3.92736248389243e-39 | 7.66935345854513e-35 | Paraxial mesoderm | 67.15 |
| 9 | Rdh10 | 0.651532513259572 | 0.584 | 0.313 | 1.06579545916311e-38 | 2.08128537265371e-34 | Paraxial mesoderm | 67.15 |
| 9 | Sfrp2 | 0.985106136508158 | 0.654 | 0.394 | 3.78995333320275e-36 | 7.40102086907833e-32 | Paraxial mesoderm | 67.15 |
| 9 | C1qtnf12 | 0.673390866977702 | 0.381 | 0.16 | 6.89307047872954e-35 | 1.34607880308631e-30 | Paraxial mesoderm | 67.15 |
| 9 | Fxyd6 | 0.623291840847122 | 0.919 | 0.866 | 7.54310043109159e-34 | 1.47301665218357e-29 | Paraxial mesoderm | 67.15 |
| 9 | Mest | 0.682141558166981 | 0.91 | 0.833 | 9.29813023452268e-29 | 1.81573887219759e-24 | Paraxial mesoderm | 67.15 |
| 9 | Wfdc2 | 0.755267415508815 | 0.285 | 0.106 | 4.82863575497271e-28 | 9.4293599023107e-24 | Paraxial mesoderm | 67.15 |
| 9 | Rbp1 | 0.792287070851712 | 0.858 | 0.745 | 1.00475083277267e-24 | 1.96207742623847e-20 | Paraxial mesoderm | 67.15 |
| 9 | 2610528A1 | 0.712893020486933 | 0.497 | 0.296 | 3.16061485555343e-18 | 6.17204868992473e-14 | Paraxial mesoderm | 67.15 |
| 10 | Col3a1 | 1.20173732867526 | 0.873 | 0.376 | 3.66203148344764e-112 | 7.15121508087654e-108 | Mesenchyme | 39.35 |
| 10 | Vim | 1.09333878956371 | 1 | 0.982 | 1.59202057392506e-96 | 3.10889777676086e-92 | Mesenchyme | 39.35 |
| 10 | Hsd11b2 | 0.914722899051287 | 0.808 | 0.367 | 1.48986300676173e-88 | 2.9094044796043e-84 | Mesenchyme | 39.35 |
| 10 | Hand1 | 1.09669386591444 | 0.828 | 0.388 | 5.83830575700849e-83 | 1.14010434822862e-78 | Mesenchyme | 39.35 |
| 10 | Igf2 | 1.03326391986152 | 1 | 0.937 | 5.23642703747623e-82 | 1.02256947187836e-77 | Mesenchyme | 39.35 |
| 10 | Fn1 | 0.968035507342969 | 0.982 | 0.862 | 9.92080727953488e-82 | 1.93733524554757e-77 | Mesenchyme | 39.35 |
| 10 | Arhgdib | 0.648454609909979 | 0.796 | 0.385 | 1.90156357047308e-74 | 3.71337334041984e-70 | Mesenchyme | 39.35 |
| 10 | Phlda2 | 1.50847460830835 | 0.562 | 0.196 | 8.58207047880582e-74 | 1.6759067231012e-69 | Mesenchyme | 39.35 |
| 10 | Lgals1 | 1.04141642331468 | 0.976 | 0.823 | 1.09206045096706e-69 | 2.13257564864847e-65 | Mesenchyme | 39.35 |
| 10 | Cdkn1c | 1.03578392754014 | 1 | 0.984 | 9.32204355279977e-66 | 1.82040866499074e-61 | Mesenchyme | 39.35 |
| 10 | Hapln1 | 0.851398698532031 | 0.636 | 0.259 | 9.68808198263921e-62 | 1.89188864956979e-57 | Mesenchyme | 39.35 |
| 10 | Sparc | 0.739680279484444 | 0.997 | 0.827 | 6.38759744769483e-59 | 1.24737002958585e-54 | Mesenchyme | 39.35 |
| 10 | S100a10 | 0.87054099225873 | 0.891 | 0.683 | 8.27025817923075e-59 | 1.61501601724018e-54 | Mesenchyme | 39.35 |
| 10 | Itm2a | 0.924888741185073 | 0.967 | 0.764 | 1.40571863915151e-57 | 2.74508735853507e-53 | Mesenchyme | 39.35 |
| 10 | Capn6 | 0.693394182755328 | 0.885 | 0.543 | 5.2654360049061e-56 | 1.02823434303806e-51 | Mesenchyme | 39.35 |
| 10 | Apoe | 0.742076889458165 | 0.982 | 0.761 | 1.56720553327743e-54 | 3.06043896538416e-50 | Mesenchyme | 39.35 |
| 10 | Dlk1 | 0.827509336019772 | 1 | 0.834 | 1.10884336593122e-52 | 2.16534932499049e-48 | Mesenchyme | 39.35 |
| 10 | Dkk1 | 0.705346782066759 | 0.459 | 0.19 | 1.06046905562856e-37 | 2.07088397183146e-33 | Mesenchyme | 39.35 |
| 10 | Asb4 | 0.701397766122193 | 0.896 | 0.634 | 1.07286790805104e-37 | 2.09509645084208e-33 | Mesenchyme | 39.35 |
| 10 | Krt8 | 0.811472043894554 | 0.553 | 0.337 | 4.0258540470394e-20 | 7.86168778305855e-16 | Mesenchyme | 39.35 |
| 11 | Ccnb2 | 1.20054507036979 | 0.983 | 0.572 | 3.19190962281925e-115 | 6.23316111144144e-111 | Pharyngeal mesoderm | 62.02 |
| 11 | Cenpa | 1.36643249649383 | 1 | 0.734 | 2.73000420648479e-109 | 5.33115221442349e-105 | Pharyngeal mesoderm | 62.02 |
| 11 | Dynll1 | 0.784336120928883 | 1 | 0.999 | 2.31714477986738e-80 | 4.52492032612501e-76 | Pharyngeal mesoderm | 62.02 |
| 11 | Pttg1 | 1.28966796672947 | 0.875 | 0.524 | 3.59605602599818e-80 | 7.02237820756925e-76 | Pharyngeal mesoderm | 62.02 |
| 11 | H2afv | 0.622627234971736 | 1 | 1 | 5.88095430239611e-72 | 1.14843275617191e-67 | Pharyngeal mesoderm | 62.02 |
| 11 | Hmgb2 | 0.929575273161726 | 1 | 0.992 | 4.5046196895173e-68 | 8.79662132968938e-64 | Pharyngeal mesoderm | 62.02 |
| 11 | Hmgn2 | 0.642485316836034 | 0.997 | 0.992 | 1.74521981831915e-66 | 3.40806526121363e-62 | Pharyngeal mesoderm | 62.02 |
| 11 | Ube2s | 0.750127033760485 | 1 | 0.985 | 4.59928873898265e-64 | 8.98149104948531e-60 | Pharyngeal mesoderm | 62.02 |
| 11 | Tuba1a | 0.77143094208425 | 1 | 0.987 | 5.28820248529929e-60 | 1.03268018132925e-55 | Pharyngeal mesoderm | 62.02 |
| 11 | Ptms | 0.646141089016583 | 1 | 0.986 | 9.64400644200624e-60 | 1.88328157799498e-55 | Pharyngeal mesoderm | 62.02 |
| 11 | H2afx | 0.770260113517744 | 1 | 0.893 | 8.04574962826407e-59 | 1.57117398740741e-54 | Pharyngeal mesoderm | 62.02 |
| 11 | Birc5 | 0.873945830621653 | 0.976 | 0.641 | 3.07207134802109e-57 | 5.99914092841558e-53 | Pharyngeal mesoderm | 62.02 |
| 11 | H1fx | 0.696478781207957 | 0.997 | 0.929 | 2.10368185085736e-54 | 4.10806991835425e-50 | Pharyngeal mesoderm | 62.02 |
| 11 | Cdc20 | 0.793456118730912 | 0.808 | 0.484 | 3.89445766820529e-44 | 7.6050969344713e-40 | Pharyngeal mesoderm | 62.02 |
| 11 | Cdca3 | 0.617887336314545 | 0.916 | 0.599 | 1.00453360920406e-43 | 1.96165323205369e-39 | Pharyngeal mesoderm | 62.02 |
| 11 | Tpx2 | 0.689941042537628 | 0.864 | 0.534 | 1.00135579945092e-38 | 1.95544760516776e-34 | Pharyngeal mesoderm | 62.02 |

**<u>Supplementary Table S1:</u> Differential gene expression analysis at day 6**

| Cluster | Gene | avg_logFC | pct.1 | pct.2 | p_val | p_val_adj | most expressed celltype | pct.celltype |
|---|---|---|---|---|---|---|---|---|
| 11 | Tubb4b | 0.658515451182894 | 0.969 | 0.852 | 1.07932305977477e-34 | 2.10770207112818e-30 | Pharyngeal mesoderm | 62.02 |
| 11 | Dlk1 | 0.591391673320079 | 0.993 | 0.836 | 3.80231157534587e-29 | 7.42515404433541e-25 | Pharyngeal mesoderm | 62.02 |
| 11 | Ptn | 0.641064336175133 | 0.99 | 0.844 | 5.76375455540121e-26 | 1.12554598957875e-21 | Pharyngeal mesoderm | 62.02 |
| 11 | Cks2 | 0.581696782682361 | 0.944 | 0.88 | 1.88492960973637e-25 | 3.68089054189319e-21 | Pharyngeal mesoderm | 62.02 |
| 12 | Ran | 0.690283265281709 | 1 | 0.999 | 2.01446732366876e-89 | 3.93385178966036e-85 | Paraxial mesoderm | 46.29 |
| 12 | Fabp5 | 0.897090195299366 | 1 | 0.963 | 1.65011549645254e-74 | 3.22234554147251e-70 | Paraxial mesoderm | 46.29 |
| 12 | Meox1 | 0.881403033129017 | 0.7 | 0.253 | 1.06449422129285e-71 | 2.07874431534068e-67 | Paraxial mesoderm | 46.29 |
| 12 | Hspe1 | 0.682076364851311 | 1 | 0.998 | 4.72823609779768e-71 | 9.23329945177931e-67 | Paraxial mesoderm | 46.29 |
| 12 | Eif5a | 0.685451878813456 | 1 | 0.998 | 2.11403215155778e-68 | 4.12828198556203e-64 | Paraxial mesoderm | 46.29 |
| 12 | Nme1 | 0.705561326869276 | 1 | 0.987 | 1.18965440030186e-63 | 2.32315711290947e-59 | Paraxial mesoderm | 46.29 |
| 12 | H2afz | 0.612889379371228 | 1 | 1 | 7.60727150994751e-61 | 1.48554798046255e-56 | Paraxial mesoderm | 46.29 |
| 12 | Cacybp | 0.659701926016483 | 0.996 | 0.973 | 3.69263405570217e-60 | 7.21097578397519e-56 | Paraxial mesoderm | 46.29 |
| 12 | Hspd1 | 0.724191578166468 | 1 | 0.99 | 2.24546598612512e-59 | 4.38494597770514e-55 | Paraxial mesoderm | 46.29 |
| 12 | Hsp90aa1 | 0.564094872683928 | 1 | 0.999 | 7.4466373648059e-59 | 1.4541793445993e-54 | Paraxial mesoderm | 46.29 |
| 12 | Ppa1 | 0.635197532403613 | 0.975 | 0.817 | 1.411681600541e-55 | 2.75673182953646e-51 | Paraxial mesoderm | 46.29 |
| 12 | Lyar | 0.626376836204106 | 0.975 | 0.803 | 2.68022540376321e-55 | 5.23394416846879e-51 | Paraxial mesoderm | 46.29 |
| 12 | Phgdh | 0.717871423914182 | 0.958 | 0.707 | 3.66604305948012e-53 | 7.15904888655278e-49 | Paraxial mesoderm | 46.29 |
| 12 | Ddx39 | 0.558975423162605 | 0.986 | 0.845 | 7.32966174712329e-52 | 1.43133634597824e-47 | Paraxial mesoderm | 46.29 |
| 12 | Dctpp1 | 0.603439573557315 | 0.989 | 0.948 | 2.09257840856946e-51 | 4.08638711625444e-47 | Paraxial mesoderm | 46.29 |
| 12 | Snhg4 | 0.632046475205438 | 0.827 | 0.495 | 1.39513040412086e-50 | 2.72441065316722e-46 | Paraxial mesoderm | 46.29 |
| 12 | Ncl | 0.589692969860831 | 1 | 0.997 | 1.71316996288527e-43 | 3.34547830352235e-39 | Paraxial mesoderm | 46.29 |
| 12 | Fdps | 0.585452515154707 | 0.989 | 0.933 | 2.05770835568404e-34 | 4.01829287697979e-30 | Paraxial mesoderm | 46.29 |
| 12 | Mest | 0.656253034391376 | 0.887 | 0.835 | 1.69641873414755e-25 | 3.31276650404333e-21 | Paraxial mesoderm | 46.29 |
| 12 | Rbp1 | 0.668315144006746 | 0.926 | 0.743 | 2.70391385905992e-24 | 5.28020298397221e-20 | Paraxial mesoderm | 46.29 |
| 13 | Hist1h2ap | 1.7705692080292 | 0.996 | 0.54 | 1.89795293103502e-134 | 3.70632248372518e-130 | Paraxial mesoderm | 55.4 |
| 13 | Hist1h1b | 1.60209069656647 | 0.996 | 0.477 | 5.10003245589463e-131 | 9.95934337987104e-127 | Paraxial mesoderm | 55.4 |
| 13 | Hist1h2ae | 1.25563729196461 | 0.978 | 0.407 | 3.00423774117674e-129 | 5.86667546096993e-125 | Paraxial mesoderm | 55.4 |
| 13 | Pclaf | 1.39013694761395 | 1 | 0.743 | 2.86454921752475e-116 | 5.59389171198233e-112 | Paraxial mesoderm | 55.4 |
| 13 | Pcna | 1.07971116611186 | 1 | 0.837 | 3.7943980293163e-114 | 7.40970047164887e-110 | Paraxial mesoderm | 55.4 |
| 13 | Dut | 0.93230851248475 | 1 | 0.952 | 1.96295812068152e-106 | 3.83326461806688e-102 | Paraxial mesoderm | 55.4 |
| 13 | Rrm2 | 1.19449352506141 | 0.996 | 0.628 | 1.45299966531011e-105 | 2.83741774641759e-101 | Paraxial mesoderm | 55.4 |
| 13 | Atad2 | 0.898869943407989 | 0.978 | 0.599 | 1.0556760950144e-90 | 2.06152427834412e-86 | Paraxial mesoderm | 55.4 |
| 13 | Tyms | 0.808100136284546 | 0.993 | 0.771 | 1.02234754293932e-88 | 1.9964402818519e-84 | Paraxial mesoderm | 55.4 |
| 13 | Clspn | 0.729423481742938 | 0.935 | 0.464 | 1.25290965778571e-84 | 2.44668197972394e-80 | Paraxial mesoderm | 55.4 |
| 13 | Lig1 | 0.72544354376551 | 0.996 | 0.756 | 3.58376873696874e-81 | 6.99838358955255e-77 | Paraxial mesoderm | 55.4 |
| 13 | Gmnn | 0.761908543410113 | 1 | 0.794 | 7.95109362095499e-79 | 1.55268956230009e-74 | Paraxial mesoderm | 55.4 |
| 13 | Nasp | 0.76242457552783 | 1 | 0.973 | 6.35336918449089e-78 | 1.24068593434738e-73 | Paraxial mesoderm | 55.4 |
| 13 | Dhfr | 0.673300319423618 | 0.982 | 0.659 | 5.01628389888322e-77 | 9.79579919773916e-73 | Paraxial mesoderm | 55.4 |
| 13 | Hist1h1e | 0.783594830895833 | 0.942 | 0.527 | 1.05870055050181e-76 | 2.06743043501994e-72 | Paraxial mesoderm | 55.4 |
| 13 | Uhrf1 | 0.664888855508089 | 0.975 | 0.622 | 6.52007680808858e-70 | 1.27324059908354e-65 | Paraxial mesoderm | 55.4 |
| 13 | Top2a | 0.716049585946181 | 0.993 | 0.571 | 5.90153219123559e-58 | 1.15245120630449e-53 | Paraxial mesoderm | 55.4 |
| 13 | Fabp5 | 0.757826573257157 | 1 | 0.963 | 2.37407766426975e-54 | 4.63609886278597e-50 | Paraxial mesoderm | 55.4 |
| 13 | AC160336. | 0.854215794355215 | 0.996 | 0.858 | 2.03767667332993e-43 | 3.97917500767869e-39 | Paraxial mesoderm | 55.4 |
| 13 | 2610528A1 | 0.688084494639113 | 0.54 | 0.296 | 1.2590883323849e-20 | 2.45874769548123e-16 | Paraxial mesoderm | 55.4 |
| 14 | Hes7 | 1.93942394971884 | 0.902 | 0.05 | 0 | 0 | Somitic mesoderm | 58.17 |
| 14 | Fgf17 | 1.22453661917039 | 0.686 | 0.03 | 0 | 0 | Somitic mesoderm | 58.17 |
| 14 | T | 1.0332579593449 | 0.752 | 0.026 | 0 | 0 | Somitic mesoderm | 58.17 |
| 14 | Sp5 | 0.976604351777598 | 0.732 | 0.036 | 0 | 0 | Somitic mesoderm | 58.17 |
| 14 | Dll3 | 1.13950155067385 | 0.843 | 0.058 | 1.71578054632362e-287 | 3.35057625086077e-283 | Somitic mesoderm | 58.17 |
| 14 | Fgf8 | 1.40679038730499 | 0.771 | 0.063 | 1.12562940395871e-222 | 2.19812910005056e-218 | Somitic mesoderm | 58.17 |
| 14 | Hoxc10 | 1.63342084662695 | 0.869 | 0.119 | 4.32006221460101e-166 | 8.43621749267285e-162 | Somitic mesoderm | 58.17 |
| 14 | Fabp7 | 1.11653145179289 | 0.856 | 0.119 | 9.7152744291776e-160 | 1.8971987905298e-155 | Somitic mesoderm | 58.17 |
| 14 | Rspo3 | 2.35071123105004 | 0.993 | 0.224 | 6.30832488334222e-156 | 1.23188968321907e-151 | Somitic mesoderm | 58.17 |
| 14 | Cdx2 | 1.37271388384523 | 0.869 | 0.135 | 2.63590828318936e-150 | 5.14740169541218e-146 | Somitic mesoderm | 58.17 |
| 14 | Hoxa9 | 0.920221702383961 | 0.941 | 0.197 | 1.47199527619166e-117 | 2.87451237534708e-113 | Somitic mesoderm | 58.17 |
| 14 | Hoxaas3 | 2.21782269185952 | 0.928 | 0.261 | 5.55011596465392e-114 | 1.08382664557762e-109 | Somitic mesoderm | 58.17 |
| 14 | Ifitm1 | 2.40363957899798 | 0.993 | 0.423 | 1.52466431589487e-102 | 2.97736447607959e-98 | Somitic mesoderm | 58.17 |
| 14 | Lef1 | 1.04869642174472 | 0.915 | 0.3 | 1.56477039525037e-86 | 3.05568362784491e-82 | Somitic mesoderm | 58.17 |
| 14 | Hoxb5os | 1.35493069791116 | 0.98 | 0.383 | 3.75554901344031e-81 | 7.33383611344624e-77 | Somitic mesoderm | 58.17 |
| 14 | Tcf15 | 1.01850674576895 | 0.601 | 0.132 | 3.89193711305507e-67 | 7.60017479437393e-63 | Somitic mesoderm | 58.17 |
| 14 | Cited1 | 1.31567880560556 | 0.765 | 0.252 | 5.35780787599939e-61 | 1.04627272202516e-56 | Somitic mesoderm | 58.17 |
| 14 | Gja1 | 0.981687293837032 | 0.935 | 0.474 | 2.45340648593636e-57 | 4.79101218573653e-53 | Somitic mesoderm | 58.17 |
| 14 | Nme1 | 0.887221862894266 | 1 | 0.987 | 1.84635012914006e-54 | 3.60555253218471e-50 | Somitic mesoderm | 58.17 |
| 14 | Ncl | 0.858769691099877 | 1 | 0.997 | 5.73452300797916e-51 | 1.11983765299817e-46 | Somitic mesoderm | 58.17 |
| 15 | Spink1 | 3.20462458879434 | 0.649 | 0.015 | 0 | 0 | Gut | 76.6 |
| 15 | Cldn7 | 1.89215757913012 | 0.968 | 0.016 | 0 | 0 | Gut | 76.6 |
| 15 | Cldn6 | 1.50534329152304 | 0.904 | 0.018 | 0 | 0 | Gut | 76.6 |
| 15 | Epcam | 2.7313601072541 | 0.979 | 0.071 | 1.61502029566185e-239 | 3.15381163336845e-235 | Gut | 76.6 |
| 15 | Pga5 | 1.66507747052838 | 0.202 | 0.002 | 1.11614072234267e-139 | 2.17959960259076e-135 | Gut | 76.6 |
| 15 | Krt19 | 1.56385430835573 | 0.915 | 0.114 | 9.85291521191297e-138 | 1.92407728258236e-133 | Gut | 76.6 |
| 15 | Apela | 1.5963641437734 | 0.872 | 0.132 | 5.3080528595839e-108 | 1.03655656241954e-103 | Gut | 76.6 |
| 15 | Krt18 | 2.96857869002506 | 0.989 | 0.227 | 5.10348944920016e-99 | 9.96609419639807e-95 | Gut | 76.6 |
| 15 | Spint2 | 1.65328108505344 | 0.926 | 0.21 | 2.54607390831059e-92 | 4.97197312814892e-88 | Gut | 76.6 |
| 15 | Car4 | 1.62549974000752 | 0.638 | 0.076 | 1.06624365995141e-89 | 2.08216061915311e-85 | Gut | 76.6 |
| 15 | Espn | 1.39374807290416 | 0.894 | 0.214 | 7.18243503639552e-74 | 1.40258591390732e-69 | Gut | 76.6 |
| 15 | Ttr | 2.74435910926513 | 0.245 | 0.012 | 7.39915096038172e-74 | 1.44490619954334e-69 | Gut | 76.6 |
| 15 | Emb | 2.91151015649097 | 0.979 | 0.354 | 1.36967059122612e-72 | 2.67469273054637e-68 | Gut | 76.6 |
| 15 | Krt8 | 2.00117179351046 | 0.989 | 0.339 | 1.4843072950938e-66 | 2.89855528585917e-62 | Gut | 76.6 |
| 15 | Bex1 | 2.268154405158 | 1 | 0.79 | 6.4885558156243e-56 | 1.26708517967511e-51 | Gut | 76.6 |
| 15 | Igfbp2 | 1.79478442167989 | 0.936 | 0.379 | 1.00756590396218e-53 | 1.96757469725734e-49 | Gut | 76.6 |
| 15 | Pyy | 2.62358667936888 | 0.319 | 0.03 | 2.43652245083174e-53 | 4.75804104198421e-49 | Gut | 76.6 |
| 15 | Slc2a3 | 2.26851779447388 | 0.979 | 0.475 | 1.27502330200191e-52 | 2.48986550414934e-48 | Gut | 76.6 |
| 15 | Bex4 | 1.77075463340151 | 1 | 0.824 | 3.83204247425211e-51 | 7.48321254371952e-47 | Gut | 76.6 |
| 15 | Rbp4 | 3.39708325898451 | 0.596 | 0.133 | 3.13014253384472e-43 | 6.11254234009197e-39 | Gut | 76.6 |
| 16 | Dppa3 | 1.73394701266714 | 0.607 | 0.002 | 0 | 0 | PGC | 64.29 |
| 16 | Utf1 | 2.26051180239239 | 0.679 | 0.006 | 1.03210877532928e-280 | 2.01550201646301e-276 | PGC | 64.29 |
| 16 | Zfp42 | 1.91421410871296 | 0.679 | 0.007 | 1.47818733017831e-256 | 2.88660421837221e-252 | PGC | 64.29 |
| 16 | Tdh | 1.58082402075462 | 0.607 | 0.006 | 3.25697555848274e-237 | 6.36022187060509e-233 | PGC | 64.29 |
| 16 | Dppa5a | 5.42065006064962 | 0.679 | 0.023 | 3.16528105002615e-109 | 6.18116083449106e-105 | PGC | 64.29 |
| 16 | Pou5f1 | 2.22895147823603 | 0.679 | 0.052 | 5.89553736236933e-53 | 1.15128053612348e-48 | PGC | 64.29 |
| 16 | Mt2 | 2.60631219680908 | 0.714 | 0.058 | 7.38612117179026e-53 | 1.4423617424272e-48 | PGC | 64.29 |
| 16 | Hba-a1 | 2.61399183692064 | 0.214 | 0.005 | 6.36917694572599e-48 | 1.24377287396137e-43 | PGC | 64.29 |
| 16 | Bst2 | 1.58967301887685 | 0.75 | 0.107 | 2.4378681042896e-32 | 4.76066883405674e-28 | PGC | 64.29 |
| 16 | Klf2 | 1.94535221744601 | 0.75 | 0.111 | 8.46725171358341e-31 | 1.65348491462857e-26 | PGC | 64.29 |
| 16 | Hbb-bh1 | 2.58430520891166 | 0.25 | 0.012 | 1.09231122073637e-28 | 2.13306535185398e-24 | PGC | 64.29 |
| 16 | Chchd10 | 2.22740317707784 | 0.964 | 0.227 | 2.07573345952315e-27 | 4.0534922997568e-23 | PGC | 64.29 |
| 16 | Hspb1 | 1.59118096638459 | 0.643 | 0.114 | 3.53284354045236e-20 | 6.89893686579536e-16 | PGC | 64.29 |
| 16 | Hbb-bs | 1.74458719474107 | 0.143 | 0.006 | 3.44545585758516e-19 | 6.7282861986923e-15 | PGC | 64.29 |
| 16 | Hba-x | 5.18366116688135 | 0.286 | 0.026 | 1.16571317920018e-17 | 2.27640469634212e-13 | PGC | 64.29 |
| 16 | Slc2a3 | 1.52698106770516 | 0.893 | 0.481 | 2.14179544943128e-11 | 4.18249815364941e-7 | PGC | 64.29 |
| 16 | Mt1 | 3.13827654802704 | 0.893 | 0.534 | 6.92989054575854e-11 | 0.00000135326902577573 | PGC | 64.29 |
| 16 | Hsp90aa1 | 1.76839649874364 | 1 | 0.999 | 2.16452142375121e-10 | 0.00000422687743630135 | PGC | 64.29 |
| 16 | Mkrn1 | 2.12856021616202 | 0.821 | 0.588 | 1.26564523994709e-9 | 0.0000247155202456868 | PGC | 64.29 |
| 16 | Ifitm1 | 2.28291211139583 | 0.821 | 0.437 | 3.4765891443945e-9 | 0.0000678908328117358 | PGC | 64.29 |

**Supplementary Table S1:** **Differential gene expression analysis at day 11**

| Cluster | Gene | avg_logFC | pct.1 | pct.2 | p_val | p_val_adj | most expressed celltype | pct.celltype |
|---|---|---|---|---|---|---|---|---|
| 1 | Angptl1 | 0.868840922543754 | 0.439 | 0.069 | 2.09883764731941e-284 | 4.18571192004911e-280 | Paraxial mesoderm | 94.01 |
| 1 | Prrx1 | 1.20854992885609 | 0.84 | 0.365 | 2.51333973593812e-257 | 5.01235343538139e-253 | Paraxial mesoderm | 94.01 |
| 1 | Gas1 | 1.0987460666599 | 0.944 | 0.611 | 9.9258789442979e-224 | 1.97951803786133e-219 | Paraxial mesoderm | 94.01 |
| 1 | Cdkn1c | 1.27353312423934 | 0.962 | 0.93 | 1.04758154633958e-209 | 2.08919187786502e-205 | Paraxial mesoderm | 94.01 |
| 1 | Col3a1 | 1.23386920529855 | 0.965 | 0.601 | 1.93203341614987e-209 | 3.85305424182769e-205 | Paraxial mesoderm | 94.01 |
| 1 | Lpar1 | 0.886170726952481 | 0.702 | 0.291 | 8.28875442029642e-194 | 1.65302629403971e-189 | Paraxial mesoderm | 94.01 |
| 1 | Mfap4 | 0.879178543580066 | 0.945 | 0.549 | 2.67759376017874e-189 | 5.33992523592446e-185 | Paraxial mesoderm | 94.01 |
| 1 | Col1a2 | 1.06637430795166 | 0.846 | 0.413 | 2.16116481916769e-181 | 4.31001099886613e-177 | Paraxial mesoderm | 94.01 |
| 1 | Igfbp4 | 0.959016302204591 | 0.959 | 0.747 | 7.12412310285697e-179 | 1.42076387040276e-174 | Paraxial mesoderm | 94.01 |
| 1 | Meg3 | 1.03635956294496 | 0.952 | 0.77 | 9.89846074081204e-131 | 1.97405002554014e-126 | Paraxial mesoderm | 94.01 |
| 1 | Ptn | 0.870368013311475 | 0.985 | 0.811 | 1.63872065351041e-128 | 3.26810059929581e-124 | Paraxial mesoderm | 94.01 |
| 1 | Sfrp2 | 1.0229000692035 | 0.616 | 0.297 | 2.77257687941329e-128 | 5.52935007061392e-124 | Paraxial mesoderm | 94.01 |
| 1 | Slc1a3 | 0.870534494563853 | 0.509 | 0.22 | 1.84932351399719e-127 | 3.68810588396459e-123 | Paraxial mesoderm | 94.01 |
| 1 | Ddit4 | 0.895798482621038 | 0.781 | 0.565 | 6.85571615736246e-116 | 1.36723547326279e-111 | Paraxial mesoderm | 94.01 |
| 1 | Mpped2 | 1.07724346614055 | 0.697 | 0.425 | 4.421986768269e-115 | 8.81876821195887e-111 | Paraxial mesoderm | 94.01 |
| 1 | Tgfb2 | 1.00920183570282 | 0.733 | 0.495 | 2.53578833530254e-114 | 5.05712267709385e-110 | Paraxial mesoderm | 94.01 |
| 1 | Igf1 | 1.06738817028719 | 0.629 | 0.324 | 7.12662142250161e-110 | 1.4212621102895e-105 | Paraxial mesoderm | 94.01 |
| 1 | Postn | 0.96989474141339 | 0.553 | 0.26 | 4.45747678938161e-109 | 8.88954596106374e-105 | Paraxial mesoderm | 94.01 |
| 1 | Jun | 0.977516493203888 | 0.838 | 0.646 | 3.51014068877044e-108 | 7.00027357561489e-104 | Paraxial mesoderm | 94.01 |
| 1 | Akap12 | 1.01577873249891 | 0.65 | 0.471 | 6.14028816206171e-67 | 1.22455766815997e-62 | Paraxial mesoderm | 94.01 |
| 2 | Dcn | 1.68668983973419 | 0.624 | 0.181 | 1.46909808502567e-230 | 2.9298223109667e-226 | Paraxial mesoderm | 63.49 |
| 2 | Plac8 | 0.894166409886549 | 0.314 | 0.038 | 7.48788575853337e-218 | 1.49330905682431e-213 | Paraxial mesoderm | 63.49 |
| 2 | Dlk1 | 1.17034925899845 | 0.956 | 0.541 | 5.35278650069644e-198 | 1.06750621183389e-193 | Paraxial mesoderm | 63.49 |
| 2 | Tshz2 | 0.957295147857404 | 0.948 | 0.661 | 1.80869781222e-186 | 3.60708604691035e-182 | Paraxial mesoderm | 63.49 |
| 2 | Cald1 | 0.921361998434676 | 0.986 | 0.828 | 4.12195583525255e-186 | 8.22041652224416e-182 | Paraxial mesoderm | 63.49 |
| 2 | Ifitm3 | 1.41976684805592 | 0.832 | 0.438 | 2.93183901124818e-184 | 5.84696654013225e-180 | Paraxial mesoderm | 63.49 |
| 2 | Zeb2 | 1.1212628401979 | 0.89 | 0.565 | 5.73868454845689e-178 | 1.14446585949876e-173 | Paraxial mesoderm | 63.49 |
| 2 | Sfrp1 | 0.983756585399303 | 0.943 | 0.634 | 1.54012048933395e-165 | 3.0714622918787e-161 | Paraxial mesoderm | 63.49 |
| 2 | Ptn | 1.01670630247269 | 0.999 | 0.812 | 5.25376920284872e-163 | 1.04775919212412e-158 | Paraxial mesoderm | 63.49 |
| 2 | Nr2f2 | 0.946202072262379 | 0.897 | 0.606 | 5.55955664891188e-161 | 1.1087423824925e-156 | Paraxial mesoderm | 63.49 |
| 2 | Emp3 | 0.898229278537707 | 0.895 | 0.546 | 2.59539911149106e-159 | 5.17600444804662e-155 | Paraxial mesoderm | 63.49 |
| 2 | Mfap2 | 0.819034828034483 | 0.941 | 0.666 | 1.26059282081703e-143 | 2.5140002625554e-139 | Paraxial mesoderm | 63.49 |
| 2 | Col1a1 | 0.917629016695656 | 0.824 | 0.428 | 5.17404036588718e-137 | 1.03185887016888e-132 | Paraxial mesoderm | 63.49 |
| 2 | Itm2a | 0.967016015927239 | 0.959 | 0.741 | 5.34633560988632e-135 | 1.06621971067963e-130 | Paraxial mesoderm | 63.49 |
| 2 | Sparcl1 | 0.934361674887383 | 0.83 | 0.507 | 2.69677474359459e-120 | 5.3781778711507e-116 | Paraxial mesoderm | 63.49 |
| 2 | Peg3 | 0.911548766351292 | 0.959 | 0.772 | 1.35332390206065e-117 | 2.69893385787955e-113 | Paraxial mesoderm | 63.49 |
| 2 | Lum | 0.92425295486981 | 0.453 | 0.153 | 3.23922791431562e-116 | 6.45999222951964e-112 | Paraxial mesoderm | 63.49 |
| 2 | Col1a2 | 0.832336498729091 | 0.761 | 0.432 | 9.45532841032053e-92 | 1.88567614487022e-87 | Paraxial mesoderm | 63.49 |
| 2 | Ecm1 | 1.08146025677571 | 0.448 | 0.209 | 6.92667670333456e-81 | 1.38138713494601e-76 | Paraxial mesoderm | 63.49 |
| 2 | Mgp | 0.852465745260807 | 0.414 | 0.248 | 4.81373460796246e-36 | 9.60003092865954e-32 | Paraxial mesoderm | 63.49 |
| 3 | 2410006H1 | 1.480606878169 | 0.998 | 0.891 | 2.22832507285285e-227 | 4.44394869279044e-223 | Paraxial mesoderm | 75.99 |
| 3 | Snhg1 | 1.23402581735086 | 0.974 | 0.831 | 4.64628633812762e-178 | 9.26608884412791e-174 | Paraxial mesoderm | 75.99 |
| 3 | Tac2 | 1.36550061486168 | 0.667 | 0.208 | 4.1914432346353e-172 | 8.35899524283318e-168 | Paraxial mesoderm | 75.99 |
| 3 | Epb41l4ao | 1.13447953209124 | 0.895 | 0.579 | 2.53195697065241e-170 | 5.04948178657211e-166 | Paraxial mesoderm | 75.99 |
| 3 | Gas5 | 0.939388389805414 | 0.995 | 0.978 | 7.90338841094351e-165 | 1.57617275079447e-160 | Paraxial mesoderm | 75.99 |
| 3 | Ndufa4l2 | 1.29430212000131 | 0.78 | 0.347 | 6.33338121000245e-150 | 1.26306621471079e-145 | Paraxial mesoderm | 75.99 |
| 3 | Rcn3 | 0.91889445615591 | 0.906 | 0.635 | 3.38136445719572e-133 | 6.74345513698542e-129 | Paraxial mesoderm | 75.99 |
| 3 | Itm2a | 1.35389641271999 | 0.964 | 0.749 | 4.62297718483448e-131 | 9.2196033997154e-127 | Paraxial mesoderm | 75.99 |
| 3 | Tmem45a | 0.874360425894646 | 0.713 | 0.305 | 1.70114544128422e-130 | 3.39259435355311e-126 | Paraxial mesoderm | 75.99 |
| 3 | P4ha2 | 0.978979208649021 | 0.857 | 0.534 | 4.02746455572346e-121 | 8.0319725634793e-117 | Paraxial mesoderm | 75.99 |
| 3 | P4ha1 | 0.931810142815925 | 0.927 | 0.749 | 8.12762030173921e-118 | 1.62089131677585e-113 | Paraxial mesoderm | 75.99 |
| 3 | Pgk1 | 0.853093031191774 | 0.998 | 0.989 | 9.32569910920857e-109 | 1.85982417334946e-104 | Paraxial mesoderm | 75.99 |
| 3 | Pmp22 | 0.880598232447151 | 0.723 | 0.384 | 2.22480617178938e-102 | 4.43693094839957e-98 | Paraxial mesoderm | 75.99 |
| 3 | 1110038B1 | 0.815467327370877 | 0.906 | 0.77 | 1.47035213023616e-98 | 2.93232325332996e-94 | Paraxial mesoderm | 75.99 |
| 3 | P4hb | 0.815469390348095 | 0.889 | 0.743 | 9.43032189655143e-94 | 1.88068909582925e-89 | Paraxial mesoderm | 75.99 |
| 3 | Meg3 | 0.894590255250779 | 0.965 | 0.779 | 1.58262526393644e-88 | 3.15622956386843e-84 | Paraxial mesoderm | 75.99 |
| 3 | Ero1l | 0.924701551203617 | 0.815 | 0.544 | 1.05231581594638e-86 | 2.09863343174187e-82 | Paraxial mesoderm | 75.99 |
| 3 | Peg3 | 1.05888509592382 | 0.941 | 0.781 | 3.1723822648522e-85 | 6.32668195079474e-81 | Paraxial mesoderm | 75.99 |
| 3 | Cxcl12 | 0.90036848872106 | 0.6 | 0.281 | 3.06134059487218e-84 | 6.10523154835359e-80 | Paraxial mesoderm | 75.99 |
| 3 | Col1a1 | 0.839238066420378 | 0.787 | 0.447 | 1.94171722753801e-81 | 3.87236666687905e-77 | Paraxial mesoderm | 75.99 |
| 4 | Top2a | 2.53928063615229 | 0.971 | 0.214 | 0 | 0 | Paraxial mesoderm | 82.68 |
| 4 | Pclaf | 2.20528774639707 | 0.988 | 0.269 | 0 | 0 | Paraxial mesoderm | 82.68 |
| 4 | Birc5 | 2.08360332340092 | 0.985 | 0.239 | 0 | 0 | Paraxial mesoderm | 82.68 |
| 4 | Cenpf | 2.02482430508502 | 0.911 | 0.181 | 0 | 0 | Paraxial mesoderm | 82.68 |
| 4 | Mki67 | 1.910845498132 | 0.914 | 0.166 | 0 | 0 | Paraxial mesoderm | 82.68 |
| 4 | Nusap1 | 1.90865606395223 | 0.83 | 0.121 | 0 | 0 | Paraxial mesoderm | 82.68 |
| 4 | Tpx2 | 1.79383726916349 | 0.923 | 0.21 | 0 | 0 | Paraxial mesoderm | 82.68 |
| 4 | Prc1 | 1.77017301264586 | 0.844 | 0.133 | 0 | 0 | Paraxial mesoderm | 82.68 |
| 4 | Cdca8 | 1.55538778477236 | 0.955 | 0.244 | 0 | 0 | Paraxial mesoderm | 82.68 |
| 4 | Cenpe | 1.52833174949065 | 0.818 | 0.155 | 0 | 0 | Paraxial mesoderm | 82.68 |
| 4 | Smc2 | 1.56568447049436 | 0.973 | 0.41 | 2.70140500161557e-289 | 5.38741199472194e-285 | Paraxial mesoderm | 82.68 |
| 4 | Ube2c | 1.96544761213097 | 0.887 | 0.257 | 9.66354757596457e-283 | 1.92720129307461e-278 | Paraxial mesoderm | 82.68 |
| 4 | Hist1h1b | 2.48408065321251 | 0.899 | 0.318 | 1.12341777690119e-264 | 2.24043207247405e-260 | Paraxial mesoderm | 82.68 |
| 4 | Hmgb2 | 1.95470329252995 | 1 | 0.76 | 9.28769807435493e-262 | 1.8522456269686e-257 | Paraxial mesoderm | 82.68 |
| 4 | H2afx | 1.98502936422569 | 0.979 | 0.598 | 1.09791484694821e-251 | 2.18957157926881e-247 | Paraxial mesoderm | 82.68 |
| 4 | Cks2 | 1.70205320560899 | 0.95 | 0.457 | 1.28975328489182e-236 | 2.57215497605976e-232 | Paraxial mesoderm | 82.68 |
| 4 | Cenpa | 1.56237931364099 | 0.763 | 0.265 | 1.98448749347563e-188 | 3.95766340823845e-184 | Paraxial mesoderm | 82.68 |
| 4 | Hist1h1e | 1.68562038166237 | 0.923 | 0.678 | 2.49174129991787e-154 | 4.96927967442622e-150 | Paraxial mesoderm | 82.68 |
| 4 | Tubb4b | 1.44235618386963 | 0.943 | 0.639 | 2.80775844182355e-134 | 5.59951266052871e-130 | Paraxial mesoderm | 82.68 |
| 4 | Arl6ip1 | 1.46476808067103 | 0.93 | 0.741 | 3.47591701654878e-100 | 6.93202130610323e-96 | Paraxial mesoderm | 82.68 |
| 5 | Pde1a | 1.06925930501028 | 0.727 | 0.153 | 1.68007777006978e-255 | 3.35057909685016e-251 | Mesenchyme | 79.37 |
| 5 | Gucy1a1 | 1.60378011606123 | 0.903 | 0.354 | 1.87369248633171e-210 | 3.73670492549132e-206 | Mesenchyme | 79.37 |
| 5 | Fn1 | 1.4852656783642 | 0.928 | 0.518 | 1.02093685703418e-179 | 2.03605437398327e-175 | Mesenchyme | 79.37 |
| 5 | Gata6 | 1.14216159151874 | 0.95 | 0.496 | 1.08542753765643e-162 | 2.16466813834821e-158 | Mesenchyme | 79.37 |
| 5 | Ptn | 1.4477125341067 | 1 | 0.822 | 3.52724041096787e-157 | 7.03437555159323e-153 | Mesenchyme | 79.37 |
| 5 | Tagln2 | 1.18405214058288 | 0.818 | 0.316 | 1.33309256616974e-156 | 2.65858650471231e-152 | Mesenchyme | 79.37 |
| 5 | Maged2 | 1.34639520628116 | 0.991 | 0.864 | 9.02860297260706e-156 | 1.80057429082703e-151 | Mesenchyme | 79.37 |
| 5 | Rbp1 | 1.56474476514966 | 0.959 | 0.674 | 1.77871178858328e-155 | 3.54728491997164e-151 | Mesenchyme | 79.37 |
| 5 | Apoe | 1.54982431329767 | 0.885 | 0.48 | 8.70198070132366e-147 | 1.73543601126498e-142 | Mesenchyme | 79.37 |
| 5 | Col3a1 | 1.22359103577646 | 0.974 | 0.625 | 2.51682179158946e-132 | 5.01929769896686e-128 | Mesenchyme | 79.37 |
| 5 | Cst3 | 1.09925915377857 | 0.994 | 0.954 | 1.56216567492182e-121 | 3.11542700549659e-117 | Mesenchyme | 79.37 |
| 5 | Gal | 1.21432909613314 | 0.27 | 0.039 | 6.28818449654333e-119 | 1.25405263414564e-114 | Mesenchyme | 79.37 |
| 5 | Gucy1b1 | 0.987858570638018 | 0.76 | 0.337 | 2.4018286633449e-118 | 4.78996690330873e-114 | Mesenchyme | 79.37 |
| 5 | Nrk | 1.07859223526286 | 0.827 | 0.422 | 3.38634067877982e-118 | 6.7533792156906e-114 | Mesenchyme | 79.37 |
| 5 | Sparc | 1.02824122288918 | 0.991 | 0.838 | 6.52922901857379e-117 | 1.30212414317417e-112 | Mesenchyme | 79.37 |
| 5 | Fbln1 | 1.17373843803892 | 0.914 | 0.643 | 1.42264759602029e-102 | 2.83718610074326e-98 | Mesenchyme | 79.37 |
| 5 | Socs2 | 0.98553081730345 | 0.602 | 0.266 | 9.4632054392137e-87 | 1.88724706074239e-82 | Mesenchyme | 79.37 |
| 5 | Map1b | 1.04075582403465 | 0.914 | 0.694 | 6.94556094586352e-80 | 1.38515321943356e-75 | Mesenchyme | 79.37 |
| 5 | Ndufa4l2 | 1.13683288490156 | 0.703 | 0.359 | 7.95641700268196e-74 | 1.58674824284486e-69 | Mesenchyme | 79.37 |
| 5 | Mfap4 | 1.08538991234825 | 0.745 | 0.591 | 3.19957001525572e-40 | 6.38090248142449e-36 | Mesenchyme | 79.37 |
| 6 | Acta2 | 4.0064354608501 | 0.942 | 0.296 | 0 | 0 | Cardiomyocytes | 99.8 |
| 6 | Csrp3 | 2.76401408505678 | 0.968 | 0.182 | 0 | 0 | Cardiomyocytes | 99.8 |
| 6 | Tnnc1 | 2.4359222061784 | 0.98 | 0.268 | 0 | 0 | Cardiomyocytes | 99.8 |
| 6 | Pgam2 | 2.38522694714772 | 0.968 | 0.175 | 0 | 0 | Cardiomyocytes | 99.8 |
| 6 | Cryab | 2.28912301439138 | 0.97 | 0.188 | 0 | 0 | Cardiomyocytes | 99.8 |
| 6 | Shox2 | 2.20010508128991 | 0.844 | 0.15 | 0 | 0 | Cardiomyocytes | 99.8 |
| 6 | Fbxo32 | 2.02600961704982 | 0.882 | 0.166 | 0 | 0 | Cardiomyocytes | 99.8 |
| 6 | Vsnl1 | 1.99360970025011 | 0.812 | 0.077 | 0 | 0 | Cardiomyocytes | 99.8 |
| 6 | Atp1b1 | 1.99278179384135 | 0.958 | 0.238 | 0 | 0 | Cardiomyocytes | 99.8 |
| 6 | Ppp1r14c | 2.11019608181192 | 0.924 | 0.23 | 2.7675423559584e-306 | 5.51930972048784e-302 | Cardiomyocytes | 99.8 |
| 6 | Tnnt2 | 2.52367315651155 | 0.994 | 0.31 | 1.37671680871766e-301 | 2.74558633162563e-297 | Cardiomyocytes | 99.8 |
| 6 | Tnni1 | 2.17969690028688 | 0.998 | 0.287 | 2.45373037351129e-297 | 4.89347448389357e-293 | Cardiomyocytes | 99.8 |
| 6 | Chchd10 | 2.16380078602983 | 0.984 | 0.293 | 5.88444041513933e-293 | 1.17353395199124e-288 | Cardiomyocytes | 99.8 |
| 6 | Myl4 | 2.6736897298911 | 0.994 | 0.346 | 9.89799677010487e-291 | 1.97395749586201e-286 | Cardiomyocytes | 99.8 |
| 6 | Cox6a2 | 2.08211652396122 | 0.922 | 0.211 | 1.36448365137464e-289 | 2.72118974593645e-285 | Cardiomyocytes | 99.8 |
| 6 | Myl7 | 3.01206971156742 | 1 | 0.431 | 2.06777358538218e-283 | 4.12376086132769e-279 | Cardiomyocytes | 99.8 |
| 6 | Myl9 | 2.81707108572271 | 0.93 | 0.299 | 6.07546045055339e-270 | 1.21162907765386e-265 | Cardiomyocytes | 99.8 |
| 6 | Actc1 | 2.19855648745787 | 0.972 | 0.389 | 9.71858343706998e-226 | 1.93817709485487e-221 | Cardiomyocytes | 99.8 |
| 6 | Tpm1 | 2.44794748357736 | 1 | 0.904 | 1.0177182165562e-224 | 2.02963543927803e-220 | Cardiomyocytes | 99.8 |
| 6 | Gyg | 1.96709913977738 | 0.974 | 0.475 | 2.16346847513522e-221 | 4.31460517996217e-217 | Cardiomyocytes | 99.8 |
| 7 | Ctla2a | 3.79291942072411 | 0.965 | 0.032 | 0 | 0 | Endothelium | 96.49 |
| 7 | Ramp2 | 2.8499592555637 | 0.961 | 0.29 | 0 | 0 | Endothelium | 96.49 |
| 7 | Icam2 | 2.64619511161468 | 0.956 | 0.019 | 0 | 0 | Endothelium | 96.49 |
| 7 | Tagln2 | 2.63481413707248 | 0.996 | 0.311 | 0 | 0 | Endothelium | 96.49 |
| 7 | Egfl7 | 2.59878802530858 | 0.987 | 0.368 | 0 | 0 | Endothelium | 96.49 |
| 7 | Ecscr | 2.54113139960765 | 0.947 | 0.029 | 0 | 0 | Endothelium | 96.49 |
| 7 | Emcn | 2.3077328153644 | 0.934 | 0.026 | 0 | 0 | Endothelium | 96.49 |
| 7 | Igfbp3 | 2.13924052414924 | 0.759 | 0.087 | 0 | 0 | Endothelium | 96.49 |
| 7 | Pecam1 | 2.04724589715099 | 0.89 | 0.02 | 0 | 0 | Endothelium | 96.49 |
| 7 | Flt1 | 2.0347999445922 | 0.857 | 0.019 | 0 | 0 | Endothelium | 96.49 |
| 7 | Rasip1 | 2.03200726223372 | 0.904 | 0.036 | 0 | 0 | Endothelium | 96.49 |
| 7 | Ednrb | 1.87596687479759 | 0.895 | 0.152 | 0 | 0 | Endothelium | 96.49 |
| 7 | Kcne3 | 1.84785327241199 | 0.673 | 0.009 | 0 | 0 | Endothelium | 96.49 |
| 7 | Kdr | 1.84361444840211 | 0.844 | 0.051 | 0 | 0 | Endothelium | 96.49 |
| 7 | Cdh5 | 1.8207457354488 | 0.871 | 0.009 | 0 | 0 | Endothelium | 96.49 |
| 7 | Fabp4 | 2.43480516250359 | 0.371 | 0.022 | 1.4870957342904e-295 | 2.96571502289534e-291 | Endothelium | 96.49 |

**Supplementary Table S1:** **Differential gene expression analysis at day 11**

| Cluster | Gene | avg_logFC | pct.1 | pct.2 | p_val | p_val_adj | most expressed celltype | pct.celltype |
|---|---|---|---|---|---|---|---|---|
| 7 | Gng11 | 2.25513704675031 | 0.969 | 0.473 | 1.06684298091973e-251 | 2.12760495684821e-247 | Endothelium | 96.49 |
| 7 | Col4a1 | 2.10981289357377 | 0.897 | 0.425 | 2.01967349475085e-184 | 4.02783485058161e-180 | Endothelium | 96.49 |
| 7 | Fabp5 | 2.18159393720349 | 0.996 | 0.923 | 5.5216360356567e-184 | 1.10117987459102e-179 | Endothelium | 96.49 |
| 7 | Odc1 | 2.34271169822384 | 0.884 | 0.556 | 7.44784028226209e-145 | 1.48532278749153e-140 | Endothelium | 96.49 |
| 8 | Myl2 | 5.27315213778398 | 0.95 | 0.129 | 0 | 0 | Cardiomyocytes | 98.65 |
| 8 | Myl3 | 4.88416409960998 | 0.984 | 0.169 | 0 | 0 | Cardiomyocytes | 98.65 |
| 8 | Myh7 | 4.35348748816714 | 0.982 | 0.125 | 0 | 0 | Cardiomyocytes | 98.65 |
| 8 | Pln | 3.27383275563014 | 0.932 | 0.08 | 0 | 0 | Cardiomyocytes | 98.65 |
| 8 | Tnni3 | 2.90605245566437 | 0.989 | 0.189 | 0 | 0 | Cardiomyocytes | 98.65 |
| 8 | Ttn | 2.75913874086919 | 0.986 | 0.218 | 0 | 0 | Cardiomyocytes | 98.65 |
| 8 | Cox6a2 | 2.72600190210877 | 0.995 | 0.212 | 0 | 0 | Cardiomyocytes | 98.65 |
| 8 | Tnni1 | 2.71668530234238 | 1 | 0.292 | 0 | 0 | Cardiomyocytes | 98.65 |
| 8 | Nexn | 2.2452349236693 | 0.98 | 0.229 | 0 | 0 | Cardiomyocytes | 98.65 |
| 8 | Atcayos | 2.15985510521637 | 0.932 | 0.126 | 0 | 0 | Cardiomyocytes | 98.65 |
| 8 | Sh3bgr | 2.13203126005548 | 0.973 | 0.166 | 0 | 0 | Cardiomyocytes | 98.65 |
| 8 | Nebl | 2.03514208174966 | 0.95 | 0.125 | 0 | 0 | Cardiomyocytes | 98.65 |
| 8 | Chchd10 | 2.43598935653931 | 0.998 | 0.297 | 1.35668679978774e-300 | 2.7056404848167e-296 | Cardiomyocytes | 98.65 |
| 8 | Crip2 | 2.59071479406776 | 0.991 | 0.36 | 3.73018007478271e-299 | 7.43909812313916e-295 | Cardiomyocytes | 98.65 |
| 8 | Tnnt2 | 2.72951263690885 | 1 | 0.314 | 8.07539907089307e-285 | 1.6104768367082e-280 | Cardiomyocytes | 98.65 |
| 8 | Tnnc1 | 2.14947998147377 | 0.995 | 0.273 | 1.18554590503304e-280 | 2.36433419840739e-276 | Cardiomyocytes | 98.65 |
| 8 | Gyg | 2.58265555476293 | 0.998 | 0.477 | 5.31779474533859e-263 | 1.06052780606287e-258 | Cardiomyocytes | 98.65 |
| 8 | Actc1 | 2.8544411912125 | 1 | 0.391 | 4.47005266624738e-261 | 8.91462603229714e-257 | Cardiomyocytes | 98.65 |
| 8 | Myl4 | 2.19266489557739 | 1 | 0.35 | 5.29487608031964e-242 | 1.05595713669815e-237 | Cardiomyocytes | 98.65 |
| 8 | Tpm1 | 2.10229708320551 | 1 | 0.904 | 6.86205884772662e-193 | 1.36850039600212e-188 | Cardiomyocytes | 98.65 |
| 9 | Myh6 | 3.95177515924212 | 1 | 0.168 | 0 | 0 | Cardiomyocytes | 100 |
| 9 | Myl4 | 3.51516610835834 | 1 | 0.351 | 0 | 0 | Cardiomyocytes | 100 |
| 9 | Myl9 | 3.30563541877203 | 0.993 | 0.302 | 0 | 0 | Cardiomyocytes | 100 |
| 9 | Ttn | 3.21820476432588 | 0.995 | 0.219 | 0 | 0 | Cardiomyocytes | 100 |
| 9 | Tnni3 | 3.17123600725553 | 0.988 | 0.191 | 0 | 0 | Cardiomyocytes | 100 |
| 9 | Cox6a2 | 3.14707882649476 | 1 | 0.214 | 0 | 0 | Cardiomyocytes | 100 |
| 9 | Csrp3 | 3.12387371767418 | 1 | 0.187 | 0 | 0 | Cardiomyocytes | 100 |
| 9 | Tnnc1 | 3.08328329708506 | 1 | 0.274 | 0 | 0 | Cardiomyocytes | 100 |
| 9 | Pgam2 | 2.80915192418944 | 0.998 | 0.181 | 0 | 0 | Cardiomyocytes | 100 |
| 9 | Cryab | 2.80413419630777 | 1 | 0.193 | 0 | 0 | Cardiomyocytes | 100 |
| 9 | Casq1 | 2.65302788590929 | 0.883 | 0.076 | 0 | 0 | Cardiomyocytes | 100 |
| 9 | Gpx3 | 2.64240187405824 | 0.993 | 0.273 | 0 | 0 | Cardiomyocytes | 100 |
| 9 | Hspb1 | 2.5704505685449 | 0.981 | 0.209 | 0 | 0 | Cardiomyocytes | 100 |
| 9 | Chchd10 | 2.56294136567125 | 0.998 | 0.299 | 7.11321331176845e-307 | 1.41858813076598e-302 | Cardiomyocytes | 100 |
| 9 | Tnni1 | 2.69292342372071 | 1 | 0.294 | 5.69332577619545e-303 | 1.13541995954666e-298 | Cardiomyocytes | 100 |
| 9 | Tnnt2 | 3.09098700479463 | 1 | 0.316 | 1.20417814398328e-298 | 2.40149247254586e-294 | Cardiomyocytes | 100 |
| 9 | Myl7 | 3.70706925665188 | 1 | 0.436 | 4.21439164837322e-282 | 8.40476126435071e-278 | Cardiomyocytes | 100 |
| 9 | Actc1 | 3.18188229621596 | 1 | 0.393 | 5.85544745734723e-273 | 1.16775188641876e-268 | Cardiomyocytes | 100 |
| 9 | Atp2a2 | 2.57573981086323 | 0.998 | 0.572 | 1.90334616231536e-244 | 3.79584325150552e-240 | Cardiomyocytes | 100 |
| 9 | Tpm1 | 2.80820580094081 | 1 | 0.905 | 3.56943330174634e-225 | 7.11852083367274e-221 | Cardiomyocytes | 100 |
| 0 | Tubb3 | 3.8449864047631 | 0.971 | 0.098 | 0 | 0 | Forebrain/Midbrain/Hindbra | 85.97 |
| 10 | Igfbpl1 | 3.14924315057788 | 0.894 | 0.146 | 0 | 0 | Forebrain/Midbrain/Hindbra | 85.97 |
| 10 | Map2 | 2.59798273848019 | 0.894 | 0.109 | 0 | 0 | Forebrain/Midbrain/Hindbra | 85.97 |
| 10 | Nhlh2 | 2.42654869698953 | 0.712 | 0.031 | 0 | 0 | Forebrain/Midbrain/Hindbra | 85.97 |
| 10 | Tagln3 | 2.23267192302369 | 0.935 | 0.142 | 0 | 0 | Forebrain/Midbrain/Hindbra | 85.97 |
| 10 | Hoxb9 | 2.21286792368711 | 0.922 | 0.14 | 0 | 0 | Forebrain/Midbrain/Hindbra | 85.97 |
| 10 | Hoxa9 | 2.13982755875104 | 0.886 | 0.173 | 0 | 0 | Forebrain/Midbrain/Hindbra | 85.97 |
| 10 | Srrm4 | 2.12940047789598 | 0.829 | 0.039 | 0 | 0 | Forebrain/Midbrain/Hindbra | 85.97 |
| 10 | Dcc | 2.12140798939274 | 0.797 | 0.063 | 0 | 0 | Forebrain/Midbrain/Hindbra | 85.97 |
| 10 | Elavl3 | 1.98050004442133 | 0.878 | 0.06 | 0 | 0 | Forebrain/Midbrain/Hindbra | 85.97 |
| 10 | Pou3f2 | 1.91469418490475 | 0.808 | 0.079 | 0 | 0 | Forebrain/Midbrain/Hindbra | 85.97 |
| 10 | Rnd2 | 2.16029955069744 | 0.803 | 0.158 | 1.88133088574372e-293 | 3.7519381854387e-289 | Forebrain/Midbrain/Hindbra | 85.97 |
| 10 | Rtn1 | 1.91434383901926 | 0.738 | 0.111 | 1.9983167631351e-279 | 3.98524312072034e-275 | Forebrain/Midbrain/Hindbra | 85.97 |
| 10 | Sox11 | 2.78214843919004 | 1 | 0.765 | 2.86675070763147e-223 | 5.71716093622944e-219 | Forebrain/Midbrain/Hindbra | 85.97 |
| 10 | Tubb2b | 2.6835000916225 | 1 | 0.645 | 2.83302434806752e-213 | 5.64990045735107e-209 | Forebrain/Midbrain/Hindbra | 85.97 |
| 10 | Tuba1a | 2.56460855862518 | 1 | 0.934 | 7.52384641535899e-209 | 1.50048069061504e-204 | Forebrain/Midbrain/Hindbra | 85.97 |
| 10 | Ebf1 | 2.27497788421435 | 0.87 | 0.298 | 4.97527345329308e-200 | 9.92218784790239e-196 | Forebrain/Midbrain/Hindbra | 85.97 |
| 10 | Ckb | 2.33214388965673 | 0.995 | 0.601 | 2.09200667114402e-175 | 4.17208890426252e-171 | Forebrain/Midbrain/Hindbra | 85.97 |
| 10 | Gadd45g | 2.18826941630538 | 0.668 | 0.265 | 6.52696165047333e-109 | 1.3016719619539e-104 | Forebrain/Midbrain/Hindbra | 85.97 |
| 10 | Crabp1 | 3.35845766216826 | 0.738 | 0.345 | 6.71865521992407e-100 | 1.33990141050946e-95 | Forebrain/Midbrain/Hindbra | 85.97 |
| 11 | Cxcl14 | 1.20579860229466 | 0.494 | 0.037 | 4.50625307524839e-288 | 8.98682050796786e-284 | Paraxial mesoderm | 98.82 |
| 11 | 2700069I18 | 1.06246146594138 | 0.618 | 0.076 | 2.30432020345764e-251 | 4.59550578175557e-247 | Paraxial mesoderm | 98.82 |
| 11 | Hoxc10 | 1.62862953186999 | 0.747 | 0.167 | 9.68965179617202e-186 | 1.93240725771059e-181 | Paraxial mesoderm | 98.82 |
| 11 | Irx3 | 1.47898481963825 | 0.715 | 0.167 | 4.63506064930798e-176 | 9.2437014529149e-172 | Paraxial mesoderm | 98.82 |
| 11 | Lix1 | 1.31326695047174 | 0.668 | 0.139 | 5.36260110346577e-172 | 1.06946353806418e-167 | Paraxial mesoderm | 98.82 |
| 11 | Prrx1 | 1.84560962197928 | 0.965 | 0.403 | 1.82485105181004e-170 | 3.63930045262477e-166 | Paraxial mesoderm | 98.82 |
| 11 | Hoxc9 | 1.1873474890048 | 0.8 | 0.232 | 1.34078028670088e-138 | 2.67391812576757e-134 | Paraxial mesoderm | 98.82 |
| 11 | Zfhx4 | 1.78997686225072 | 0.921 | 0.487 | 5.42825810928707e-138 | 1.08255751473512e-133 | Paraxial mesoderm | 98.82 |
| 11 | Crabp2 | 1.1986190034208 | 0.897 | 0.312 | 1.64653139545688e-134 | 3.28367756195966e-130 | Paraxial mesoderm | 98.82 |
| 11 | Zfhx3 | 1.80457083065236 | 0.929 | 0.537 | 1.72289025867591e-129 | 3.43596004287736e-125 | Paraxial mesoderm | 98.82 |
| 11 | Plpp3 | 1.23339155399855 | 0.85 | 0.39 | 2.00523919145184e-112 | 3.9990485195124e-108 | Paraxial mesoderm | 98.82 |
| 11 | Twist1 | 1.37496512638867 | 0.903 | 0.495 | 7.34186410954663e-105 | 1.46418795936688e-100 | Paraxial mesoderm | 98.82 |
| 11 | Lpar1 | 1.04572031235262 | 0.803 | 0.325 | 4.33641865559442e-103 | 8.64811972485196e-99 | Paraxial mesoderm | 98.82 |
| 11 | Cdo1 | 1.1396845705967 | 0.838 | 0.371 | 2.82641398694598e-101 | 5.63671741416636e-97 | Paraxial mesoderm | 98.82 |
| 11 | Sema3a | 1.06969860559831 | 0.629 | 0.206 | 1.79708393026709e-99 | 3.58392448213165e-95 | Paraxial mesoderm | 98.82 |
| 11 | Hmga2 | 1.10725736271724 | 0.953 | 0.599 | 4.23650148185819e-86 | 8.44885490526979e-82 | Paraxial mesoderm | 98.82 |
| 11 | Tcf4 | 1.28981416306067 | 0.941 | 0.783 | 4.25988576573103e-82 | 8.49549018259739e-78 | Paraxial mesoderm | 98.82 |
| 11 | Mest | 1.23739722994387 | 0.926 | 0.666 | 2.21275564221314e-77 | 4.41289857726567e-73 | Paraxial mesoderm | 98.82 |
| 11 | Gas1 | 1.03691410466887 | 0.944 | 0.642 | 3.24876010260794e-76 | 6.47900227263102e-72 | Paraxial mesoderm | 98.82 |
| 11 | Igf1 | 1.19713574208978 | 0.741 | 0.347 | 5.02472035704601e-68 | 1.00207998080569e-63 | Paraxial mesoderm | 98.82 |
| 12 | Stmn2 | 4.17946102663931 | 0.956 | 0.101 | 0 | 0 | Forebrain/Midbrain/Hindbra | 90.88 |
| 12 | Rtn1 | 4.05812636540372 | 0.98 | 0.109 | 0 | 0 | Forebrain/Midbrain/Hindbra | 90.88 |
| 12 | Cartpt | 3.84229752444086 | 0.429 | 0.014 | 0 | 0 | Forebrain/Midbrain/Hindbra | 90.88 |
| 12 | Ina | 3.68235809106217 | 0.949 | 0.042 | 0 | 0 | Forebrain/Midbrain/Hindbra | 90.88 |
| 12 | Stmn3 | 3.46139925517146 | 0.98 | 0.061 | 0 | 0 | Forebrain/Midbrain/Hindbra | 90.88 |
| 12 | Tubb3 | 3.02721446983429 | 0.997 | 0.108 | 0 | 0 | Forebrain/Midbrain/Hindbra | 90.88 |
| 12 | Pcsk1n | 2.79334897648644 | 0.875 | 0.056 | 0 | 0 | Forebrain/Midbrain/Hindbra | 90.88 |
| 12 | Kif5c | 2.75799555823605 | 0.963 | 0.133 | 0 | 0 | Forebrain/Midbrain/Hindbra | 90.88 |
| 12 | Gng3 | 2.67070243088882 | 0.851 | 0.037 | 0 | 0 | Forebrain/Midbrain/Hindbra | 90.88 |
| 12 | Crmp1 | 2.54153876876738 | 0.953 | 0.079 | 0 | 0 | Forebrain/Midbrain/Hindbra | 90.88 |
| 12 | Mapt | 2.50025294783188 | 0.895 | 0.055 | 0 | 0 | Forebrain/Midbrain/Hindbra | 90.88 |
| 12 | Igfbpl1 | 2.59421897704572 | 0.895 | 0.155 | 4.86923483009211e-269 | 9.71071502165269e-265 | Forebrain/Midbrain/Hindbra | 90.88 |
| 12 | Mllt11 | 2.71939975929202 | 0.986 | 0.322 | 1.21155721441269e-236 | 2.41620855270323e-232 | Forebrain/Midbrain/Hindbra | 90.88 |
| 12 | Hist3h2ba | 2.54794738712998 | 0.976 | 0.31 | 2.70037334145428e-219 | 5.38535455486227e-215 | Forebrain/Midbrain/Hindbra | 90.88 |
| 12 | Gap43 | 2.7367869196805 | 0.75 | 0.132 | 1.66015610652228e-216 | 3.31084932323739e-212 | Forebrain/Midbrain/Hindbra | 90.88 |
| 12 | Nefl | 2.94608397537811 | 0.72 | 0.142 | 2.94345029083083e-190 | 5.87012291500392e-186 | Forebrain/Midbrain/Hindbra | 90.88 |
| 12 | Tubb2b | 3.34689543249946 | 1 | 0.649 | 9.48768464131709e-182 | 1.89212894801787e-177 | Forebrain/Midbrain/Hindbra | 90.88 |
| 12 | Sst | 3.0200197389121 | 0.264 | 0.014 | 3.06157064012199e-178 | 6.10569032759528e-174 | Forebrain/Midbrain/Hindbra | 90.88 |
| 13 | Fabp7 | 4.04612033643945 | 0.997 | 0.149 | 0 | 0 | Forebrain/Midbrain/Hindbra | 63.23 |
| 13 | Sox2 | 2.320468484657 | 0.983 | 0.075 | 0 | 0 | Forebrain/Midbrain/Hindbra | 63.23 |
| 13 | Pantr1 | 2.29968464137079 | 0.938 | 0.142 | 0 | 0 | Forebrain/Midbrain/Hindbra | 63.23 |
| 13 | Hes5 | 2.29306009759037 | 0.746 | 0.043 | 0 | 0 | Forebrain/Midbrain/Hindbra | 63.23 |
| 13 | Rfx4 | 1.67572654716034 | 0.869 | 0.04 | 0 | 0 | Forebrain/Midbrain/Hindbra | 63.23 |
| 13 | Pax3 | 1.64405061436472 | 0.825 | 0.061 | 0 | 0 | Forebrain/Midbrain/Hindbra | 63.23 |
| 13 | Msx3 | 1.5725481210646 | 0.684 | 0.047 | 0 | 0 | Forebrain/Midbrain/Hindbra | 63.23 |
| 13 | Fez1 | 1.43761174061296 | 0.821 | 0.147 | 6.27112146869194e-220 | 1.25064975450123e-215 | Forebrain/Midbrain/Hindbra | 63.23 |
| 13 | Ezr | 1.58268529774223 | 0.918 | 0.234 | 4.86266334420435e-208 | 9.69760950734673e-204 | Forebrain/Midbrain/Hindbra | 63.23 |
| 13 | Ckb | 2.44208498109252 | 0.99 | 0.606 | 2.77892338259236e-152 | 5.54200690190394e-148 | Forebrain/Midbrain/Hindbra | 63.23 |
| 13 | Metrn | 1.4972287944492 | 0.866 | 0.324 | 4.0770621439914e-142 | 8.13088503376205e-138 | Forebrain/Midbrain/Hindbra | 63.23 |
| 13 | Phgdh | 1.56390789208974 | 0.945 | 0.441 | 8.84456361222072e-133 | 1.76387132118518e-128 | Forebrain/Midbrain/Hindbra | 63.23 |
| 13 | Tubb2b | 1.37694191244972 | 1 | 0.65 | 4.08402045365712e-120 | 8.1447619907284e-116 | Forebrain/Midbrain/Hindbra | 63.23 |
| 13 | Dbi | 1.86282535321749 | 1 | 0.849 | 3.52652485910952e-114 | 7.03294852652212e-110 | Forebrain/Midbrain/Hindbra | 63.23 |
| 13 | Hmgb3 | 1.52659671579267 | 0.986 | 0.755 | 1.40226547757583e-113 | 2.79653804192948e-109 | Forebrain/Midbrain/Hindbra | 63.23 |
| 13 | Igfbp2 | 1.4505384945042 | 0.962 | 0.434 | 2.55130110775763e-111 | 5.08805979920105e-107 | Forebrain/Midbrain/Hindbra | 63.23 |
| 13 | Ccnd1 | 1.79565918135515 | 0.749 | 0.321 | 1.61446107777268e-83 | 3.21971972740205e-79 | Forebrain/Midbrain/Hindbra | 63.23 |
| 13 | Hmgb2 | 1.57672141901156 | 0.924 | 0.772 | 5.40175424550117e-50 | 1.0772718491803e-45 | Forebrain/Midbrain/Hindbra | 63.23 |
| 13 | Cdca8 | 1.43191372700677 | 0.601 | 0.284 | 2.17594840100645e-45 | 4.33949389612717e-41 | Forebrain/Midbrain/Hindbra | 63.23 |
| 13 | Cenpf | 1.42145199605005 | 0.509 | 0.224 | 2.0828181157746e-34 | 4.15376416828928e-30 | Forebrain/Midbrain/Hindbra | 63.23 |
| 14 | Mki67 | 1.71737699790102 | 0.91 | 0.196 | 1.0589825574245e-197 | 2.11192891427168e-193 | Paraxial mesoderm | 91.4 |
| 14 | Cenpf | 1.94250349863075 | 0.928 | 0.209 | 1.47444499013778e-193 | 2.94048564383177e-189 | Paraxial mesoderm | 91.4 |
| 14 | Pclaf | 2.06548107918052 | 0.993 | 0.297 | 1.75175676215577e-188 | 3.49352851076726e-184 | Paraxial mesoderm | 91.4 |
| 14 | Tpx2 | 1.76686122325701 | 0.932 | 0.237 | 7.05266420859058e-186 | 1.40651282311922e-181 | Paraxial mesoderm | 91.4 |
| 14 | Birc5 | 1.7377303835296 | 0.964 | 0.269 | 4.79635572100386e-177 | 9.565372214398e-173 | Paraxial mesoderm | 91.4 |
| 14 | Top2a | 2.03750973050173 | 0.932 | 0.245 | 2.75101273348827e-174 | 5.48634469439566e-170 | Paraxial mesoderm | 91.4 |
| 14 | Cdca8 | 1.6071559947671 | 0.964 | 0.271 | 4.55080079431583e-173 | 9.07566202410407e-169 | Paraxial mesoderm | 91.4 |
| 14 | Cenpe | 1.49666792494701 | 0.821 | 0.181 | 7.63959300535261e-170 | 1.52356403305747e-165 | Paraxial mesoderm | 91.4 |
| 14 | Nusap1 | 1.46948968702249 | 0.763 | 0.151 | 7.9847321322887e-162 | 1.59239512914234e-157 | Paraxial mesoderm | 91.4 |
| 14 | Ube2c | 1.87323968408397 | 0.885 | 0.282 | 2.6570010368364e-134 | 5.29885716776283e-130 | Paraxial mesoderm | 91.4 |
| 14 | Hmgb2 | 2.04116197292207 | 1 | 0.77 | 7.14019969714992e-134 | 1.42397002560261e-129 | Paraxial mesoderm | 91.4 |
| 14 | H2afx | 1.91451030779969 | 0.996 | 0.612 | 1.93174314531298e-133 | 3.85247535469767e-129 | Paraxial mesoderm | 91.4 |
| 14 | Cks2 | 1.62052264244681 | 0.971 | 0.475 | 1.44409849183053e-125 | 2.87996562225763e-121 | Paraxial mesoderm | 91.4 |
| 14 | Cenpa | 1.61655621022953 | 0.849 | 0.281 | 1.09795736549513e-121 | 2.18965637400693e-117 | Paraxial mesoderm | 91.4 |

**<u>Supplementary Table S1:</u> Differential gene expression analysis at day 11**

| Cluster | Gene | avg_logFC | pct.1 | pct.2 | p_val | p_val_adj | most expressed celltype | pct.celltype |
|---|---|---|---|---|---|---|---|---|
| 14 | Hist1h1b | 2.65921318284773 | 0.875 | 0.342 | 3.0470058143411e-118 | 6.07664369554046e-114 | Paraxial mesoderm | 91.4 |
| 14 | Hist1h1a | 1.46174031206909 | 0.677 | 0.176 | 2.58340946410387e-116 | 5.15209349426235e-112 | Paraxial mesoderm | 91.4 |
| 14 | Prrx1 | 1.81084959149235 | 0.889 | 0.41 | 2.27312312722265e-110 | 4.53328945262012e-106 | Paraxial mesoderm | 91.4 |
| 14 | Tubb4b | 1.45071938676181 | 0.957 | 0.65 | 5.67608596547373e-79 | 1.13198182409443e-74 | Paraxial mesoderm | 91.4 |
| 14 | Hist1h4d | 1.47793114216139 | 0.821 | 0.486 | 1.29560686718141e-67 | 2.58382877521988e-63 | Paraxial mesoderm | 91.4 |
| 14 | Hist1h1e | 1.67222186872334 | 0.921 | 0.687 | 5.19809919710171e-67 | 1.03665692287799e-62 | Paraxial mesoderm | 91.4 |
| 15 | Tlx1 | 1.58114500319523 | 0.735 | 0.068 | 0 | 0 | Paraxial mesoderm | 93.77 |
| 15 | Kazald1 | 1.53437609290485 | 0.669 | 0.069 | 4.91013115433537e-265 | 9.79227456109103e-261 | Paraxial mesoderm | 93.77 |
| 15 | Slc40a1 | 1.30676893555575 | 0.603 | 0.107 | 1.26476606291379e-146 | 2.52232295926898e-142 | Paraxial mesoderm | 93.77 |
| 15 | Tac2 | 2.05211604695107 | 0.693 | 0.231 | 4.05980742640355e-83 | 8.0964739504766e-79 | Paraxial mesoderm | 93.77 |
| 15 | Tshz2 | 1.16249262463772 | 0.969 | 0.685 | 3.47772182984785e-78 | 6.93562064526557e-74 | Paraxial mesoderm | 93.77 |
| 15 | Sparcl1 | 1.35807454901958 | 0.938 | 0.531 | 9.50716689967431e-76 | 1.89601429480205e-71 | Paraxial mesoderm | 93.77 |
| 15 | Ntrk2 | 1.12073501660484 | 0.642 | 0.21 | 2.73671985416191e-73 | 5.45784040515509e-69 | Paraxial mesoderm | 93.77 |
| 15 | Tgfbi | 1.22622656082169 | 0.716 | 0.333 | 3.59377969751223e-60 | 7.16707485074863e-56 | Paraxial mesoderm | 93.77 |
| 15 | Ppp1r14a | 0.973240467373996 | 0.712 | 0.309 | 1.7853181887218e-59 | 3.56046006376788e-55 | Paraxial mesoderm | 93.77 |
| 15 | Akap12 | 1.08341162275599 | 0.852 | 0.483 | 1.1054316632903e-56 | 2.20456236609985e-52 | Paraxial mesoderm | 93.77 |
| 15 | Tcf21 | 0.853897166986069 | 0.588 | 0.209 | 1.06990943274893e-54 | 2.13372038173118e-50 | Paraxial mesoderm | 93.77 |
| 15 | Hmga2 | 1.1844888396227 | 0.887 | 0.604 | 1.07520469974849e-50 | 2.14428073270842e-46 | Paraxial mesoderm | 93.77 |
| 15 | Kcnj8 | 0.949732479008597 | 0.498 | 0.171 | 9.98579896341058e-50 | 1.99146788727297e-45 | Paraxial mesoderm | 93.77 |
| 15 | Cdh11 | 0.862574923852912 | 0.844 | 0.527 | 5.15436805629212e-48 | 1.02793562146634e-43 | Paraxial mesoderm | 93.77 |
| 15 | Gm49708 | 1.12709556218954 | 0.763 | 0.466 | 4.10692204432532e-47 | 8.19043463299799e-43 | Paraxial mesoderm | 93.77 |
| 15 | Mfap2 | 0.805834848476802 | 0.973 | 0.689 | 2.83145438158886e-45 | 5.64676947320267e-41 | Paraxial mesoderm | 93.77 |
| 15 | Dnm3os | 0.837582129914915 | 0.844 | 0.514 | 1.88491410898155e-40 | 3.75908420754191e-36 | Paraxial mesoderm | 93.77 |
| 15 | Fos | 1.19204631608758 | 0.821 | 0.54 | 3.25538608829255e-38 | 6.49221647588183e-34 | Paraxial mesoderm | 93.77 |
| 15 | Ier2 | 0.86705808583072 | 0.949 | 0.792 | 8.27947473588801e-36 | 1.65117564657815e-31 | Paraxial mesoderm | 93.77 |
| 15 | Jak1 | 0.806342924862596 | 0.786 | 0.591 | 6.95876799616049e-34 | 1.38778710147429e-29 | Paraxial mesoderm | 93.77 |
| 16 | Car3 | 1.54202242904238 | 0.49 | 0.021 | 0 | 0 | Paraxial mesoderm | 86.83 |
| 16 | Msc | 1.41847740568688 | 0.671 | 0.02 | 0 | 0 | Paraxial mesoderm | 86.83 |
| 16 | Itga4 | 1.21485334610529 | 0.704 | 0.057 | 0 | 0 | Paraxial mesoderm | 86.83 |
| 16 | Myog | 1.07054898873467 | 0.337 | 0.016 | 4.56062749403351e-211 | 9.09525941135103e-207 | Paraxial mesoderm | 86.83 |
| 16 | Pdgfa | 1.30422982729989 | 0.79 | 0.176 | 7.93742241858623e-156 | 1.58296015293865e-151 | Paraxial mesoderm | 86.83 |
| 16 | Tnnt1 | 1.71831785644439 | 0.811 | 0.261 | 7.12714819431323e-124 | 1.42136716439189e-119 | Paraxial mesoderm | 86.83 |
| 16 | Ppp1r14b | 1.50300419754447 | 1 | 0.96 | 9.83752099807932e-116 | 1.96189681264696e-111 | Paraxial mesoderm | 86.83 |
| 16 | Hspe1 | 1.00985084716045 | 0.996 | 0.983 | 6.84439862228908e-97 | 1.36497841724311e-92 | Paraxial mesoderm | 86.83 |
| 16 | Hspd1 | 1.27613177787966 | 1 | 0.948 | 7.37516239844265e-96 | 1.47082863712142e-91 | Paraxial mesoderm | 86.83 |
| 16 | Anp32b | 0.954277940017528 | 0.996 | 0.97 | 5.30751347588341e-82 | 1.05847741249543e-77 | Paraxial mesoderm | 86.83 |
| 16 | Ran | 0.984299811295397 | 1 | 0.97 | 5.74022127998471e-81 | 1.14477232986735e-76 | Paraxial mesoderm | 86.83 |
| 16 | Dctpp1 | 1.09599061206594 | 0.967 | 0.726 | 7.91349148046048e-81 | 1.57818760594823e-76 | Paraxial mesoderm | 86.83 |
| 16 | Ncl | 1.09095543242307 | 1 | 0.982 | 1.1862256731534e-78 | 2.36568985996982e-74 | Paraxial mesoderm | 86.83 |
| 16 | H2afz | 1.09279633033463 | 1 | 0.991 | 1.38500998133307e-73 | 2.76212540577253e-69 | Paraxial mesoderm | 86.83 |
| 16 | Capn6 | 1.07323996101786 | 0.802 | 0.326 | 3.86506507204279e-73 | 7.70809927317493e-69 | Paraxial mesoderm | 86.83 |
| 16 | Ranbp1 | 0.970497914185662 | 0.988 | 0.901 | 6.89646787074073e-69 | 1.37536258746182e-64 | Paraxial mesoderm | 86.83 |
| 16 | Ybx3 | 1.02490690589749 | 0.947 | 0.727 | 1.84025452524814e-68 | 3.67001959970236e-64 | Paraxial mesoderm | 86.83 |
| 16 | S100a10 | 1.25190228302706 | 0.782 | 0.364 | 1.49196810735902e-57 | 2.97543199650609e-53 | Paraxial mesoderm | 86.83 |
| 16 | Nrk | 1.10389717241644 | 0.831 | 0.437 | 8.61067985416106e-53 | 1.71722788331534e-48 | Paraxial mesoderm | 86.83 |
| 16 | Itm2a | 1.11671145918834 | 0.942 | 0.761 | 4.97573156403442e-50 | 9.92310145815384e-46 | Paraxial mesoderm | 86.83 |
| 17 | Hes5 | 2.4293212347557 | 0.919 | 0.048 | 0 | 0 | Forebrain/Midbrain/Hindbra | 55.68 |
| 17 | Sox2 | 2.22300224912322 | 0.978 | 0.087 | 0 | 0 | Forebrain/Midbrain/Hindbra | 55.68 |
| 17 | Pax3 | 1.97119611165647 | 0.951 | 0.068 | 0 | 0 | Forebrain/Midbrain/Hindbra | 55.68 |
| 17 | Hist1h3c | 2.01009639123787 | 0.984 | 0.132 | 1.70065892873412e-256 | 3.39162410157446e-252 | Forebrain/Midbrain/Hindbra | 55.68 |
| 17 | Fabp7 | 3.45445041328335 | 0.995 | 0.16 | 7.61798426698961e-238 | 1.51925460236574e-233 | Forebrain/Midbrain/Hindbra | 55.68 |
| 17 | Pantr1 | 2.26682541686 | 0.989 | 0.152 | 3.11396646205415e-231 | 6.21018331527458e-227 | Forebrain/Midbrain/Hindbra | 55.68 |
| 17 | Prc1 | 2.27426790884898 | 0.951 | 0.167 | 1.55364220551203e-191 | 3.09842865045264e-187 | Forebrain/Midbrain/Hindbra | 55.68 |
| 17 | Rrm2 | 2.05697230905716 | 0.968 | 0.215 | 2.72647760822397e-173 | 5.43741429408107e-169 | Forebrain/Midbrain/Hindbra | 55.68 |
| 17 | Cenpf | 2.29195033925449 | 0.973 | 0.217 | 3.63770900174471e-155 | 7.25468306217948e-151 | Forebrain/Midbrain/Hindbra | 55.68 |
| 17 | Top2a | 2.1244833110862 | 0.995 | 0.252 | 5.0796774296809e-145 | 1.01304006980126e-140 | Forebrain/Midbrain/Hindbra | 55.68 |
| 17 | Hist1h1b | 3.2806617655197 | 1 | 0.345 | 1.95142237866738e-139 | 3.89172164977636e-135 | Forebrain/Midbrain/Hindbra | 55.68 |
| 17 | Pclaf | 2.17906374672465 | 1 | 0.305 | 2.10945533509297e-135 | 4.20688677477592e-131 | Forebrain/Midbrain/Hindbra | 55.68 |
| 17 | Cdca8 | 2.00943967572218 | 0.957 | 0.28 | 4.87247411585189e-129 | 9.71717512924343e-125 | Forebrain/Midbrain/Hindbra | 55.68 |
| 17 | Hist1h4d | 2.12514432551521 | 0.995 | 0.486 | 5.24076578043619e-114 | 1.04516591959239e-109 | Forebrain/Midbrain/Hindbra | 55.68 |
| 17 | Dek | 2.04391739912215 | 1 | 0.754 | 2.32973693372003e-107 | 4.64619436691786e-103 | Forebrain/Midbrain/Hindbra | 55.68 |
| 17 | Hist1h1e | 2.32079169304474 | 1 | 0.688 | 8.08637994352414e-104 | 1.61266675213702e-99 | Forebrain/Midbrain/Hindbra | 55.68 |
| 17 | Ccnd1 | 2.10477989332379 | 0.924 | 0.323 | 8.98090671371577e-102 | 1.79106222591634e-97 | Forebrain/Midbrain/Hindbra | 55.68 |
| 17 | Hmgb2 | 2.39410857768192 | 1 | 0.772 | 3.21387027258803e-101 | 6.4094214846223e-97 | Forebrain/Midbrain/Hindbra | 55.68 |
| 17 | Ckb | 2.16006167201731 | 0.995 | 0.611 | 3.13964077196214e-95 | 6.2613855915241e-91 | Forebrain/Midbrain/Hindbra | 55.68 |
| 17 | Ube2c | 1.95123492075096 | 0.843 | 0.29 | 5.63270272012842e-82 | 1.12332990347521e-77 | Forebrain/Midbrain/Hindbra | 55.68 |
| 18 | Tnnc2 | 4.920172199468 | 0.716 | 0.008 | 0 | 0 | Cardiomyocytes | 85.07 |
| 18 | Acta1 | 4.57498477156116 | 0.672 | 0.014 | 0 | 0 | Cardiomyocytes | 85.07 |
| 18 | Myog | 4.21036821510322 | 1 | 0.018 | 0 | 0 | Cardiomyocytes | 85.07 |
| 18 | Myh3 | 3.8193921604858 | 0.925 | 0.004 | 0 | 0 | Cardiomyocytes | 85.07 |
| 18 | Neb | 3.6977693334292 | 0.97 | 0.019 | 0 | 0 | Cardiomyocytes | 85.07 |
| 18 | Il17b | 2.88291837342083 | 0.821 | 0.005 | 0 | 0 | Cardiomyocytes | 85.07 |
| 18 | Mybph | 2.77111494503765 | 0.672 | 0.005 | 0 | 0 | Cardiomyocytes | 85.07 |
| 18 | Mymx | 2.62449191633778 | 0.866 | 0.007 | 0 | 0 | Cardiomyocytes | 85.07 |
| 18 | Myod1 | 2.57062220503359 | 0.97 | 0.017 | 0 | 0 | Cardiomyocytes | 85.07 |
| 18 | Tnnt3 | 2.94324463932628 | 0.687 | 0.018 | 6.67361750662993e-283 | 1.33091953934721e-278 | Cardiomyocytes | 85.07 |
| 18 | Myl1 | 4.57942317032187 | 0.985 | 0.076 | 1.52909084963995e-178 | 3.04946588143694e-174 | Cardiomyocytes | 85.07 |
| 18 | Mylpf | 6.21916241368571 | 0.985 | 0.086 | 9.84135435420213e-166 | 1.96266129885853e-161 | Cardiomyocytes | 85.07 |
| 18 | Sln | 2.98019430868111 | 0.731 | 0.046 | 1.79086372077764e-147 | 3.57151951834685e-143 | Cardiomyocytes | 85.07 |
| 18 | Tnnt1 | 4.50602596149168 | 1 | 0.271 | 2.59453555624994e-71 | 5.17428225982925e-67 | Cardiomyocytes | 85.07 |
| 18 | Cdkn1a | 3.22821490804274 | 0.985 | 0.273 | 6.55802018585276e-63 | 1.30786596566462e-58 | Cardiomyocytes | 85.07 |
| 18 | Nes | 2.62649119603937 | 0.985 | 0.333 | 3.00644941366596e-56 | 5.99576206567402e-52 | Cardiomyocytes | 85.07 |
| 18 | Acta2 | 2.68799938098343 | 1 | 0.331 | 1.53060120003444e-51 | 3.05247797322868e-47 | Cardiomyocytes | 85.07 |
| 18 | Tpm2 | 4.53448607771586 | 1 | 0.548 | 2.28642122531781e-49 | 4.55980984965131e-45 | Cardiomyocytes | 85.07 |
| 18 | Actc1 | 2.82583231144205 | 1 | 0.42 | 6.7264672834626e-43 | 1.34145937034095e-38 | Cardiomyocytes | 85.07 |
| 18 | Tnnc1 | 2.71349577084873 | 0.821 | 0.308 | 2.4884134699939e-29 | 4.96264298320884e-25 | Cardiomyocytes | 85.07 |
| 19 | Fcer1g | 4.86246971436003 | 0.964 | 0.011 | 0 | 0 | Blood progenitors 1 | 98.21 |
| 19 | C1qb | 4.86009928098934 | 0.929 | 0.008 | 0 | 0 | Blood progenitors 1 | 98.21 |
| 19 | Cd52 | 4.75814626101734 | 0.982 | 0.008 | 0 | 0 | Blood progenitors 1 | 98.21 |
| 19 | Tyrobp | 4.61850584070407 | 0.964 | 0.006 | 0 | 0 | Blood progenitors 1 | 98.21 |
| 19 | C1qc | 4.10185214030983 | 0.911 | 0.004 | 0 | 0 | Blood progenitors 1 | 98.21 |
| 19 | C1qa | 3.82551764546773 | 0.857 | 0.002 | 0 | 0 | Blood progenitors 1 | 98.21 |
| 19 | Rac2 | 3.53117872396329 | 0.982 | 0.005 | 0 | 0 | Blood progenitors 1 | 98.21 |
| 19 | Lst1 | 3.4112322284918 | 0.929 | 0.017 | 0 | 0 | Blood progenitors 1 | 98.21 |
| 19 | Lyz2 | 3.20000589702903 | 0.857 | 0.01 | 0 | 0 | Blood progenitors 1 | 98.21 |
| 19 | Aif1 | 3.09001814995431 | 0.964 | 0.004 | 0 | 0 | Blood progenitors 1 | 98.21 |
| 19 | Fyb | 3.03502182331594 | 0.964 | 0.014 | 0 | 0 | Blood progenitors 1 | 98.21 |
| 19 | Arhgdib | 2.80683394630845 | 0.964 | 0.244 | 1.8043332173379e-57 | 3.59838173533698e-53 | Blood progenitors 1 | 98.21 |
| 19 | Tmsb4x | 3.49216403477365 | 1 | 0.985 | 8.08283320595124e-38 | 1.61195942626286e-33 | Blood progenitors 1 | 98.21 |
| 19 | Fth1 | 3.64357307788043 | 1 | 0.999 | 1.92665452563276e-37 | 3.84232712046941e-33 | Blood progenitors 1 | 98.21 |
| 19 | Cyba | 2.95185991362191 | 0.946 | 0.549 | 2.50356417256837e-34 | 4.9928580293531e-30 | Blood progenitors 1 | 98.21 |
| 19 | Plin2 | 2.90624392960845 | 0.964 | 0.54 | 1.65680521866076e-31 | 3.30416664757516e-27 | Blood progenitors 1 | 98.21 |
| 19 | Selenop | 2.97178546255373 | 0.839 | 0.32 | 7.74991787433488e-31 | 1.5455661216786e-26 | Blood progenitors 1 | 98.21 |
| 19 | Ctsd | 3.32260157908163 | 0.946 | 0.591 | 6.19193079171008e-28 | 1.23485675779074e-23 | Blood progenitors 1 | 98.21 |
| 19 | Ctsb | 3.60696329125924 | 0.929 | 0.629 | 2.88787181665803e-27 | 5.7592827639611e-23 | Blood progenitors 1 | 98.21 |
| 19 | Apoe | 4.54082087607736 | 0.839 | 0.505 | 5.85815522890415e-21 | 1.16829189730035e-16 | Blood progenitors 1 | 98.21 |
| 20 | Afp | 5.59357457315371 | 0.824 | 0.01 | 0 | 0 | Gut | 61.76 |
| 20 | Apoa2 | 5.07404776856709 | 0.559 | 0.005 | 0 | 0 | Gut | 61.76 |
| 20 | Alb | 3.81446174939934 | 0.5 | 0.002 | 0 | 0 | Gut | 61.76 |
| 20 | Apoa1 | 3.56093287295114 | 0.765 | 0.007 | 0 | 0 | Gut | 61.76 |
| 20 | Apoc2 | 3.2260021989082 | 0.5 | 0.003 | 0 | 0 | Gut | 61.76 |
| 20 | Ambp | 2.960641729961 | 0.794 | 0.002 | 0 | 0 | Gut | 61.76 |
| 20 | Epcam | 2.7938058024085 | 0.912 | 0.016 | 0 | 0 | Gut | 61.76 |
| 20 | Apom | 2.71508992641866 | 0.529 | 0.002 | 0 | 0 | Gut | 61.76 |
| 20 | Fgb | 2.61804924702685 | 0.647 | 0.001 | 0 | 0 | Gut | 61.76 |
| 20 | Serpina6 | 2.59272959935098 | 0.382 | 0.001 | 0 | 0 | Gut | 61.76 |
| 20 | Spp2 | 2.44400399534659 | 0.647 | 0.001 | 0 | 0 | Gut | 61.76 |
| 20 | Krt8 | 3.06320198319738 | 0.941 | 0.024 | 1.8214963769307e-243 | 3.6326102245129e-239 | Gut | 61.76 |
| 20 | Ttr | 5.05375777414103 | 0.941 | 0.025 | 1.96596725562459e-231 | 3.92072849789212e-227 | Gut | 61.76 |
| 20 | Krt18 | 3.76050282594035 | 0.971 | 0.035 | 1.32566380987562e-186 | 2.64377133603494e-182 | Gut | 61.76 |
| 20 | Rbp4 | 5.19145047007387 | 0.941 | 0.044 | 1.48355186321373e-144 | 2.95864748080714e-140 | Gut | 61.76 |
| 20 | Trf | 3.78816539822325 | 0.676 | 0.026 | 2.10265083939386e-119 | 4.19331656900318e-115 | Gut | 61.76 |
| 20 | Bex1 | 3.7222152392822 | 1 | 0.403 | 1.12001871905105e-29 | 2.23365333140352e-25 | Gut | 61.76 |
| 20 | Gstm1 | 2.59871428773203 | 0.971 | 0.365 | 7.06673759213656e-27 | 1.40931947799979e-22 | Gut | 61.76 |
| 20 | Apoe | 3.31359653151899 | 0.912 | 0.506 | 4.50961080315348e-16 | 8.99351682472899e-12 | Gut | 61.76 |
| 20 | Mt1 | 3.32660308225839 | 0.824 | 0.349 | 9.13144091878818e-13 | 1.82108326243393e-8 | Gut | 61.76 |

**Supplementary Table S2:** **Differential gene expression analysis (between the cardiomyocytes clusters of day 11)**

| cluster | gene | avg_log2FC | pct.1 | pct.2 | p_val | p_val_adj |
|---|---|---|---|---|---|---|
| 6 | Actg1 | 1.9520497598933 | 1 | 0.98 | 9.3292424993531E-145 | 1.86053083164599E-140 |
| 6 | Actb | 2.13442707317024 | 0.992 | 0.933 | 1.47819468257727E-121 | 2.94796365546385E-117 |
| 6 | Acta2 | 2.68484766092077 | 0.942 | 0.608 | 6.15351234694026E-117 | 1.2271949673503E-112 |
| 6 | Cald1 | 1.53253259486526 | 0.934 | 0.62 | 2.23908523204642E-115 | 4.46540767827017E-111 |
| 6 | Vsnl1 | 1.52489294166401 | 0.811 | 0.334 | 7.30934997371835E-97 | 1.45770366525865E-92 |
| 6 | Gucy1a1 | 1.29918997463378 | 0.886 | 0.637 | 2.80083956814816E-80 | 5.58571435075788E-76 |
| 6 | Tpm2 | 1.257035819452 | 0.916 | 0.701 | 4.59570578360096E-74 | 9.16521604423539E-70 |
| 6 | Shox2 | 1.19964209190323 | 0.843 | 0.415 | 1.98337277675618E-66 | 3.95544032868485E-62 |
| 6 | Ptn | 2.23634395737377 | 0.811 | 0.513 | 2.86394482682357E-66 | 5.71156516813425E-62 |
| 6 | Maged2 | 1.18372950759593 | 0.962 | 0.851 | 6.3454307639539E-62 | 1.26546925725533E-57 |
| 6 | Meg3 | 1.50051497903603 | 0.871 | 0.595 | 9.24182981137988E-62 | 1.84309811928349E-57 |
| 6 | Mdk | 1.10488062248775 | 0.95 | 0.805 | 4.01236480067597E-55 | 8.00185912198808E-51 |
| 6 | Cdkn1c | 1.57559149761615 | 0.958 | 0.906 | 4.07081202660618E-55 | 8.11842042466071E-51 |
| 6 | Prnp | 1.07638223833597 | 0.861 | 0.756 | 5.77542993384708E-50 | 1.15179399170712E-45 |
| 6 | Tagln | 1.79456404971038 | 0.357 | 0.071 | 1.70746117533773E-45 | 3.40518982197603E-41 |
| 6 | Pnmt | 1.36580058833633 | 0.536 | 0.225 | 4.11175508636781E-39 | 8.20007316874332E-35 |
| 6 | Csrp2 | 1.25999558667678 | 0.97 | 0.964 | 4.18961116963935E-29 | 8.35534155561176E-25 |
| 6 | Mgp | 1.11264458561913 | 0.516 | 0.308 | 3.78991869328861E-24 | 7.55823485002547E-20 |
| 6 | Gal | 1.32615716611783 | 0.219 | 0.105 | 0.000000000758751728 | 0.00001513178571312 |
| 6 | Fabp5 | 1.26011794397178 | 0.882 | 0.853 | 0.000000011982820195 | 0.000238973383146723 |
| 8 | Myl2 | 3.91924440186805 | 0.952 | 0.346 | 5.27177377234951E-167 | 1.05134984341966E-162 |
| 8 | Myl3 | 2.87809664062242 | 0.984 | 0.559 | 4.41193166838023E-155 | 8.79871532625069E-151 |
| 8 | Myh7 | 3.01672448698971 | 0.982 | 0.669 | 1.28109789427715E-145 | 2.55489353055692E-141 |
| 8 | Mpped2 | 1.34135916286582 | 0.84 | 0.197 | 2.77773947114668E-134 | 5.53964582730782E-130 |
| 8 | Pln | 2.13067249544387 | 0.936 | 0.494 | 3.59481088311273E-112 | 7.16913134419172E-108 |
| 8 | Hopx | 1.18782692538119 | 0.884 | 0.427 | 9.23373034467432E-88 | 1.8414828426384E-83 |
| 8 | Retreg1 | 0.603194149438958 | 0.573 | 0.095 | 2.72638502325051E-87 | 5.43722965186849E-83 |
| 8 | Mhrt | 0.724323835327955 | 0.557 | 0.107 | 1.91874214902498E-76 | 3.82654746780053E-72 |
| 8 | Spink4 | 1.27694594691667 | 0.671 | 0.24 | 4.4274041281581E-67 | 8.82957205278569E-63 |
| 8 | Mb | 0.939573463172259 | 0.605 | 0.234 | 4.19177329513539E-54 | 8.3596534824885E-50 |
| 8 | Crip2 | 0.701507105371309 | 0.991 | 0.941 | 5.40765839945553E-51 | 1.07844931460342E-46 |
| 8 | Actn2 | 0.708032558494068 | 0.943 | 0.699 | 1.02100697320104E-46 | 2.03619420665484E-42 |
| 8 | Fhl2 | 0.959723969666818 | 0.842 | 0.636 | 8.2634084299767E-46 | 1.64797154319025E-41 |
| 8 | Mllt3 | 0.717005407346828 | 0.776 | 0.485 | 7.76876246428685E-43 | 1.54932429825273E-38 |
| 8 | Cyba | 0.604797659948406 | 0.9 | 0.712 | 1.37263368048981E-41 | 2.73744334900083E-37 |
| 8 | Gyg | 0.619106891466274 | 0.998 | 0.987 | 3.83198467925149E-41 | 7.64212704583124E-37 |
| 8 | Kcne1 | 0.648115701807294 | 0.5 | 0.181 | 7.29894536982245E-39 | 1.45562867510369E-34 |
| 8 | Cox8b | 0.614498061937409 | 0.79 | 0.508 | 3.59412544647057E-33 | 7.16776437789626E-29 |
| 8 | 2410006H16Rik | 0.688849147010127 | 0.973 | 0.894 | 6.00894579360899E-33 | 1.19836405961944E-28 |
| 8 | Cited1 | 0.969814821272424 | 0.662 | 0.418 | 6.17370735109916E-25 | 1.23122245702971E-20 |
| 9 | Myh6 | 1.59235974577339 | 1 | 0.862 | 3.97446437709652E-105 | 7.92627430724359E-101 |
| 9 | Casq1 | 1.79759134202071 | 0.883 | 0.494 | 1.91745079450269E-86 | 3.82397211947671E-82 |
| 9 | Myl7 | 0.817380821068884 | 1 | 1 | 1.32413621983951E-67 | 2.64072486322594E-63 |
| 9 | Pam | 1.0724249042968 | 0.984 | 0.876 | 1.22914310728383E-66 | 2.45128009885614E-62 |
| 9 | Myl4 | 0.757030806112411 | 1 | 0.997 | 1.73885433631051E-65 | 3.46779720290405E-61 |
| 9 | Itga6 | 1.09158197561414 | 0.81 | 0.44 | 2.25547832085908E-65 | 4.49810041528927E-61 |
| 9 | Tesc | 0.972035422900798 | 0.892 | 0.623 | 1.7106469861392E-56 | 3.41154328445741E-52 |
| 9 | Stard10 | 0.885684559827296 | 0.874 | 0.564 | 5.29535366765539E-55 | 1.05605238194051E-50 |
| 9 | Palld | 0.912296477734801 | 0.963 | 0.828 | 2.76924818275449E-54 | 5.52271165086727E-50 |
| 9 | Slc8a1 | 0.851869941194078 | 0.991 | 0.927 | 7.73181438459535E-52 | 1.54195574271985E-47 |
| 9 | Smpx | 0.808396923233642 | 0.953 | 0.767 | 8.92677490500744E-47 | 1.78026671930563E-42 |
| 9 | Ccnd2 | 0.877567370057972 | 0.946 | 0.738 | 2.75286870172281E-46 | 5.49004605184581E-42 |
| 9 | Obscn | 0.824131869765032 | 0.93 | 0.694 | 5.16813613304334E-46 | 1.03068138901283E-41 |
| 9 | Mlf1 | 0.833262290040485 | 0.892 | 0.7 | 2.20424632681714E-42 | 4.39592844957143E-38 |
| 9 | Sln | 1.60475410018664 | 0.445 | 0.131 | 1.07163298871321E-40 | 2.13715766939075E-36 |
| 9 | Atcayos | 0.739159367184943 | 0.977 | 0.81 | 1.95143439768714E-36 | 3.89174561930747E-32 |
| 9 | Calca | 0.902898889852385 | 0.614 | 0.254 | 3.74568819625215E-36 | 7.47002596978566E-32 |
| 9 | Nppa | 1.94085644980968 | 0.319 | 0.079 | 1.73789152270502E-31 | 3.46587706373062E-27 |
| 9 | Ankrd1 | 0.904934716761817 | 0.543 | 0.262 | 7.29936675209728E-26 | 1.45571271137076E-21 |
| 9 | Mest | 1.17185018930633 | 0.686 | 0.481 | 1.43322925683164E-23 | 0.0000000000000000002 |

**Supplementary Table S3: Differential gene expression analysis (between the myoblasts clusters of day 11)**

| gene | avg_log2FC | pct.1 | pct.2 | p_val | p_val_adj |
|---|---|---|---|---|---|
| *Myh3* | 3.79102992898606 | 0.925 | 0.029 | 1.80751541028337e-56 | 3.60472798272812e-52 |
| *Mylpf* | 6.1754758583588 | 0.985 | 0.14 | 1.94407330217701e-49 | 3.8770653865316e-45 |
| *Myl1* | 4.58211870440919 | 0.985 | 0.156 | 2.97563722722983e-47 | 5.93431332226446e-43 |
| *Cryab* | 3.62013700948436 | 0.896 | 0.086 | 9.79636086801911e-46 | 1.95368824790905e-41 |
| *Tnni1* | 4.22431378539966 | 0.94 | 0.132 | 3.98684193695189e-45 | 7.95095887486315e-41 |
| *Sln* | 3.26438545845109 | 0.731 | 0.021 | 4.81864586392395e-43 | 9.60982544642354e-39 |
| *Acta1* | 4.6073109457577 | 0.672 | 0.004 | 5.04301114000861e-42 | 1.00572771165192e-37 |
| *Ttn* | 4.33390152089812 | 0.97 | 0.226 | 2.69224982829685e-41 | 5.3691538325724e-37 |
| *Cdkn1a* | 3.47319533545016 | 0.985 | 0.276 | 2.20943539225039e-40 | 4.40627700276496e-36 |
| *Tnnt2* | 4.30705152686729 | 0.985 | 0.325 | 1.26633249091905e-39 | 2.52544688663986e-35 |
| *Tnnc2* | 4.88194860331071 | 0.716 | 0.041 | 1.98647570683138e-38 | 3.96162850213382e-34 |
| *Myog* | 2.99021913209157 | 1 | 0.337 | 7.46873712999516e-37 | 1.48949024583493e-32 |
| *Actc1* | 4.29935629564102 | 1 | 0.65 | 2.61594495273662e-36 | 5.21697901924264e-32 |
| *Tnnt3* | 2.92595185281961 | 0.687 | 0.037 | 3.19062973330052e-36 | 6.36307287712122e-32 |
| *Tpm2* | 3.70901765014235 | 1 | 0.856 | 1.13969515712656e-35 | 2.2728940518575e-31 |
| *Mef2c* | 3.12750534342919 | 0.91 | 0.239 | 4.84096832943235e-35 | 9.65434313938694e-31 |
| *Acta2* | 3.21139834641567 | 1 | 0.551 | 1.94103369500933e-34 | 3.8710034979571e-30 |
| *Tnnc1* | 5.1043129199129 | 0.821 | 0.218 | 7.79119167555789e-30 | 1.55379735585651e-25 |
| *Myl4* | 4.87508192544678 | 0.836 | 0.321 | 2.15336043882606e-26 | 4.29444672315081e-22 |
| *Meg3* | 3.03687972714313 | 0.881 | 0.667 | 4.13770990416055e-21 | 8.25183486186738e-17 |

pct.1 refers to the percentage of cells where the feature is detected in the cluster 18.
pct.2 refers to the percentage of cells where the feature is detected in the cluster 16.

**<u>Supplementary Table S4</u>: qPCR primers**

| gene | forward | reverse | product length |
|---|---|---|---|
| *Isl1* | GCATCATGATGAAGCAGCTC | CATCGATGCTACTTCACTGC | 275 |
| *Mesp1* | GTCTGCAGCGGGGTGTCGTG | CGGCGGCGTCCAGGTTTCTA | 189 |
| *Myl2* | CCCAGATCCAGGAGTTCAAG | CTGGTCGATCTCCTCTTTGG | 341 |
| *Myl7* | ATCCTGAGTGCCTTCCGCATG | GGTGTCAGCGCAAACACTTGC | 134 |
| *TBP* | CCCCACAACTCTTCCATTCT | GCAGGAGTGATAGGGGTCAT | 103 |
| *Tbx1* | CGAGATGATCGTCACCAAGG | CCAGGAGGAGCTATGGAAAG | 154 |
| *Tcf21* | CTGTAGTTCCACACAAGCGG | CGGTTACATTCACCCAGTCA | 107 |
| *Myf5* | GACAGGGCTGTTACATTCAGG | TGAGGGAACAGGTGGAGAAC | 110 |
| *Tnnt2* | CAAGGAGCTGTGGCAGAGTA | TTCTGGTTGTCATTGATCCG | 120 |
| *MyoD* | GTCGTAGCCATTCTGCCG | AGCACTACAGTGGCGACTCA | 110 |
| *Ebf3* | AGAGCCGAACAACGAGAAAA | GCACATCTCCGGATTCTTGT | 163 |
